# Discovery and characterization of cell-permeable inhibitors of *Leishmania mexicana* CLK1 using an in-cell target engagement assay

**DOI:** 10.1101/2025.06.17.659316

**Authors:** Carolina M. C. Catta-Preta, Priscila Zonzini Ramos, Juliana B. T. Carnielli, Stanley N. S. Vasconcelos, Adam Dowle, Rebeka C. Fanti, Caio Vinicius dos Reis, Adriano Cappellazzo Coelho, Katlin B. Massirer, Jeremy C. Mottram, Rafael M. Couñago

**Affiliations:** Centro de Química Medicinal (CQMED), Centro de Biologia Molecular e Engenharia Genética (CBMEG), Universidade Estadual de Campinas, UNICAMP, 13083-886-Campinas, SP, Brazil; York Biomedical Research Institute, Department of Biology, University of York, York YO10 5DD, United Kingdom; Bioscience Technology Facility, Department of Biology, University of York, York YO10 5DD, United Kingdom; Departamento de Biologia Animal, Instituto de Biologia, Universidade Estadual de Campinas (UNICAMP), Campinas 13083-862, Brazil; Structural Genomics Consortium and Division of Chemical Biology and Medicinal Chemistry, UNC Eshelman School of Pharmacy, University of North Carolina, Chapel Hill, North Carolina 27599, USA

**Keywords:** Protein kinase, infectious diseases, target engagement, leishmaniasis, kinase inhibitor, drug discovery

## Abstract

Leishmaniasis is a neglected tropical disease with limited treatment options and significant unmet medical need. Here, we report the development of a bioluminescence resonance energy transfer (BRET)-based target engagement assay in live intact cells to identify and validate cell-permeable, ATP-competitive inhibitors of *Leishmania mexicana* (Lmx)CLK1. LmxCLK2, a closely related paralogue with an identical protein kinase domain, is also considered in our analysis. Genetic and pharmacological evidence indicates that simultaneous deletion or inhibition of both LmxCLK1/2 is lethal to the parasite. Using our newly developed assay, we screened a library of human kinase inhibitors and identified WZ8040, a third-generation EGFR inhibitor, as a potent LmxCLK1 ligand. WZ8040 demonstrated robust target engagement in both promastigotes and macrophage-internalized amastigotes, with low micromolar EC₅₀ values for parasite killing and minimal toxicity to host macrophages. Biochemical assays confirmed that WZ8040 covalently binds and inhibits LmxCLK1, with mass spectrometry identifying Cys172 as the primary site of modification. Genetic validation using overexpression and knockout lines supports LmxCLK1 as the primary target of WZ8040. However, the retained activity of WZ8040 in mutant lines with the Cys172Ala substitution suggests that covalent binding is not essential for compound efficacy. Our findings highlight the utility of BRET-based assays for target validation in kinetoplastid parasites and underscore the potential of CLK1 as a druggable kinase in *Leishmania*. This integrated approach provides a framework for accelerating the discovery of novel anti-leishmanial agents through target engagement-guided strategies.

**Highlights:**

- A live-cell NanoBRET assay enables *in situ* target engagement of CLK1 in *Leishmania mexicana*.
- The covalent inhibitor WZ8040 effectively targets LmxCLK1 in both promastigote and intracellular amastigote stages.
- CLK1 inhibition by WZ8040 does not depend on covalent modification of a single cysteine residue.
- Combined target engagement and phenotypic data support LmxCLK1 as a druggable kinase in *Leishmania*.

---

Leishmaniasis represents a group of neglected tropical diseases (NTDs) that disproportionately impact impoverished populations, with over one billion individuals at risk and an estimated 700,000 to one million new cases reported annually. Visceral leishmaniasis, the most severe manifestation, carries an untreated mortality rate approaching 90%.^1,2^ Despite its substantial global health burden, leishmaniasis remains critically underfunded in research and development, largely due to its prevalence in resource-limited settings.

Current therapeutic options include pentavalent antimonials such as sodium stibogluconate and meglumine antimoniate, as well as amphotericin B, miltefosine, and paromomycin. While these agents exhibit varying degrees of efficacy, their clinical utility is often constrained by significant drawbacks, including severe toxicity, the emergence of drug resistance, high treatment costs, and logistical challenges in administration—particularly in low-resource environments. For instance, amphotericin B necessitates hospitalization due to its intravenous administration and nephrotoxic profile, whereas miltefosine, the only oral drug option, is associated with potential teratogenicity. These limitations underscore the urgent need for the development of novel, safer, and more accessible therapeutic strategies.^3–5^

The discovery of anti-leishmanial drugs presents inherent challenges, largely due to the complex biology of the parasite and its intracellular niche. Leishmaniasis is caused by protozoan parasites of the genus *Leishmania*, which infect mammalian macrophages and replicate within acidic parasitophorous vacuoles.^5^ To effectively target the intracellular amastigote form, therapeutic compounds must overcome multiple biological barriers and retain their efficacy after crossing several cellular membranes and compartments with varying pH conditions. These stringent requirements necessitate innovative drug discovery strategies capable of identifying and optimizing compounds with suitable physicochemical and pharmacokinetic properties.^6^

Historically, phenotypic screening of compound libraries has served as the primary approach for identifying anti-leishmanial agents. While this method has yielded some promising leads, it is often hampered by difficulties in elucidating the molecular targets of active compounds, which can obscure potential off-target effects and complicate downstream optimization. In response, the field has increasingly embraced target-based drug discovery, which enables rational design and iterative optimization around validated molecular targets. However, this strategy also presents challenges—most notably, confirming that observed cellular activity is due to on-target effects and overcoming barriers to compound delivery and efficacy within the parasite’s intracellular niche.^7–9^

To address these obstacles, innovative methodologies that integrate chemical and genetic tools have emerged. Chemoproteomics, for example, facilitates the elucidation of mechanisms of action for compounds identified through phenotypic screens.^10–12^ In parallel, genetic techniques enable the validation of candidate proteins as *bona fide* drug targets.^4,13^ Furthermore, in-cell target engagement assays have become essential for confirming compound binding to putative targets within living cells.^14^ These assays not only demonstrate cellular permeability but also provide direct evidence of target engagement.

While in-cell target engagement assays are well-established for human proteins, such as kinases and deacetylases,^14,15^ their application in infectious disease research has been comparatively limited. Recently, our group demonstrated the utility of these assays in pathogens including Gram-positive bacteria and *Mycobacteria*.^16^ Building on this foundation, we have now developed a target engagement assay for both *Leishmania* promastigotes, the most commonly used form in drug discovery, and intracellular amastigotes, the clinically relevant stage of the parasite.

Using this assay, we identified WZ8040 as a cell-permeable inhibitor of *Leishmania mexicana* CLK1, a key protein kinase involved in regulating the parasite cell cycle.^4^ This compound exhibited potent binding to LmxCLK1 within macrophage-internalized amastigotes and demonstrated antiparasitic activity in the low micromolar range. Our findings highlight the potential of target engagement assays to accelerate the discovery of novel anti-leishmanial agents. We anticipate that similar strategies could be broadly applied to other candidate drug targets in *Leishmania*, paving the way for a new generation of therapeutics against this NTD.

## Results & Discussion

### Development of an in-cell BRET-based target engagement assay in *Leishmania*

CLK1 (KKT10) and its close paralog CLK2 (KKT19) are integral components of a highly divergent kinetochore complex in *Leishmania* and related kinetoplastids, which lacks detectable homology to canonical eukaryotic kinetochore proteins.^17–19^ Chemical inhibition of CLK1/CLK2 with the covalent inhibitor AB1 disrupts kinetochore function, resulting in chromosome missegregation and parasite death in *Trypanosoma brucei*, and leading to defective cytokinesis in *Leishmania*.^20–22^ Notably, LmxCLK1 and LmxCLK2 share 93.4% overall amino acid sequence identity, with their kinase domains—residues 73–422 in LmxCLK1 and 67–416 in LmxCLK2—being completely identical (**Supplementary Figure 1A**). The sequence divergence is confined to their N-terminal regions, spanning approximately 50 residues.

Given their essential redundant role in cell division and conserved active sites, LmxCLK1 and LmxCLK2 represent promising therapeutic targets. To investigate these kinases, all experimental work presented here was performed using full-length LmxCLK1 (TritrypDB LmxM.09.0400, UniProt ID: E9AMP2), with a specific focus on the ATP-binding site within the kinase domain, which is fully conserved between the two paralogs. Consistent with their primary sequence, deep learning-based structural predictions using AlphaFold3^23^ indicate that both proteins possess a canonical kinase domain fold preceded by a less ordered N-terminal region (**Supplementary Figures S1B-C**).

To facilitate the discovery of novel, cell-permeable ligands for LmxCLK1/2, we sought to develop a live-cell, target engagement assay for their ATP-binding sites in *Leishmania*. Bioluminescence resonance energy transfer (BRET)-based target engagement assays have been successfully utilized to study in-cell target engagement in human cells.^14,15^ These assays employ energy transfer between a BRET donor (e.g., NanoLuc, NLuc) and an acceptor fluorophore (e.g., BODIPY) to monitor ligand binding in real time. Commercially available BRET probes based on promiscuous ATP-competitive ligands can target over 340 wild-type human protein kinases, including human CLK1 (HsCLK1), the closest human homolog of LmxCLK1/2,^24,25^ which shares approximately 37 % amino acid sequence identity.

To evaluate whether a commercial HsCLK1 BRET probe (K-5, Promega catalog #N2482) could interact with LmxCLK1 *in vitro*, we assessed its binding affinity using recombinant LmxCLK1. Full-length LmxCLK1 was expressed in *Escherichia coli* as an N-terminal fusion with NLuc, connected via a short, flexible linker (**Supplementary Figures S2A–C**). By titrating probe K-5 against a fixed concentration (125 pM) of purified NLuc::LmxCLK1 in the presence of a NLuc substrate (Promega catalog #N2161), we determined an equilibrium dissociation constant (*K*_D_) of 914.6 nM (95% CI: 751.9–1211.0 nM) in the absence of ATP (**Figure 1A**). These results confirm that the K-5 BRET probe binds to recombinant NLuc::LmxCLK1 *in vitro*.

**Figure 1.**
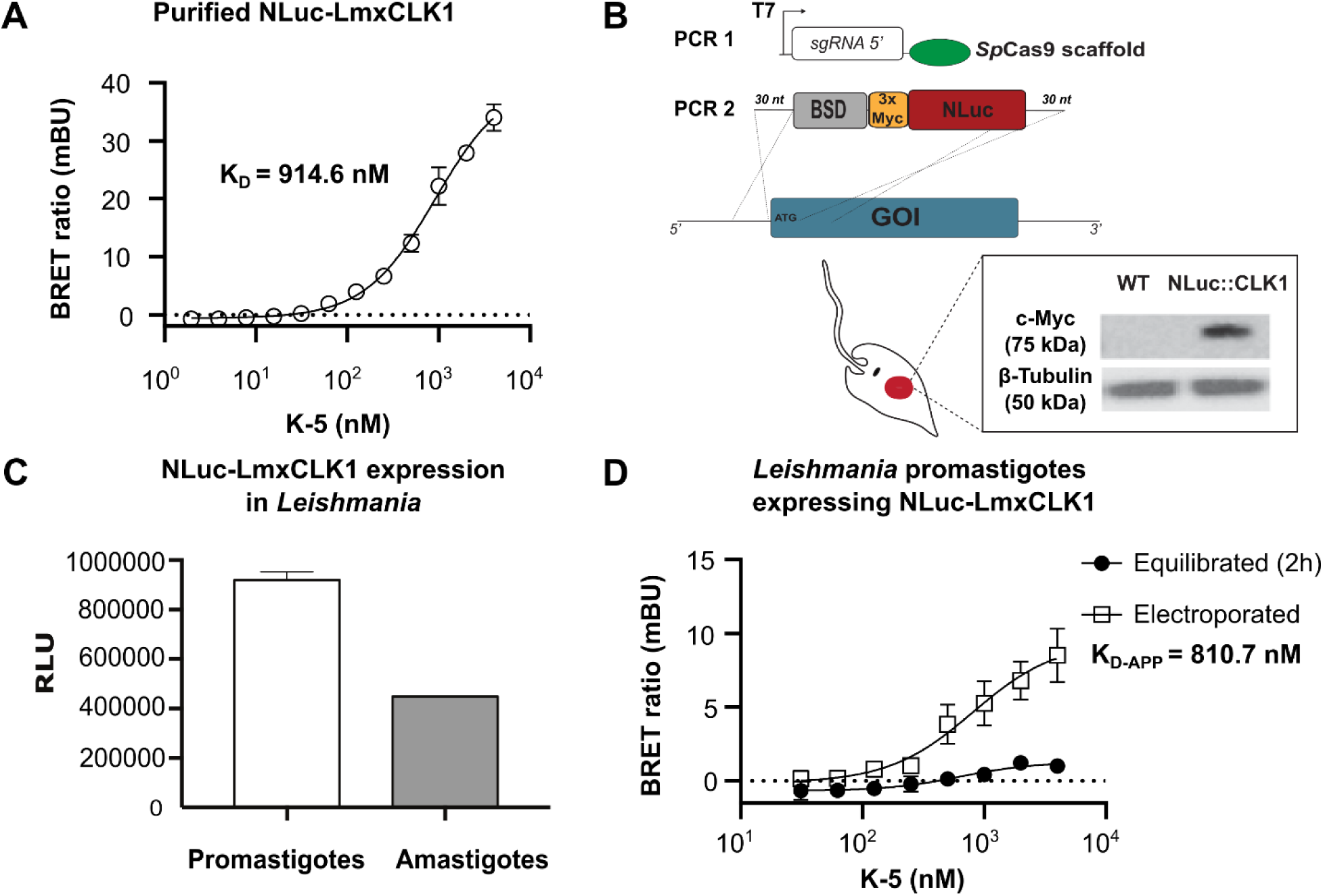
Validation of NanoBRET probe K-5 engagement with NLuc::LmxCLK1 *in vitro* and in cells. (A) Saturation binding of NanoBRET probe K-5 to purified recombinant NLuc::LmxCLK1 at a fixed concentration (125 pM) in the presence of NLuc substrate. The binding curve indicates a dissociation constant (*K*_D_) of 914.6 nM (95% CI: 751.9–1211.0 nM) in the absence of ATP. Data represent mean ± SD from two independent experiments performed in duplicate. The *K*_D_ was determined by fitting the data to a four-parameter dose-response model. (B) CRISPR/Cas9 strategy to generate a *Leishmania* mutant cell line expressing NLuc fused to endogenous CLK1. The sgRNA expression cassette containing a T7 promoter (PCR1) and a repair template containing a blasticidin S deaminase (BSD) and 3x Myc::NLuc (PCR2) were transfected into *L. mexicana* promastigotes. Expression of the fusion protein (∼75 kDa) was confirmed by Western blot using an anti-c-Myc antibody (1:5000). (C) Expression of NLuc::LmxCLK1 in both promastigote and amastigote life stages. Relative luminescence units (RLU) were measured in axenically cultured promastigotes (10⁵ parasites/well) and intramacrophage amastigotes (48 h post-infection at a 10:1 parasite-to-macrophage ratio). Data represent mean ± SD from two independent experiments performed in duplicate. (D) Intracellular BRET signal in promastigotes expressing NLuc::LmxCLK1 following 2 h of K-5 probe electroporation (white squares), but not after passive incubation (black circles). A *K*_D-APP_ of 810.7 nM (95% CI: 426.2–n.d.) was observed for electroporated cells. Data represent mean ± SD from two independent experiments performed in duplicate. The *K*_D-APP_ was determined by fitting the data to a four-parameter dose-response model. CI, confidence interval; *K*_D_, dissociation constant; *K*_D-APP_, apparent dissociation constant; mBU, milliBRET units; n.d., not determined; RLU, relative luminescence units.

Encouraged by these results, we employed a CRISPR/Cas9-based strategy to genetically modify *L. mexicana* to express NLuc fused to the N-terminus of the endogenous LmxCLK1 protein (NLuc::LmxCLK1). This approach ensured that the fusion protein was expressed under native regulatory elements, thereby preserving physiological expression levels and lifecycle-specific regulation. Western blot analysis confirmed that the fusion protein was expressed at the expected molecular weight (**Figure 1B**). Notably, the growth kinetics of NLuc::LmxCLK1-expressing parasites were comparable to those of wild-type cells, indicating that the fusion did not impair parasite viability. Luminescence assays further demonstrated robust luciferase activity in both promastigote and amastigote forms, confirming expression and functionality of the NLuc::LmxCLK1 fusion across parasite life stages (**Figure 1C**).

We next sought to detect the interaction between NLuc::LmxCLK1 and the NanoBRET probe K-5 in genetically modified *L. mexicana* promastigotes. To ensure that any observed BRET signal originated from intracellular interactions, a non-cell-permeable NLuc inhibitor was included in all assays to quench extracellular luminescence.^26^ However, titration of the K-5 probe (up to 4.0 µM) in promastigote culture media yielded minimal BRET signals (**Figure 1D**, black circles), comparable to background levels, regardless of incubation conditions—including variations in media composition and extended incubation times of up to six hours (data not shown). These findings are consistent with the well-documented challenge of poor compound permeability in *Leishmania*, which represents a major bottleneck in drug discovery for these parasites.^9^ Our results suggest that limited probe permeability is a key barrier to establishing a functional BRET assay in this system.

To overcome this limitation, we electroporated the genetically modified promastigotes in the presence of increasing concentrations of the K-5 probe. Under these conditions, we observed a robust, dose-dependent BRET signal, with an apparent dissociation constant (*K*_D-APP_) of 810.7 nM (95% CI: 426.2 nM – not determined) (**Figure 1D**, white squares), closely matching the *K*_D_ obtained using purified protein. These results strongly indicate that the absence of a BRET signal in non-electroporated cells was primarily due to poor probe permeability. Thus, the development of a *Leishmania*-permeable BRET probe will be essential for establishing a reliable, live-cell target engagement assay in this parasite for LmxCLK1.

The K-5 BRET probe consists of a promiscuous kinase-binding ligand, a BODIPY fluorophore, and a linker— each component contributing to the overall physicochemical properties and cellular permeability of the probe. Although K-5 demonstrated specific interaction with purified LmxCLK1 *in vitro*, its limited permeability in promastigotes necessitated the exploration of alternative scaffolds. Designing BRET probes for novel targets in infectious pathogens such as LmxCLK1 poses a significant challenge, primarily due to the scarcity of known ligands that can be adapted for use in probe development. For instance, while the covalent inhibitor AB1 is a potent and selective inhibitor of CLK1 in trypanosomatids,^20^ it was unsuitable as a BRET probe scaffold due to its irreversible binding mechanism, which precludes its use in competitive displacement assays.

To overcome this limitation, we expressed full-length recombinant full-length LmxCLK1 in *E. coli* (**Supplementary Figure S2D–F**) and employed a thermal shift assay (differential scanning fluorimetry, DSF),^27,28^ to screen a commercial library of approximately 380 human kinase inhibitors for binding to the purified protein (**Figure 2A** and **Supplementary Table S3**). Twelve compounds induced significant thermal stabilization (ΔT_m_ > 3.0 °C), GZD824, a reversible ATP-competitive inhibitor of Bcr-Abl,^29^ producing the largest shift (ΔT_m_ = 17.4 ± 0.3 °C). This stabilization exceeded that of staurosporine, a known promiscuous kinase inhibitor (ΔT_m_ = 10.1 ± 0.2 °C).

**Figure 2.**
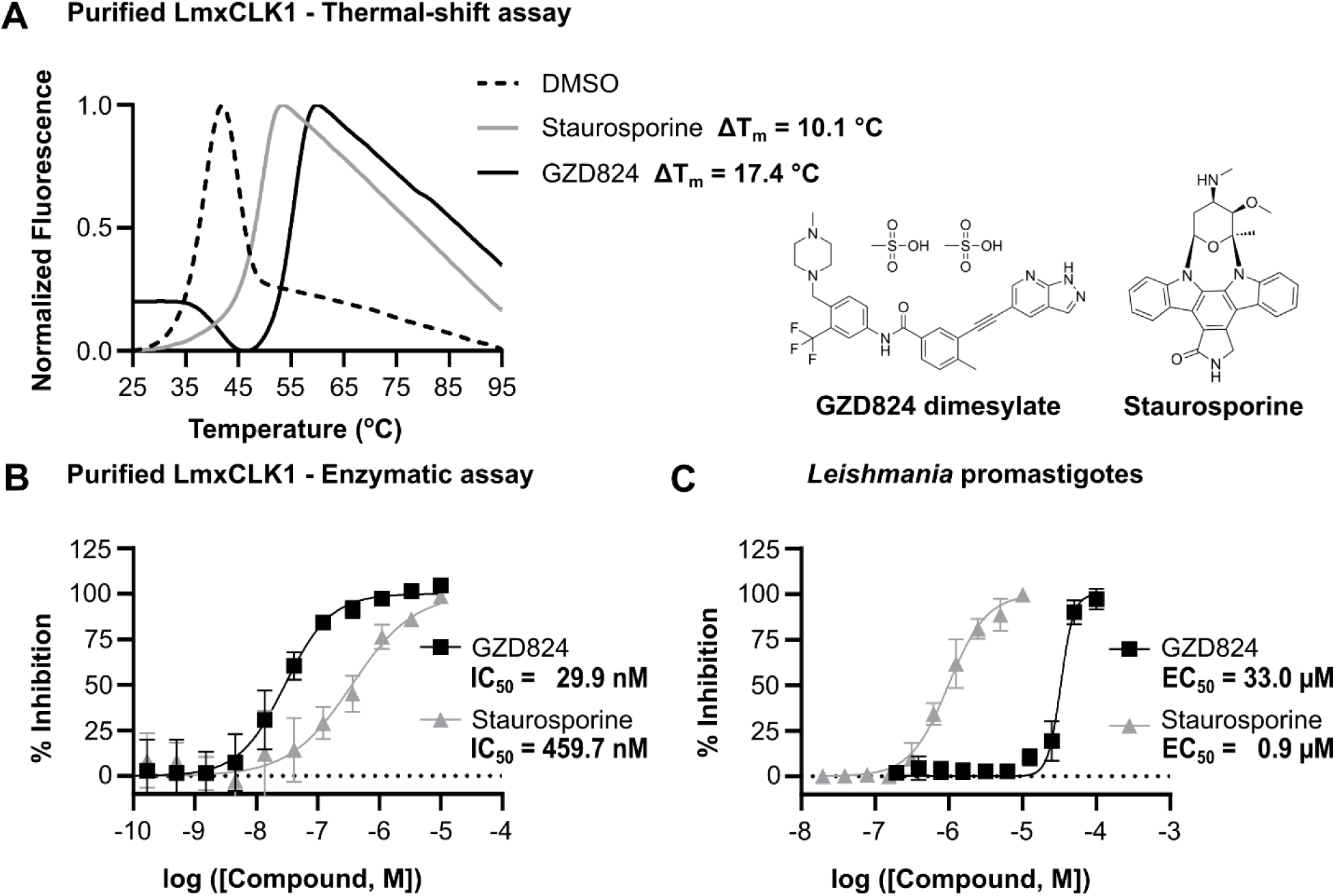
Identification of a LmxCLK1 ligand scaffold for BRET probe development. (A) Thermal melting curves for GZD824 and staurosporine, normalized and averaged from two independent experiments. Ligands were tested at a fixed concentration (10 µM), resulting in a rightward shift in the protein unfolding transition. The melting temperature shift (ΔTm) was calculated as the difference in temperature at which 50% relative fluorescence was observed, using DMSO as a reference. GZD824 and staurosporine induced ΔTm values of 17.4 ± 0.3 °C and 10.1 ± 0.2 °C, respectively. Chemical structures of both compounds are shown. (B) Enzymatic inhibition of LmxCLK1 kinase by GZD824 and staurosporine. Dose–response curves were generated from blank-corrected fluorescence intensity measurements, normalized and fitted using a four-parameter dose-response model. GZD824 and staurosporine inhibited LmxCLK1 with IC₅₀ values of 29.9 nM (95% CI: 20.9–43.9 nM) and 459.7 nM (95% CI: 255.9–1411.0 nM), respectively. Data represent mean ± SD of two independent experiments performed in duplicate. (C) Anti-leishmanial activity of GZD824 and staurosporine in *L. mexicana* promastigotes after 48 h of compound exposure. Dose– response curves were used to calculate EC₅₀ values, with GZD824 showing an EC₅₀ of 33.0 µM (95% CI: 30.5–35.2 µM) and staurosporine an EC₅₀ of 0.9 µM (95% CI: 0.8–1.0 µM), calculated using a using a four-parameter dose-response model. Data represent mean ± SD of two independent experiments performed in triplicate.

To validate these findings, we established a TR-FRET-based enzymatic assay to assess the potency of GZD824. This assay measured the phosphorylation of a generic peptide substrate (STK-Substrate 3; Cisbio, #61ST3BLE) by purified full-length LmxCLK1 (**Supplementary Figure S3**). GZD824 inhibited LmxCLK1 with an IC_50_ of 29.9 nM (95% CI 20.9 - 43.9 nM), while staurosporine exhibited a significantly higher IC_50_ of 459.7 nM (95% CI 255.9 - 1411.0 nM) (**Figure 2B**). Using the Cheng-Prusoff equation,^30^ we calculated inhibitory constant (K*_i_*) values of 2.7 nM for GZD824 and 41.8 nM for staurosporine. These results establish GZD824 as a potent LmxCLK1 inhibitor.

Next, we evaluated the cellular permeability of GZD824 by assessing its ability to reduce the viability of *L. mexicana* promastigotes. Cell viability was measured using the resazurin reduction assay (Alamar Blue), which quantifies the conversion of resazurin to resorufin as an indicator of metabolic activity.^31^ GZD824 demonstrated anti-leishmanial activity with an EC_50_ of 33.0 µM (95% CI: 30.5–35.2 µM), indicating that it can permeate *Leishmania* cells. However, its potency was modest compared to the pan-kinase inhibitor staurosporine, which exhibited an EC_50_ of 0.9 µM (95% CI: 0.8–1.0 µM) (Figure 2C). Taken together, these enzymatic and phenotypic assays strongly support that GZD824 binds to LmxCLK1 and is capable of penetrating *Leishmania* promastigotes.

Building on these results, we synthesized two BRET probes—SV363 and SV366—each incorporating GZD824 as the ligand, conjugated to either EverFluor590 (a BODIPY-based dye) or 5-carboxyfluorescein (5-FAM), respectively (**Figure 3A**). Among the two, SV366 demonstrated markedly stronger binding to purified NLuc::LmxCLK1, with an EC₅₀ of 8.4 nM. In contrast, SV363 exhibited substantially weaker binding (EC₅₀ > 1 µM) and was therefore not pursued further as a candidate BRET probe. Further characterization of SV366 confirmed that its binding could be competitively displaced by unmodified GZD824 and staurosporine (**Figures 3B-C**), yielding a *K*_D_ of 5.9 nM (95% CI: 3.9–8.9 nM).

**Figure 3.**
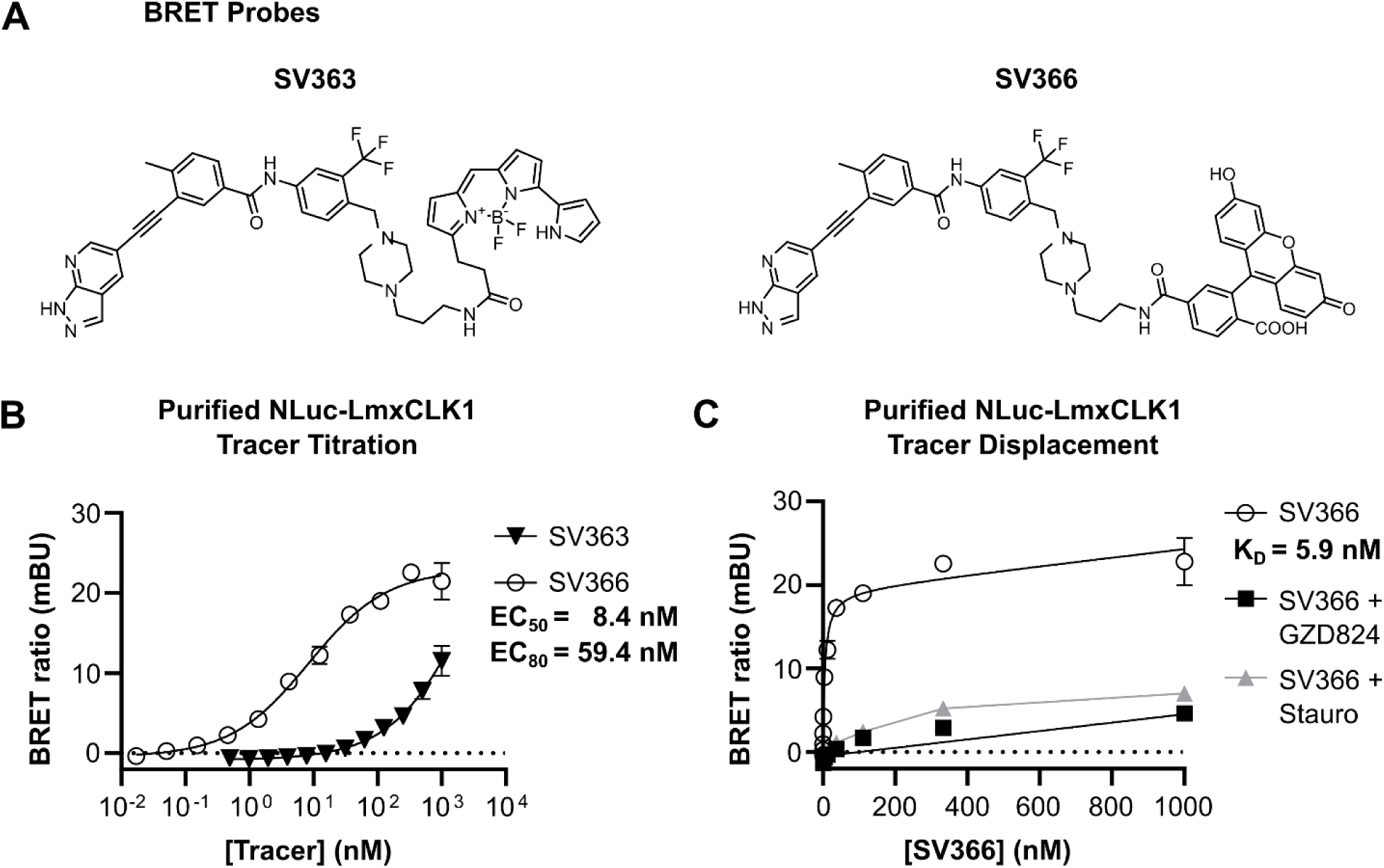
Characterization of new BRET probes targeting LmxCLK1. (A) Chemical structures of SV363 (GZD824 conjugated to Everfluor590) and SV366 (GZD824 conjugated to 5-Carboxyfluorescein). (B) Concentration-dependent binding of SV363 and SV366 to purified NLuc::LmxCLK1. Data represent mean ± SD of two independent experiments performed in duplicate. For SV366, the calculated EC₅₀ was 8.4 nM (95% CI: 5.8–12.7 nM), and the EC₈₀ was 59.4 nM (95% CI: 31.9–143.1 nM). EC values were calculated using a using a four-parameter dose-response model. (C) Saturation binding of SV366 to purified NLuc::LmxCLK1 in the presence or absence of 50 µM of the competitive inhibitors GZD824 and staurosporine. The dissociation constant (*K*_D_) of 5.9 nM (95% CI: 3.9–8.9 nM) was determined by nonlinear regression using a one-site binding model that accounts for non-specific binding. Data represent mean ± SD of two independent experiments performed in duplicate.

Finally, we evaluated the performance of BRET tracer SV366 in live *L. mexicana* cells. In promastigotes, titration of increasing concentrations of SV366 directly into the culture medium produced dose-dependent BRET signals, with an EC_50_ of 383.2 nM (95% CI: 291.2–634.8 nM) (**Figure 4A**). Target specificity was confirmed through competitive displacement assays using unmodified GZD824, which effectively reduced the BRET signal and yielded an IC_50_ of 4.6 µM (95% CI: 3.6–6.0 µM; **Figure 4B**). Similarly, in infected macrophages, SV366 generated probe-dependent BRET signals in intracellular amastigotes, with an EC_50_ of 72.6 nM (95% CI: 64.0–82.9 nM) (**Figure 4C**). GZD824 also displaced SV366 in a dose-dependent manner within macrophage-internalized amastigotes, yielding an IC_50_ of 13.0 µM (95% CI: 11.1–15.6 µM; **Figure 4D**).

**Figure 4.**
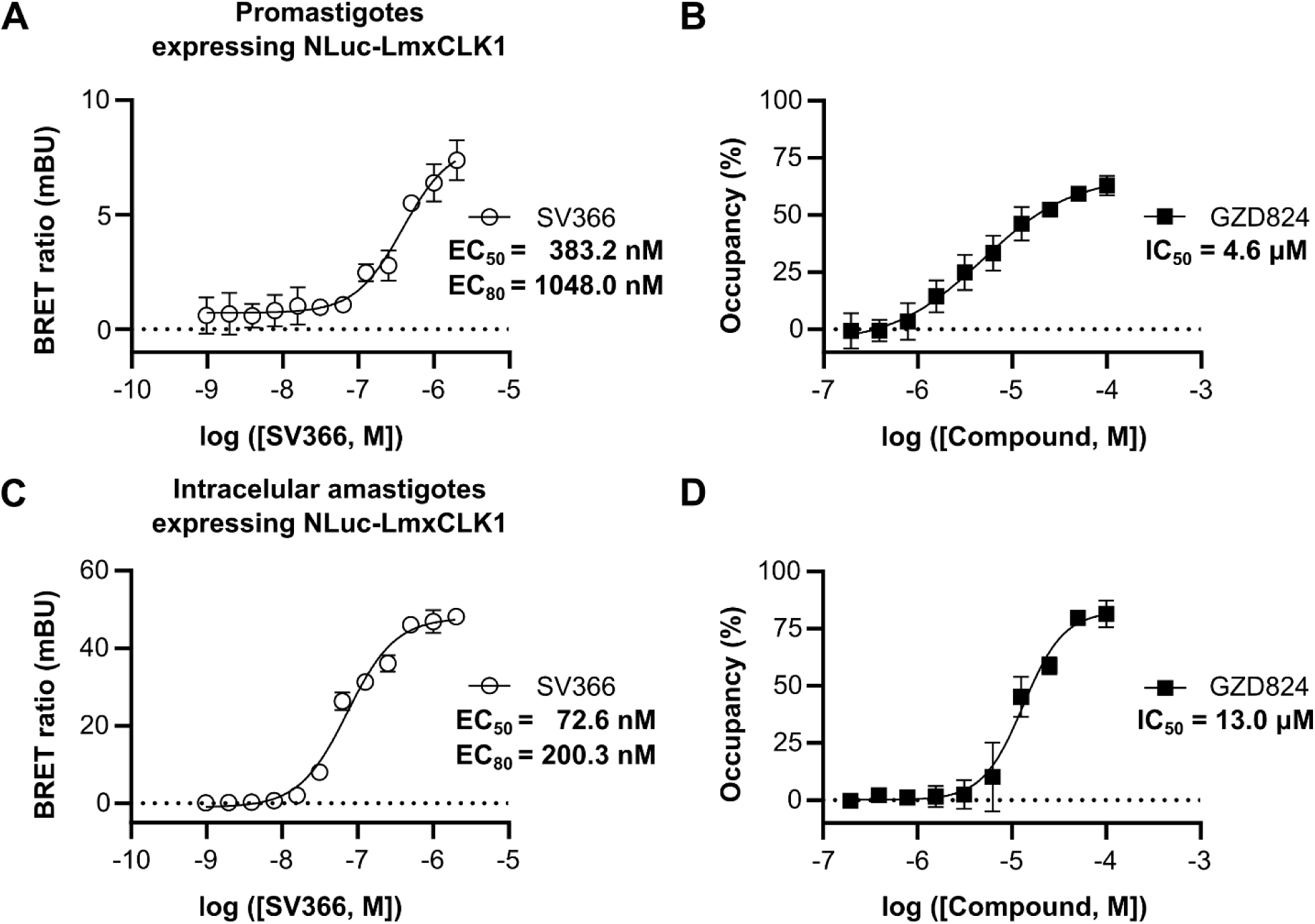
Validation of compound target specificity in live *L. mexicana* promastigotes and intracellular amastigotes. (A) Concentration-dependent intracellular BRET signal following 1-hour incubation with SV366 in *L. mexicana* promastigotes expressing NLuc::LmxCLK1. Data represent mean ± SD of two independent experiments performed in triplicate. The calculated EC_50_ was 383.2 nM (95% CI: 291.2–634.8 nM), and the EC_80_ was 1048.0 nM (95% CI: 640.9–2629.0 nM). (B) Dose-dependent displacement of SV366 by GZD824 in promastigotes. Data represent mean ± SD of two independent experiments performed in triplicate. The estimated IC_50_ for GZD824 was 4.6 µM (95% CI: 3.6–6.0 µM). (C) Concentration-dependent intracellular BRET signal following 1-hour incubation with SV366 in intracellular amastigotes expressing NLuc::LmxCLK1. Data represent mean ± SD of two independent experiments performed in duplicate. The calculated EC₅₀ was 72.6 nM (95% CI: 64.0–82.9 nM), and the EC_80_ was 200.3 nM (95% CI: 160.3–257.2 nM). (D) Dose-dependent displacement of SV366 by GZD824 in intracellular amastigotes. Data represent mean ± SD of two independent experiments performed in duplicate. The estimated IC_50_ for GZD824 was 13.0 µM (95% CI: 11.1–15.6 µM).

The approximately 5-fold lower EC_50_ observed for SV366 in intracellular amastigotes compared to promastigotes likely reflects differences in cellular context, such as altered membrane permeability, intracellular accumulation, or enhanced probe retention within the internalized amastigotes. Although fluorescein fluorescence is known to diminish at acidic pH,^32,33^ the observed BRET signal confirms that SV366 remains fluorescent after traversing the vacuole (pH 5.5), and successfully reaches the parasite cytoplasm, where the pH is near-neutral (∼7.4).^9^ In contrast, the unmodified parent compound GZD824 exhibited ∼3-fold lower potency in intracellular amastigotes, suggesting that its intracellular availability is not similarly enhanced. This divergence highlights the impact of fluorophore conjugation on intracellular pharmacokinetics and supports the utility of SV366 as a robust and functional BRET probe for quantifying target engagement in both extracellular and intracellular *Leishmania* stages.

### Identification of new, cell permeable ligands of LmxCLK1

To identify LmxCLK1 ligands capable of displacing SV366 in live *Leishmania* cells, we screened the same library of approximately 380 human kinase inhibitors previously evaluated by DSF (**Figure 5A** and **Supplementary Table S3**), using a BRET-based assay in promastigotes with SV366 as the tracer (**Figure 5B** and **Supplementary Table S4**). All compounds were tested at a fixed concentration of 10 µM to enable direct comparison of their target occupancy in live cells.

**Figure 5.**
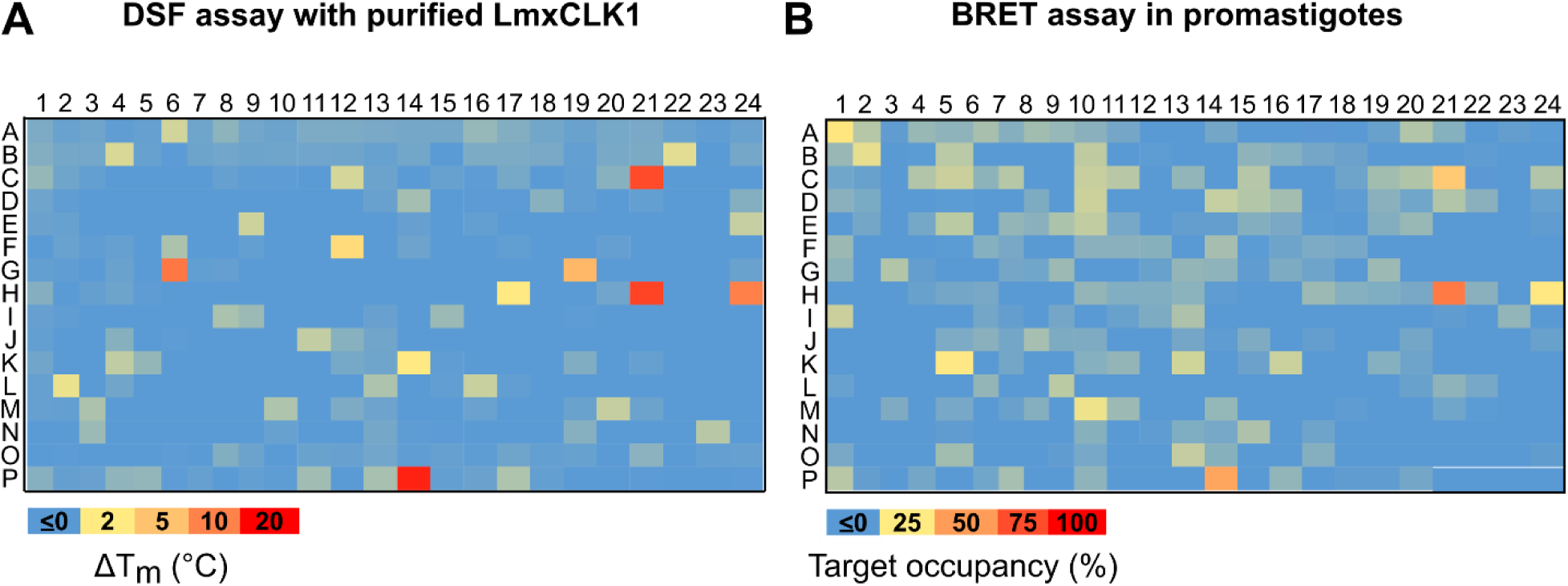
Discovery of cell-permeable ligands targeting LmxCLK1. (A) Heatmap from DSF screening of approximately 380 human kinase inhibitors against recombinant LmxCLK1. Assays were performed using 2 µM protein and 10 µM of each compound. Thermal shift (ΔTm) values were calculated based on the temperature at which 50% relative fluorescence was observed, using DMSO as a reference. Data represent the mean of two independent experiments. ΔTm values are visualized using a color gradient from blue (low shift) to red (high shift), as indicated below the heatmap. (B) Heatmap from BRET-based target engagement assay in *L. mexicana* promastigotes expressing NLuc::LmxCLK1. Parasites were incubated with 1 µM SV366 and 10 µM of each compound. Data represent the mean of two independent experiments. Compound target occupancy is represented using a color gradient from blue (low engagement) to red (high engagement), as shown below the heatmap.

Comparison of the cellular BRET data with DSF results revealed a lack of direct correlation between thermal stabilization (ΔT_m_) and target engagement in live cells. For instance, GZD824, which produced the highest T_m_ shift (ΔT_m_ = 17.4 ± 0.3 °C), achieved only moderate target occupancy (46.5 ± 3.2%) at 10 µM in live cells. In contrast, WZ8040, which induced a smaller T_m_ shift (ΔT_m_ = 14.8 ± 0.1 °C), exhibited higher target occupancy (60.8 ± 0.8%) under the same conditions. Similarly, WZ3146 produced a comparable T_m_ shift (ΔT_m_ = 14.3 ± 0.1 °C) but showed substantially lower target occupancy (35.2 ± 3.3%). Notably, Pelitinib and staurosporine, despite having similar T_m_ shifts (ΔT_m_ = 10.8 ± 1.7 °C and 10.1 ± 0.2 °C, respectively), displayed markedly different behaviors in live cells: Pelitinib showed negligible target engagement, whereas staurosporine achieved an occupancy of 25.4 ± 2.0%.

These findings underscore the limitations of relying solely on assays with purified proteins, such as DSF, to identify compounds with cellular activity. While DSF provides valuable insights into ligand binding and protein stabilization, it does not account for critical factors such as compound permeability, intracellular distribution, or metabolic stability within the cellular environment. The discrepancies observed between thermal shifts and target occupancy in live cells highlight that protein-ligand binding alone is not always predictive of cellular efficacy.

In contrast, BRET-based target engagement assays in *Leishmania* offer an efficient approach to overcoming these limitations by directly measuring target engagement in live parasites. Integrating this method into the drug discovery workflow bridges the gap between *in vitro* binding data and *in vivo* cellular activity. This approach enables the prioritization of *Leishmania*-permeable ligands and accelerates the identification, validation, and development of promising therapeutic candidates to treat leishmaniasis.

### Characterization of WZ8040 as a potent, covalent inhibitor of LmxCLK1 with anti-leishmanial activity

Based on the results of our BRET assay, we selected WZ8040 and WZ3146 for further characterization (Figure 6A). Both compounds are covalent, irreversible inhibitors of human EGFR.^34,35^ Covalent inhibitors may offer distinct advantages in the treatment of infectious diseases due to their ability to form irreversible bonds with target proteins, resulting in sustained inhibition even at lower systemic concentrations. This property is particularly beneficial in the context of pathogenic infections, where prolonged target engagement may help overcome challenges such as rapid pathogen replication, limited drug exposure in infected tissues, and the emergence of resistance.^36–38^

**Figure 6.**
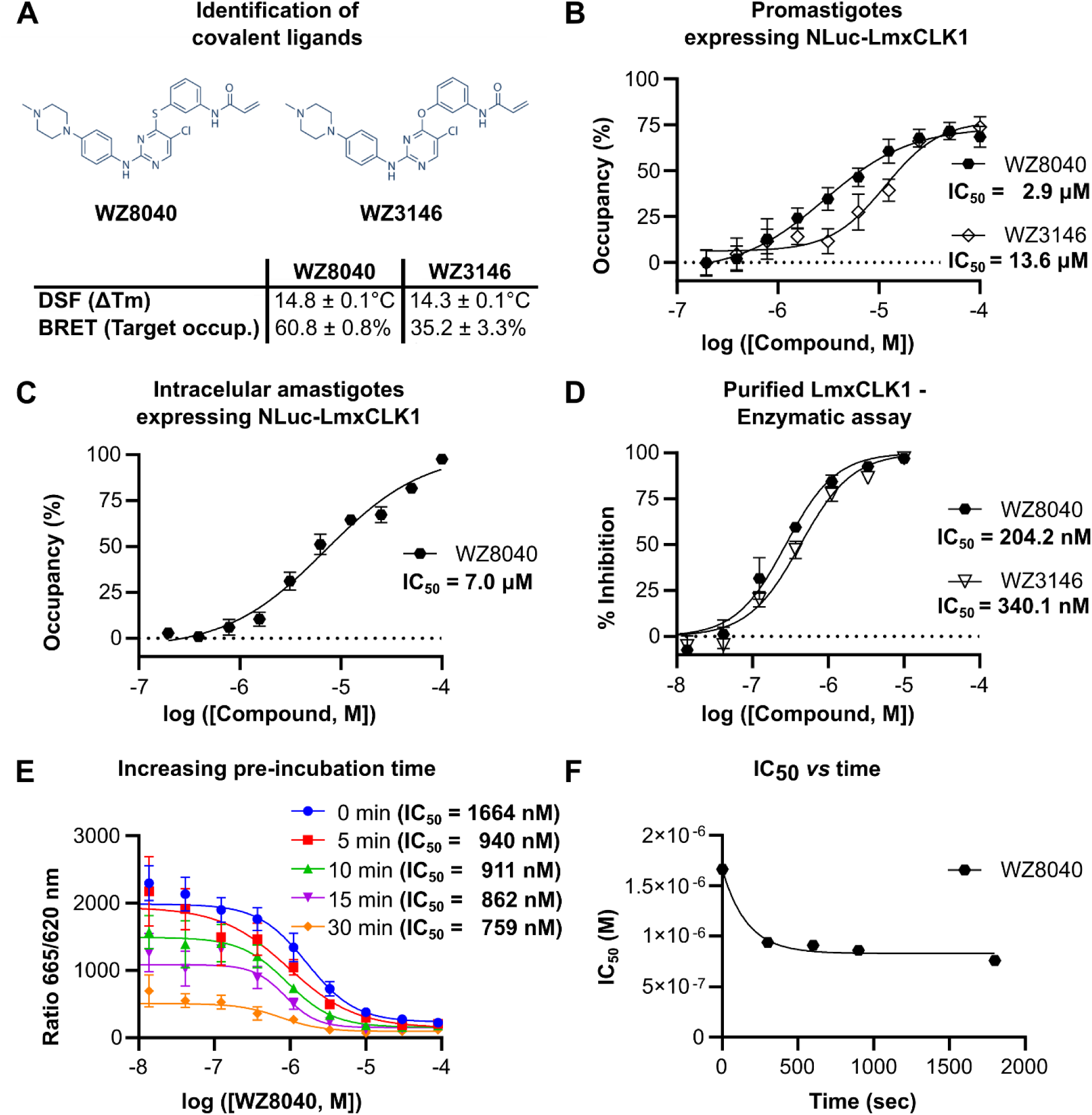
Identification and characterization of covalent LmxCLK1 inhibitors. (A) Chemical structures of covalent inhibitors WZ8040 and WZ3146, identified through both DSF and BRET-based screening assays. (B) Dose-dependent displacement of tracer SV366 by WZ8040 and WZ3146 in *L. mexicana* promastigotes expressing NLuc::LmxCLK1. Data represent mean ± SD of two independent experiments performed in triplicate. IC₅₀ values were 2.9 µM (95% CI: 2.3–3.6 µM) for WZ8040 and 13.6 µM (95% CI: 10.6–17.6 µM) for WZ3146. (C) Dose-dependent displacement of SV366 by WZ8040 in intracellular amastigotes expressing NLuc::LmxCLK1. Data represent mean ± SD of two independent experiments performed in duplicate. The calculated IC₅₀ was 7.0 µM (95% CI: 5.0–12.7 µM). (D) *In vitro* LmxCLK1 kinase inhibition by WZ8040 and WZ3146. Blank-corrected fluorescence intensity data were fitted using a four-parameter dose-response model. Data represent mean ± SD of two independent experiments performed in duplicate. (E) Time-dependent inhibition of LmxCLK1 by WZ8040. IC₅₀ values decreased with increasing pre-incubation times (0, 5, 10, 15, and 30 minutes), consistent with covalent binding kinetics. (F) Kinetic analysis of time-dependent IC₅₀ values for WZ8040, fitted to an irreversible inhibition model.

To assess the cell permeability of WZ8040 and WZ3146, we used SV366 in BRET-based assays in promastigotes. WZ8040 and WZ3146 exhibited IC_50_ values of 2.9 µM (95% CI: 2.3–3.6 µM) and 13.6 µM (95% CI: 10.6–17.6 µM), respectively (**Figure 6B**), confirming that both compounds effectively bind to LmxCLK1 within cultured *Leishmania* promastigotes. These results were consistent with single-concentration BRET assays, which showed greater target engagement by WZ8040 compared to WZ3146. Furthermore, in macrophage-internalized *Leishmania* amastigotes, WZ8040 demonstrated an IC_50_ of 7.0 µM (95% CI: 5.0–12.7 µM) using the same BRET-based assay (**Figure 6C**), validating its cell permeability and target engagement in the clinically relevant amastigote stage.

To confirm the enzymatic inhibition of LmxCLK1 by WZ8040 and WZ3146, we employed a TR-FRET-based enzymatic assay. This assay yielded IC_50_ values of 204.2 nM (95% CI: 163–260.1 nM) for WZ8040 and 340.1 nM (95% CI: 263.5–453.6 nM) for WZ3146 (**Figure 6D**), indicating that both compounds are potent inhibitors of LmxCLK1. Mass spectrometry further validated the formation of covalent adducts between the compounds and purified LmxCLK1. Incubation with the compounds resulted in mass shifts consistent with the addition of a single molecule of WZ8040 (+481.01 Da) or WZ3146 (+464.95 Da) (**Supplementary Figure S4**). Additionally, to establish that WZ8040 acts through a covalent mechanism, we evaluated the time dependency of its inhibitory activity (**Figure 6E**). As anticipated, the IC_50_ values of WZ8040 decreased with longer incubation times (**Figure 6F**), consistent with time-dependent covalent inhibition. However, under the conditions of our enzymatic assay, we were unable to reliably determine the kinetic parameters *K*_I_/*K*_inact_, which are critical for quantifying the potency of mechanism-based covalent inhibitors.^39,40^

To identify the site of covalent modification in LmxCLK1, we analyzed tryptic peptides of compound-treated protein using LC-MS/MS. Database searches allowing for carbamidomethylation and WZ8040 adducts at cysteine residues revealed high overall sequence coverage (92%) and identified four peptides that contained the WZ8040 modification (**Supplementary Figures S5-7**). Our analysis indicates that Cys172 is the primary site of covalent modification, with additional, lower-frequency adducts detected at Cys240, Cys245, and Cys327 (**Supplementary Figure S7**). Both Cys172 and Cys245 are located within the ATP-binding pocket of LmxCLK1, whereas Cys240 and Cys327 lie outside this region (**Supplementary Figure S8A,B**).

Notably, residue Cys172 in *L. mexicana* CLK1 is also targeted by the covalent inhibitor AB1.^22^ Moreover, this residue is structurally equivalent to Cys215 in *T. brucei* CLK1, which is also targeted by AB1,^20^ and to Cys797 in human EGFR, the known covalent binding site of WZ4002, a close analog of WZ8040 (**Supplementary Figure S8A,B**).^34^ Importantly, cysteines positioned near the conserved DFG motif—such as Cys172—have been reported as reactive sites for other covalent kinase inhibitors, including SM1-71.^41,42^

We next assessed whether WZ8040 could reduce the viability of *L. mexicana* promastigotes using the resazurin reduction assay (Alamar Blue).^31^ WZ8040 significantly decreased the number of metabolically active promastigotes, with an EC_50_ value of 7.9 µM (95% CI: 6.6–9.5 µM) (**Figure 7A**). For comparison, staurosporine and GZD824 exhibited EC_50_ values of 0.9 µM (95% CI: 0.8–1.0 µM) and 33.0 µM (95% CI: 30.5–35.2 µM), respectively (**Figure 2C**). These results demonstrate that WZ8040 is effective against the promastigote stage, the predominant form of the parasite within the insect vector.

**Figure 7.**
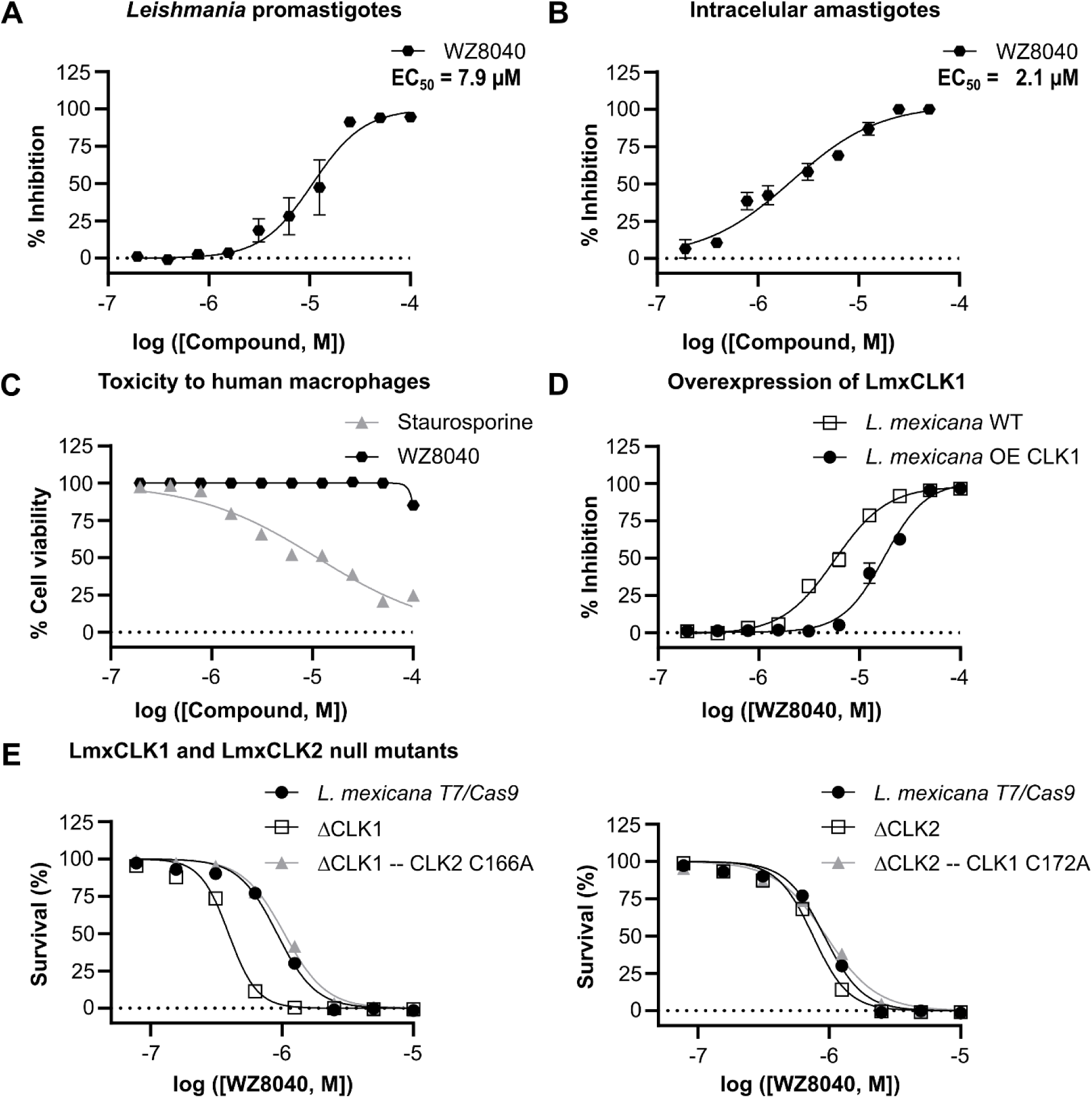
Phenotypic validation of WZ8040 activity in *L. mexicana* promastigotes and intracellular amastigotes. (A) Determination of the EC₅₀ of WZ8040 in promastigotes using a resazurin-based viability assay. The calculated EC₅₀ was 7.9 µM (95% CI: 6.6–9.5 µM). Data represent mean ± SD of four independent experiments performed in triplicate. (B) Dose–response curve for WZ8040 in intracellular amastigotes, based on quantification of total amastigotes in 250 THP-1-derived macrophages. The estimated EC₅₀ was 2.1 µM (95% CI: 1.4–3.4 µM). Data represent mean ± SD of two independent experiments performed in duplicate. (C) Cytotoxicity assessment of WZ8040 in THP-1-derived macrophages using a resazurin assay, with staurosporine as a positive control. Data represent mean ± SD of two independent experiments performed in triplicate. (D) Target validation of LmxCLK1 through EC₅₀ shift analysis in *L. mexicana* promastigotes. Overexpression of LmxCLK1 (filled symbols) reduced sensitivity to WZ8040, with an EC₅₀ of 17.6 µM (95% CI: 16.1–19.2 µM), compared to wild-type parasites (empty symbols; EC₅₀ = 5.6 µM, 95% CI: 5.3–6.0 µM), indicating a ∼3-fold shift. (E) Sensitivity of LmxCLK1 and LmxCLK2 null mutants (ΔCLK1 and ΔCLK2) to WZ8040. ΔCLK1 parasites were significantly more sensitive (EC₅₀ = 0.39 µM, 95% CI: 0.37–0.41 µM) than the parental T7/Cas9 line (EC₅₀ = 0.92 µM, 95% CI: 0.86–0.98 µM). In contrast, ΔCLK2 showed no significant difference (EC₅₀ = 0.75 µM, 95% CI: 0.70–0.81 µM). Interestingly, introducing the structurally equivalent Cys166Ala mutation in LmxCLK2 within the ΔCLK1 background restored sensitivity to parental levels (EC₅₀ = 1.04 µM, 95% CI: 0.96–1.13 µM). For the ΔCLK2 with Cys172Ala mutation in LmxCLK1, no significant difference was observed (EC₅₀ = 0.95 µM, 95% CI: 0.89–1.01 µM). The viability of the mutants and their progenitor lines after 96h treatment with AB1 (0–80 µM) was assessed by measuring the conversion of resazurin (Alamar blue) to resorufin. Values are mean ± SEM.

We then evaluated the efficacy of WZ8040 against *L. mexicana* intracellular amastigotes. Human THP-1 macrophages were infected with *L. mexicana* and treated with increasing concentrations of WZ8040 for 48 hours. Visual inspection and quantification of intracellular amastigotes revealed an EC_50_ value of 2.1 µM (95% CI: 1.4–3.4 µM) (**Figure 7B**), indicating potent activity against the clinically relevant stage of the parasite.

A major challenge in developing kinase inhibitors for infectious diseases is the high conservation of ATP-binding sites across kinases, which can result in off-target effects and host toxicity. However, WZ8040 is a third-generation EGFR inhibitor that selectively targets the T790M mutant form of EGFR, exhibiting over 10-fold selectivity compared to the wild-type enzyme.^34^ In our studies, WZ8040 demonstrated minimal toxicity toward human macrophages (THP-1 cells) used in the amastigote viability assays. Even at a high concentration of 100 µM—approximately 50-fold higher than its EC_50_ against *Leishmania* amastigotes— WZ8040 caused only a modest reduction in macrophage viability (∼15%) (**Figure 7C**). To minimize potential host cell toxicity, we limited the maximum concentration of WZ8040 used in amastigote viability assays to 50 µM.

To further confirm that the anti-leishmanial activity of WZ8040 results from on-target inhibition of LmxCLK1, we generated a genetically engineered *L. mexicana* line overexpressing LmxCLK1, using the same strain employed in all previous phenotypic-based assays. Overexpression of LmxCLK1 reduced parasite sensitivity to WZ8040 by approximately 3-fold compared to wild-type parasites, with EC_50_ values of 17.6 µM (95% CI: 16.1–19.2 µM) and 5.6 µM (95% CI: 5.3–6.0 µM), respectively (**Figure 7D**).

As discussed above, the *Leishmania* genome encodes two closely related CLK paralogs having identical kinase domains. To assess the relative contribution of LmxCLK1 and LmxCLK2 to WZ8040 sensitivity, we utilized previously generated T7/Cas9 knockout lines for each gene.^4^ Interestingly, deletion of LmxCLK1 significantly enhanced sensitivity to WZ8040, whereas deletion of LmxCLK2 showed similar effect compared to the parental T7/Cas9 line (**Figure 7E**). A similar pattern of enhanced sensitivity in the CLK1 knockout line was observed with the covalent inhibitor AB1, further supporting CLK1 as the primary target of these compounds in *Leishmania*.^22^ This differential susceptibility may reflect lower expression levels of CLK2 in promastigotes, making CLK1 the predominant functional target under normal conditions. Alternatively, CLK1 and CLK2 may differ in their interaction partners, or regulatory dynamics, which could influence their accessibility to WZ8040 or their functional importance in kinetochore signaling. It is also possible that compensatory mechanisms are more readily activated in the absence of CLK2 than CLK1.

To evaluate the contribution of covalent binding to WZ8040 efficacy against *Leishmania*, we used CLK2 knockout cell lines expressing the Cys172Ala mutant of LmxCLK1^22^ (**Figure 7E**). Unexpectedly, these mutants exhibited similar sensitivity to WZ8040 as both the parental line and CLK2 knockout parasites expressing wild-type LmxCLK1. This contrasts sharply with previous findings showing that engineered *Leishmania* lacking CLK2 and expressing Cys172Ala LmxCLK1 are over 100-fold less sensitive to the covalent inhibitor AB1.^22^ These results suggest that covalent modification at Cys172 is not essential for WZ8040 activity, raising the possibility that the compound acts through non-covalent interactions or engages alternative cysteine residues in LmxCLK1. However, mass spectrometry analysis identified only three additional cysteines—Cys240, Cys245, and Cys327 (out of 11 total cysteines in LmxCLK1)—that were minimally modified upon compound treatment.

### Conclusion

In conclusion, we have developed a BRET-based target engagement assay to identify cell-permeable, bioactive compounds targeting *Leishmania* CLK1, a protein kinase implicated in cell cycle control and part of the parasite’s highly divergent kinetochore complex. This platform enables direct assessment of compound binding in live parasites, providing critical insight beyond biochemical potency alone. By confirming intracellular target engagement, the assay helps de-risk compound progression by prioritizing molecules that are both active and cell-permeable. This approach is particularly valuable in the context of NTD drug discovery, where resources are limited and early prioritization of viable compounds is essential. Applying this strategy, we demonstrated that WZ8040—a selective inhibitor of mutant human EGFR— engages *Leishmania* CLK1 in intracellular amastigotes, resulting in parasite clearance while sparing host macrophages. These findings highlight the utility of BRET-based assays for validating target engagement in live parasites and underscore their potential to accelerate the development of therapeutics against high-value targets in infectious diseases.

## Materials and Methods

### Bacterial strains and DNA cloning

Routine DNA cloning was performed using *E. coli* Mach-1 cells (Invitrogen, Carlsbad, USA). Recombinant protein expression was carried out in *E. coli* BL21(DE3)-R3-λ-PPase cells.^43^ This chloramphenicol-resistant derivative of *E. coli* BL21(DE3) carries plasmids for the co-expression of λ-phosphatase and three tRNAs corresponding to rarely used codons in *E. coli* for Arg, Ile, and Leu (plasmid pACYC-LIC+). Information on plasmids and oligonucleotides is provided in **Supplementary Tables S1 and S2**, respectively.

To generate a NLuc-fused LmxCLK1 construct (CQMED clone ID: LmxCLK1-cb020) the coding sequence for the full-length *L. mexicana* CLK1 protein (residues Met1 to Met422, TritrypDB LmxM.09.0400, UniProt ID: E9AMP2) was PCR-amplified from *L. mexicana* (strain MNYC/BZ/62/M379) genomic DNA using oligonucleotides LmxCLK1-fb002 and LmxCLK1-rb002. The amplicons and the pFN31A backbone vector (Promega) were digested with the restriction enzymes *Sgf*I and *Pme*I (Promega), and ligated using T4 DNA ligase overnight at 4 °C. After heat inactivation of the ligase (65 °C for 10 minutes), the assembled construct was transformed into *E. coli* Mach1 cells. Positive clones were identified by colony PCR, and the insert was verified by Sanger sequencing using primers NLUC-N-terminal and Flexi-Rev-Seq.

For recombinant production of NLuc-fused LmxCLK1 (residues Met1 to Met422) in *E. coli*, the coding sequence for the fusion protein was PCR-amplified using LmxCLK1-cb020 (see above) as template and oligonucleotide primers NLuc_M1_LIC_Fwd and LmxCLK1-rb001. The resulting amplicons were introduced into expression vector pNIC28-Bsa4 (GenBank ID: EF198106) via ligation-independent cloning, using previously described procedures (CQMED clone ID: LmxCLK1-cb021).^44,45^ This cloning strategy introduced a N-terminal 6xHis tag to the recombinant protein sequence, which is cleavable by treatment with the tobacco etch virus (TEV) protease. Following TEV treatment, the recombinant protein has an additional N-terminal serine residue. Resulting constructs were introduced into Mach1 *E. coli* cells via heat shock and positive clones were identified via colony PCR and verified by DNA sequencing using vector-specific primers: pLIC-forward and pLIC-reverse.

For recombinant production of LmxCLK1 (residues Met1 to Met422) in *E. coli*, the coding sequence was PCR-amplified from *L. mexicana* genomic DNA using oligonucleotide primers LmxCLK1-fb001 and LmxCLK1-rb001. The resulting amplicons were introduced into expression vector pNIC28-Bsa4 (GenBank ID: EF198106) via ligation-independent cloning as described above (CQMED clone ID: LmxCLK1A-cb001).

For overexpression in *L. mexicana*, LmxCLK1 was cloned into pNUS-6myc-G418 by PCR amplification of the coding sequence with oligonucleotides LmxCLK1-fb010 and LmxCLK1-rb010, followed by digestion with restriction endonuclease AvrII (NEB) of the linear insert and of the backbone plasmid, and ligation with T4 ligase (NEB) overnight at 4 °C, before being inserted into Mach1 *E. coli* cells (CQMED clone ID LmxCLK1-cb029). Positive clones were screened by colony PCR and sequenced using the primers pNUS-Fwd-Seq and pNUS-Rev-Seq.

In order to modify the pPLOT template vector for generation of CRISPR/Cas9 repair templates for transfection in *Leishmania* spp., the NLuc coding sequence was introduced by PCR amplification from NLuc-EcoDHFR using oligonucleotides NLuc-fb002 and NLuc-rb003, which contain a 5′ 20 nt overlap with the pPLOTv1 blast-mNeonGreen-blast *Hind*III/*Bam*HI linearized. Fragments were purified using PureLink™ PCR Purification Kit (Invitrogen) before Gibson Assembly (NEB) was performed, and the final construct was transformed into Mach1 *E. coli* cells. Positive clones were screened by colony PCR using the same cloning primers and sequenced using the NLUC-N-terminal primer.

### Recombinant protein production

Recombinant protein production followed a standard protocol established in our group and previously described elsewhere.^46,47^ Briefly, freshly-transformed BL21(DE3)-R3-lambda-PPase *E. coli* colonies were inoculated into 50 mL of (LB) media (containing 50 µg/mL kanamycin and 34 µg/mL chloramphenicol) and incubated at 37 °C overnight in an orbital shaker incubator (140 rpm). The resulting culture was used to inoculate 1.5 L of Terrific Broth (TB) media containing 50 µg/mL kanamycin and allowed to grow at 37 °C with aeration (140 r.p.m.) until an OD_600_ of ∼2. The culture was then transferred to an orbital shaker incubator set at 18 °C and 140 r.p.m. for 30 min before the addition of the inducing agent (Isopropyl β-D-1-thiogalactopyranoside [IPTG] at 0.2 mM final concentration). Cells were allowed to grow under these conditions for 16 h and harvested by centrifugation (15 min, 7,500 × g at 4 °C). Cell pellets were weighed and resuspended in an equivalent volume (1:1 ratio of wet weight to buffer) of 2× Binding Buffer (1× Binding Buffer was 50 mM HEPES pH 7.5, 500 mM NaCl, 10% glycerol, 10 mM imidazole, 0.5 mM tris(2-carboxyethyl)phosphine (TCEP) with protease inhibitors set II (Calbiochem, at 1:200 ratio) and frozen at - 80 °C until use.

For purification, cell pellets were thawed and sonicated on ice. Polyethyleneimine (pH 7.5) was added to cell lysates to a final concentration of 0.15% (w/v) and the samples were centrifuged at 55,000 × g for 30 min at 4 °C. The supernatants were loaded onto an immobilized metal affinity chromatography (IMAC) column (5 mL HisTrap FF Crude, GE Healthcare) and washed in Binding Buffer containing 30 mM imidazole. Proteins were eluted in Binding Buffer with 300 mM imidazole. To remove the N-terminal 6xHis-tag introduced during cloning (see above), proteins were incubated with TEV protease during dialysis overnight at 4 °C into Gel Filtration Buffer (20 mM HEPES, 500 mM NaCl, 5% glycerol, 0.5 mM TCEP). Recombinant proteins were further purified using reverse affinity chromatography on Ni-Sepharose followed by gel filtration (Superdex 200 16/60, GE Healthcare), and concentrated using 10 kDa molecular weight cut-off centrifugal concentrators (Millipore). LmxCLK1 (CQMED clone ID LmxCLK1-cb001) and NLuc-LmxCLK1 (CQMED clone ID LmxCLK1-cb021) proteins were stored at −80 °C for *in vitro* biochemical assays at 0.5 mg/mL and 0.7 mg/mL, respectively, measured by UV absorbance with a NanoDrop spectrophotometer (Thermo Scientific), using the calculated molecular weight and estimated extinction coefficient.

Purified proteins were further analyzed by LC-ESI-MS in a XEVO G2 Sx Q-ToF (Waters). Expected intact masses for LmxCLK1 (CQMED clone ID LmxCLK1-cb001) and NLuc-LmxCLK1 (CQMED clone ID LmxCLK1-cb021) were: 48,791.14 Da and 68,411.71 Da, respectively. Representative protein production results can be seen in **Supplementary Figure S2**.

### Differential scanning fluorimetry (DSF)

DSF measurements were carried out in 384-well PCR plates as described before.^47^ Briefly, recombinant LmxCLK1 (CQMED clone ID LmxCLK1-cb001) was screened against a library of 378 structurally diverse ATP-competitive kinase inhibitors available from Selleckchem (Houston, TX, United States; catalog #L1200). Each well contained 20 μL of 2 μM kinase in Gel Filtration Buffer complemented with 5x SYPRO Orange (Invitrogen). Compounds, previously solubilized in 100% DMSO, were used at 10 µM final concentration (final DMSO assay concentration was 0.1%). Plates were sealed using optically clear films and transferred to a QuantStudio 6 qPCR instrument (Applied Biosystems). The fluorescence intensity was measured over a temperature gradient from 25 to 95 °C at a constant rate of 0.05 °C/s and protein melting temperatures were calculated based on a Boltzmann function fitting to experimental data, as implemented in the Protein Thermal Shift Software (Applied Biosystems). Protein in 0.1% DMSO was used as reference for thermal shift calculations. Compounds that caused a shift in the mid-point melting temperature of the protein (ΔTm) > 2 °C compared to the reference were considered positive hits. Data shown are from two independent experiments in which single ΔTm values were acquired for each one of the tested compounds. A list of all compounds used in these experiments and ΔTm values can be found in **Supplementary Table S3**.

### Enzyme inhibition assays

We monitored the activity of recombinantly produced full-length LmxCLK1 (CQMED clone ID LmxCLK1-cb001) using a commercially-available homogenous time-resolved fluorescence (HTRF)-based assay (HTRF KinEASE-STK S3 kit, Cisbio - catalog #62ST3PEC). Assay development included establishing reaction parameters (optimal enzyme concentration, ATP concentration and reaction time). Typical assay development results can be seen in **Supplementary Figure S3**.

For IC_50_ measurements, compound stock solutions (in 100% DMSO) were serially diluted in 100% DMSO (15-point, 3-fold serial dilution and a highest final assay concentration of 10 µM) prior to transfer (100 nL) to a black, 384-well, round bottom, low-volume plate (Corning catalog #4514) containing a mixture of LmxCLK1 and peptide substrate in kinase buffer using an automated liquid transferring system (CYBIO FeliX®, Analytik Jena, Jena, Germany). Following a 30-min incubation period at 25 °C, the reaction was started by the addition of ATP and allowed to proceed for 75 min at 30 °C. Final concentrations for reaction components were: 6 nM LmxCLK1, 66 µM ATP (10× K_M,ATP_), 1 µM peptide (STK Substrate 3-biotin, Cisbio). All components were prepared in 1× Enzymatic Buffer (Cisbio) supplemented with 5 mM MgCl_2_, 1 mM DTT, 0.01% Tween-20. The reaction was stopped by the addition of Detection Buffer containing EDTA, Streptavidin-XL665 (Cisbio) and STK antibody-Eu3+-Cryptate (Cisbio) and incubated for 1 h at 25 °C. Fluorescence signal was measured using a CLARIOstar microplate reader (BMG LABTECH) with excitation at 330 nm, emission at 665 nm (acceptor) and 620 nm (donor), integration delay 60 μs and integration time 400 μs. All reactions were done in duplicate and two independent experiments were performed. Enzyme activity was expressed as the background-corrected HTRF ratio of the acceptor and donor emission signals according to equation [1]:

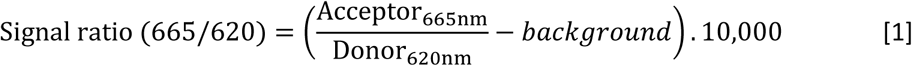

where *background* is the background fluorescence obtained in the absence of enzyme (blank).

To determine IC_50_ values, the background-corrected HTRF ratio was plotted as a function of test compound concentration, and the data were fitted to the sigmoidal dose-response (variable slope) equation [2] available in GraphPad Prism (v. 9.1.0):

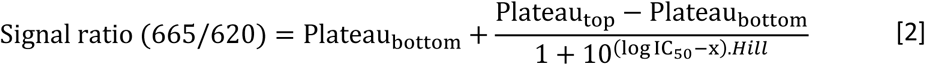

where *Hill* = Hill slope describing the steepness of the curve and x = test compound concentration.

To obtain normalized enzymatic activity we used the background corrected HTRF ratio of enzymatic reactions performed in the presence of vehicle only (DMSO) or 10 µM staurosporine as controls for 100% and 0% enzyme activity, respectively.

To establish the contribution of compound pre-incubation to the IC_50_ assay, we pre-incubated WZ8040 with LmxCLK1 for 5, 10, 15 and 30 minutes at room temperature, prior to starting the phosphorylation reaction by adding ATP and the peptide substrate. The enzymatic reactions were carried following the procedures described above.

For fitting of k_inact_ and K_I_ values from endpoint pre-incubation IC_50_ data, we used the equation [3], first described by Krippendorff and colleagues:^39^

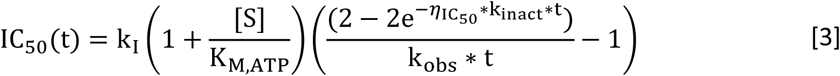

where 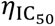 is defined by equation [4]:

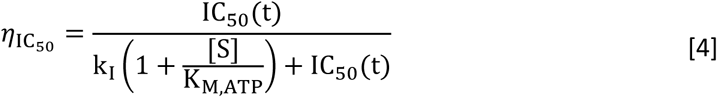

However, under the experimental conditions used in our enzymatic assay, we were unable to determine reliable values of k_inact_ and K_I_.

### Intact mass spectrometry analysis for investigation of covalent adduct formation

We used intact mass spectrometry to verify the formation of covalent adducts between purified LmxCLK1 (CQMED clone ID LmxCLK1-cb001) and the two covalent inhibitors WZ8040 and WZ3146. These experiments were performed at an inhibitor to protein ratio of 3:1 and incubation of 2 hours at room temperature. Unmodified protein (LmxCLK1-cb001) was used as control. Samples were analyzed by LC-ESI-MS in a XEVO G2 Sx Q-ToF (Waters) at LACTAD-UNICAMP, Brazil. Representative results can be seen in **Supplementary Figure S4**.

### Identification of LmxCLK1 modification by WZ8040 using LC-MS/MS

To identify LmxCLK1 residues forming covalent adducts with WZ8040, the compound was incubated with purified LmxCLK1 at an inhibitor to protein ratio of 3:1 for 2 hours at room temperature. Unmodified protein (LmxCLK1-cb001) was used as control. Triplicate samples were partially separated in SDS-PAGE. Following gel extraction, samples were reduced with DTE and alkylated with iodoacetamide before digestion with Promega sequencing grade trypsin (#V5111). Resulting extracted peptides were analyzed by LC-MS/MS.

Peptides were loaded onto an mClass nanoflow UPLC system (Waters) equipped with a nanoEaze M/Z Symmetry 100 Å C_18_, 5 µm trap column (180 µm x 20 mm, Waters) and a PepMap, 2 µm, 100 Å, C_18_ EasyNano nanocapillary column (75 µm x 500 mm, Thermo). The trap wash solvent was aqueous 0.05% (v:v) trifluoroacetic acid and the trapping flow rate was 15 µL/min. The trap was washed for 5 min before switching flow to the capillary column. Separation used gradient elution of two solvents: solvent A, aqueous 0.1% (v:v) formic acid; solvent B, acetonitrile containing 0.1% (v:v) formic acid. The flow rate for the capillary column was 300 nL/min and the column temperature was 40°C. The linear multi-step gradient profile was: 3-10% B over 7 mins, 10-35% B over 30 mins, 35-99% B over 5 mins and then proceeded to wash with 99% solvent B for 4 min. The column was returned to initial conditions and re-equilibrated for 15 min before subsequent injections.

The nanoLC system was interfaced with an Orbitrap Fusion Tribrid mass spectrometer (Thermo) with an EasyNano ionization source (Thermo). Positive ESI-MS and MS^2^ spectra were acquired using Xcalibur software (version 4.0, Thermo). Instrument source settings were: ion spray voltage, 1,900 V; sweep gas, 0 Arb; ion transfer tube temperature; 275°C. MS^1^ spectra were acquired in the Orbitrap with: 120,000 resolution, scan range: *m/z* 375-1,500; AGC target, 4e^5^; max fill time, 100 ms. Data dependent acquisition was performed in top speed mode using a 1 s cycle, selecting the most intense precursors with charge states >1. Easy-IC was used for internal calibration. Dynamic exclusion was performed for 50 s post precursor selection and a minimum threshold for fragmentation was set at 5e^3^. MS^2^ spectra were acquired in the linear ion trap with: scan rate, turbo; quadrupole isolation, 1.6 *m/z*; activation type, HCD; activation energy: 32%; AGC target, 5e^3^; first mass, 110 *m/z*; max fill time, 100 ms. Acquisitions were arranged by Xcalibur to inject ions for all available parallelizable time.

LC-MS/MS chromatograms in .raw format were imported into Progenesis QI (Version 2.2., Waters) for alignment and peak picking and a concatenated MS2 peak list was exported for database searching against expected LmxCLK1. Mascot Daemon (version 2.6.0, Matrix Science) was used to submit the search to a locally-running copy of the Mascot program (Matrix Science Ltd., version 2.7.0). Search criteria specified: Enzyme, trypsin; Max missed cleavages, 1; Variable modifications, Carbamidomethyl (C), WZ8040 (+480.150 Da), Oxidation (M); Peptide tolerance, 3 ppm; MS/MS tolerance, 0.5 Da; Instrument, ESI-TRAP. Peptide identifications were passed through the percolator algorithm to achieve a 1% false discovery rate assessed against a reverse database and individual matches filtered to require minimum expect score of 0.05. High sequence coverage (92%) was achieved for LmxCLK1 (**Supplementary Figure S5**). Four peptides (YGPCLLDWIMK, HLPPDPCR, ICDLGGCCDER, and CGTEEAR) were matched that contained the WZ8040 modification. The obtained expect scores and annotated fragmentation spectra are shown in **Supplementary Figure S6**. Peptide identifications with FDR <1% were imported into Progenesis QI, mapped between runs and filtered to the best scoring identification per assigned mass. Relative peptide quantification was extracted from integration of precursor ion areas following normalization between all runs to total peptide signal. An ANOVA was applied between the treated and non-treated samples to look for significant difference in peptide abundance between groups. The Hochberg and Banjamini multiple test correction was used to calculate q-values from the ANOVA p-values. A summary of the quantification results is shown in **Supplementary Figure S7**.

### Cell culture

*L. mexicana* (strain MNYC/BZ/62/M379) promastigotes were routinely cultivated in medium 199 (M199) (Earle’s Salts - Gibco) supplemented with 10% (v/v) heat-inactivated fetal calf serum (HIFCS – Cultilab) and 1% Penicillin/Streptomycin solution (Sigma-Aldrich) at 26 °C. Parasites were sub cultured at 2.0 × 10^5^ parasites mL^−1^, reaching mid-log phase at 48-72 h with ∼5.0 × 10^6^ parasites mL^−1^, and stationary phase at 168 h with ∼2.0 × 10^7^ parasites mL^−1^. Phenotypic drug response assays in null mutant and point-mutation mutant lines were performed in HOMEM supplemented with 10% (v/v) heat-inactivated fetal calf serum (HIFCS – Gibco).

THP-1 monocytes were differentiated in adherent macrophages with 50 ng of PMA for 72 h at 37 °C and 5% CO_2_ before infection with stationary phase promastigotes (ratio 10 promastigotes:1 macrophage) for 6 h, followed by washing to remove non-internalized *Leishmania*. Infected macrophages were incubated at 34 °C and 5% CO_2_ for 48 h in RPMI (Gibco) supplemented with 10% HIFCS and 1% Penicillin/Streptomycin solution.

For evaluation of drug effect on promastigote (2.0×10^6^ cells/mL) and uninfected macrophage (2.0×10^5^ cells/mL) viability, cells were cultivated as described above in 96-well plates in the presence of test compounds for 24 h and incubated with 10% AlamarBlue™ Cell Viability Reagent (ThermoFisher) for 24 h before measuring fluorescence at λex 540 nm/λem 590 nm. Values of untreated cells (1% DMSO) were set as 100% viable cells. For evaluation of drug effect in intracellular amastigotes, the infections were performed as describe above before addition of test compounds followed by incubation for 48 h. Cells were fixed in 4% paraformaldehyde before incubation with 5 µg/mL of DAPI (4’, 6-diamino-2-fenilindol) and slides sealed with ProLong™ Gold Antifade mounting media (ThermoFisher). Slides were imaged on a Zeiss AxioImager microscope with a ×63 lens. Multiple field of view were counted per drug concentration and DMSO control to reach a minimum of 200 macrophages, in duplicate. The total number of amastigotes was normalized by the number of macrophages counted. The EC_50_ values were determined by fitting the data to the sigmoidal dose-response (variable slope) equation from GraphPad Prism (v.9.1.0).

### Genetic manipulation of *Leishmania mexicana*

Endogenously NLuc tagged mutants were generated using a CRISPR/Cas9-based approach^48^, which consists on generating by PCR a 5’UTR guide RNA and a repair template containing the 3myc-NLuc-tag and the blasticidin resistance marker amplified from the vector, using the oligonucleotides detailed at **Supplementary Table S1**. All DNA, including the circularized overexpression vector LmxCLK1-cb029, were sterilized by ethanol precipitation prior to electroporation. Promastigotes of *L. mexicana* were transfected in 1× Tb-BSF buffer (66.67 mM Na_2_HPO_4_, 23.33 mM NaH_2_PO_4_, 5 mM KCl, 50 mM HEPES, 150µM CaCl_2_ pH 7.3)^49^ using two pulses with program X-001 in the Amaxa Nucleofector IIb (Lonza). Parasites were immediately transferred to M199 10% HIFCS and incubated overnight at 25 °C to recover. The suitable antibiotics were added to cells and cloned by serial dilution of 1 in 6, 1 in 66 and 1 in 726 in 96-well microplates. Clones were selected from most diluted plates after 7-15 days of selection (20 μg/mL Blastidicin, 50 μg/mL hygromycin B, 10 μg/mL G418).

### Western blotting

Promastigotes were re-suspended in lysis buffer (50 mM Tris-HCl, 10 mg/mL CHAPS, pH 7.4) and freshly added 10% (v/v) Protein inhibitor cocktail I (Melford), 1 mM DTT and 1 mM PMSF for protein quantification using QPro-BCA kit (Cyanagen). Samples were prepared in Laemmli buffer, boiled for 10 min at 65 °C and 20 µg per lane loaded on 12% SDS-acrylamide gel for electrophoretic run (120 V, constant). Gels were transferred to PVDF membranes using wet transfer system (Biorad) at a constant amperage of 350 mA for 2 h. Membranes were blocked with 5% skimmed milk in Tris Buffered Saline-Tween 0.05% for 1 h before overnight incubation with Rabbit anti-c-myc Antibody (Sigma-Aldrich) diluted 1:5,000 in 3% skimmed milk in TBST at 4 °C. After washing membranes, proceeded to incubation with secondary antibody anti-rabbit IgG (H + L) conjugated to HRP (Sigma-Aldrich) diluted 1:20,000 in 3% skimmed milk in TBST for 1 h at room temperature. Membranes were washed, incubated with ECL Prime (GE Healthcare) and developed in Alliance Q9 Advanced (Uvitec Cambridge).

### Determination of energy transfer probe affinity using BRET

For BRET assays, we used recombinant NLuc-LmxCLK1 (cb021) in Kinase Buffer (50 mM HEPES, 1 mM EGTA, 10 mM MgCl_2_, 2 mM DTT and 0.01% Tween-20) at a final concentration of 125 pM or *L. mexicana* engineered strain expressing NLuc-LmxCLK1 from its endogenous *loci*, either as promastigotes or intramacrophage amastigotes. Cells or purified protein were aliquoted (85 µL) onto a 96-well plate (white, flat bottom, non-binding - Greiner, catalog #655904; or TC-treated - Greiner, catalog #655083). For tracer titration, energy transfer probes (5 µL) were added to cells or purified protein as a serial dilution series (>8 points, maximum final probe concentration of 5.0 µM). This was prepared by first diluting the concentrated probe stock solution into 100% DMSO (to 100X the final assay concentration) and then in Tracer Dilution Buffer (TDB - 12.5 mM HEPES, pH 7.5; 31.25% PEG 400) to 20x the final assay concentration. The final volume of the mixture was brought to 100 µL by the addition of 10 µL of either sterile Opti-MEM™ I Reduced Serum Medium (Thermo-Scientific) or Kinase buffer for wells containing *Leishmania* or purified protein, respectively. For tracer displacement assay, test compounds were prepared as stock solutions in DMSO (100x the final assay concentration) and further diluted in Opti-MEM for working stock (10x the final assay concentration), also bringing the assay final volume to 100 µL. Final DMSO concentration in these assays was 1%. Plates were incubated for 30 min at 25 °C (purified protein and promastigotes) or 34 °C (intramacrophage amastigotes), and 50 µL of Promega cocktail containing furimazine and NLuc extracellular inhibitor (catalog #N2161) were added to each well. Light emissions at 460+10 nm (BRET donor) and at 610+20 nm (BRET acceptor) were sequentially recorded (integration time = 0.5 s, gain = 3600) using a luminometer (BMG LABTECH Clariostar or Biotek Synergy HT). Raw BRET values were calculated by dividing the acceptor luminescence by the donor luminescence. These are reported as milli-BRET (mBRET) units (mBU), according to equation [3]:

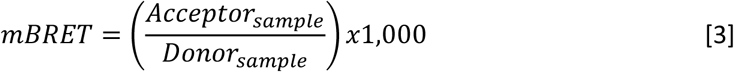

BRET values were converted to a blank-corrected probe occupancy (%) according to equation [4]:

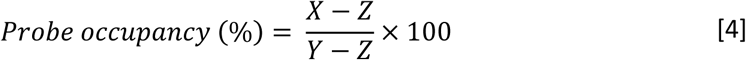

where X = mBRET value in the presence of the test compound and the probe, Y = mBRET value in the presence of probe only, and Z = mBRET value in the absence of probe and test compound.

To determine apparent dissociation constants from saturation binding curves, mBRET values were plotted as a function of probe concentration, and probe affinity values (*K*_D_) were determined using the hyperbolic dose-response equation [5] for binding to a single site available in GraphPad Prism:

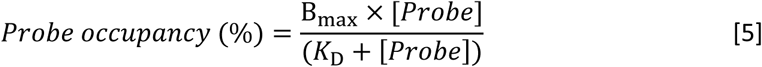

where Bmax = maximum specific binding extrapolated to very high probe concentrations.

Results for NanoBRET screening against *L. mexicana* promastigotes expressing NLuc-LmxCLK1 can be found in Supplementary Table S4.

### General Chemistry Information

All high-grade chemicals were purchased from commercial sources (Sigma-Aldrich, Combi-blocks, Thermo Fisher Scientific, Chem-Impex, Setareh Biotech, Chemodex and Lumiprobe) and used as received. The final compounds were characterized by 1H NMR and electrospray ionization-mass spectrometry (ESI-MS). NMR spectra were recorded on a Bruker 400 MHz or 500 MHz spectrometer. All 1H NMR experiments are reported in δ units, parts per million (ppm), and were measured relative to the solvent signals CDCl3 (7.26 ppm) or CD3OD (3.31 ppm). Samples were submitted for MS analysis to control the exact mass of the compound synthetized. 1μl of the sample (diluted for a final concentration of 0.1 μg/μl in 100% acetonitrile) was analyzed by reverse phase HPLC-ESI-MS using an Acquity H-class HPLC system (Waters Corp. Milford, MA, USA) which is directly connected to the XEVO G2 Sx Q-ToF (Waters) to determine the intact mass of the compound. The LC is equipped with C18 column (ACQUITY UPLC Protein BEH C18 Column, 1.7 µm, 2.1 mm X 50 mm, Waters) for small molecule separation kept at 45 °C. The mobile phase solvent A was 0.1% Formic acid (FA) in water, and solvent B was 0.1% FA in 100% Acetonitrile (ACN). Samples were loaded at a flow rate of 0.5 μl/min, and eluted from C18 column at a flow rate of 400 μl/min with one linear gradient step: one from 3 to 90% solvent B over 2.5 min. The column was regenerated by washing at 100% solvent B for 1.5 min and re-equilibrated at 1% solvent B for 3 min. Exact mass analysis was performed in positive ion electrospray in resolution mode. For internal calibration, the lockspray properties were the following: scan time was fixed at 0.5 seconds, with a mass window of 0.5 Da around Leu-enkephalin (556.2771 Da). The ToF-MS acquisition ranged from 50 Da to 2,000 Da with a scan time fixed at 0.5 second. The cone voltage on the ESI source was fixed at 40 V. MS raw data was analyzed using MassLynx software developed by Waters.

### Synthesis and characterization of GZD824-based BRET probes

All intermediates not described here were synthesized as reported by Ren and colleagues.^29^

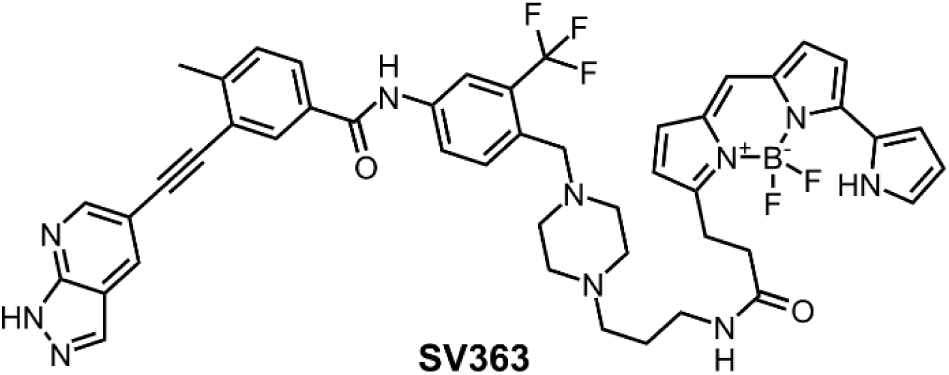

To a stirred solution of DIPEA (8.58 µL, 0.0493 mmol) in DMF (500 µL) was added the 3-((1H-pyrazolo[3,4-b]pyridin-5-yl)ethynyl)-N-(4-((4-(3-aminopropyl)piperazin-1-yl)methyl)-3-(trifluoromethyl)phenyl)-4-methylbenzamide trifluoroacetate (2.84 mg, 0.00493 mmol). The mixture was stirred for 5 min. EverFluor 590 SE (2.10 mg, 0.00493 mmol) was added and the reaction mixture was capped and stirred in the dark for 16 h under N2. The crude product was purified by flash chromatography (MeOH/DCM) to afford the desired fluorescent tracer in 80% yield. 1H NMR (500 MHz, CD3OD) δ 8.71 (d, J = 1.8 Hz, 1H), 8.45 (d, J = 1.9 Hz, 1H), 8.17 (s, 1H), 8.14 (dd, J = 9.3, 1.8 Hz, 2H), 7.94 (dd, J = 8.2, 1.6 Hz, 1H), 7.91 – 7.87 (m, 1H), 7.71 (d, J = 8.5 Hz, 1H), 7.47 (d, J = 8.1 Hz, 1H), 7.23 (s, 1H), 7.22 – 7.20 (m, 1H), 7.20 – 7.17 (m, 2H), 7.00 (d, J = 4.6 Hz, 1H), 6.92 (d, J = 3.9 Hz, 1H), 6.38 – 6.32 (m, 2H), 3.63 (s, 2H), 3.28 (m, 2H), 3.25 (t, J = 6.6 Hz, 2H), 2.69 – 2.57 (m, 9H), 2.51 (dt, J = 25.0, 13.1 Hz, 6H), 1.76 – 1.68 (m, 2H). MS (ESI): calculated for C47H45BF5N10O2 [M+H]+: 887.3740; found: 887.3777.

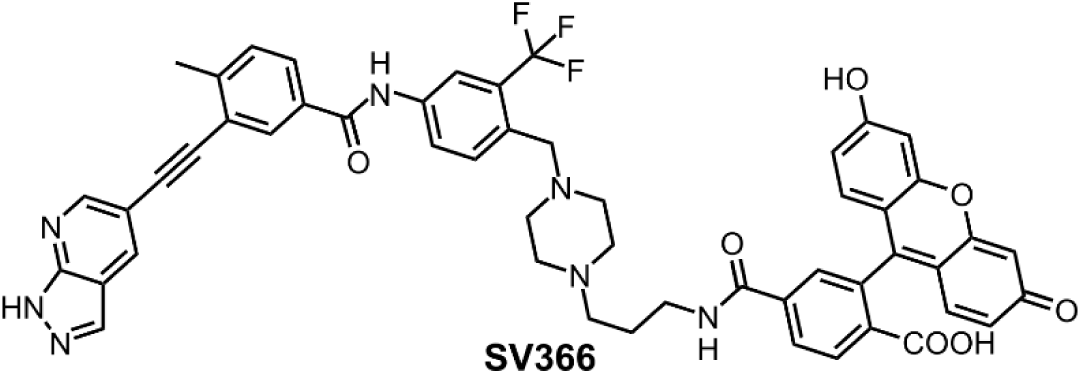

To a stirred solution of DIPEA (8.58 µL, 0.0493 mmol) in DMF (500 µL) was added 5-Carboxyfluorescein N-succinimidyl ester (2.45 mg, 0.00493 mmol). The mixture was stirred for 5 min. The 3-((1H-pyrazolo[3,4-b]pyridin-5-yl)ethynyl)-N-(4-((4-(3-aminopropyl)piperazin-1-yl)methyl)-3-(trifluoromethyl)phenyl)-4-methylbenzamide trifluoroacetate (2.84 mg, 0.00493 mmol) was added and the reaction mixture was capped and stirred in the dark for 16 h under N2. The crude product was purified by flash chromatography (MeOH/DCM) to afford the desired fluorescent tracer in 83% yield. 1H NMR (500 MHz, CD3OD) δ 8.61 (d, J = 1.5 Hz, 1H), 8.35 (td, J = 6.2, 1.6 Hz, 2H), 8.08 (dd, J = 16.7, 6.5 Hz, 3H), 7.91 (ddd, J = 10.8, 7.9, 1.7 Hz, 1H), 7.83 (ddd, J = 15.2, 8.6, 1.8 Hz, 1H), 7.78 (d, J = 7.9 Hz, 1H), 7.71 – 7.65 (m, 1H), 7.37 (d, J = 8.1 Hz, 1H), 7.23 (d, J = 7.9 Hz, 1H), 6.94 (dd, J = 9.7, 5.7 Hz, 2H), 6.51 – 6.44 (m, 4H), 3.56 (d, J = 7.7 Hz, 2H), 3.50 (s, 2H), 3.43 – 3.38 (m, 2H), 3.38 – 3.31 (m, 2H), 2.53 (s, 3H), 2.48 (dd, J = 15.0, 7.6 Hz, 4H), 2.21 – 2.14 (m, 2H), 2.09 – 2.02 (m, 2H). MS (ESI): calculated for C52H43F3N7O7 [M+H]+: 934.3176; found: 934.3206.

## Author Contributions

C.M.C.C.P. and P.Z.R. contributed equally to this work.

C.M.C.C.P. designed, executed, and analyzed the molecular biology and genetic manipulation experiments in *Leishmania*, BRET-based target engagement assays *in vitro* and in cells.

P.Z.R. designed, executed, and analyzed molecular biology and cloning experiments, recombinant protein production and enzymatic assays.

J.B.T.C. designed, executed, and analyzed the experiments with *Leishmania* T7/Cas9 knockout and cysteine-to-alanine mutant lines.

S.N.S.V. performed compound synthesis and characterization.

A.D. designed, executed, and analyzed LC-MS/MS experiments for determination of LmxCLK1 cysteines modified by WZ8040.

R.C.F. performed BRET-based target engagement assays *in vitro*.

C.V.R. assisted with molecular biology and protein production.

R.M.C., K.B.M., J.C.M., and A.C.C provided project coordination and secured funding.

C.M.C.C.P., P.Z.R. and R.M.C. wrote the manuscript.

All authors reviewed and approved the final version of the manuscript.

## Acknowledgment

We thank all members of CQMED-UNICAMP for their help and support. This study was financed, in part, by the São Paulo Research Foundation (FAPESP – grant # 2013/50724-5, 2014/50897-0 and 2016/21171-6), Embrapii (Empresa Brasileira de Pesquisa e Inovação Industrial), Promega, and CNPq (Conselho Nacional de Desenvolvimento Científico e Tecnológico – grant # 465651/2014-3 and 405235/2021-6). We thank the staff of the Life Sciences Core Facility (LaCTAD) from the State University of Campinas (UNICAMP) for genomics and proteomics analyses. P.Z.R. was a recipient of CAPES (Coordenação de Aperfeiçoamento de Pessoal de Nível Superior – 88887.136432/2017-00) and FAPESP (2024/16466-3) fellowships. R.C.F. was the recipient of a CAPES M.Sc. fellowship (130075/2020-5); S.N.S.V. was the recipient of a FAPESP post-doctoral fellowship (2018/09475-5); C.M.C.C.-P.’s fellowship was in part funded by Promega Corporation; C.V.R. was the recipient of a CAPES post-doctoral fellowship (88887.146077/2017-00). We thank our colleagues in the Bioscience Technology Facility of University of York who provided insight and expertise that greatly assisted our mass spectrometry analysis. The York Centre of Excellence in Mass Spectrometry was created thanks to a major capital investment through Science City York, supported by Yorkshire Forward with funds from the Northern Way Initiative, and subsequent support from EPSRC (EP/K039660/1; EP/M028127/1).

## Supporting Information

Supplementary Figures S1-S7 and Supplementary Tables S1-S4 (PDF).

## Notes

### Competing Interest Statement

The authors have declared no competing interest.

## References

(1) Leishmaniasis. https://www.who.int/news-room/fact-sheets/detail/leishmaniasis (accessed 2025-06-06).

(2) Visceral leishmaniasis | DNDi. https://dndi.org/diseases/visceral-leishmaniasis/ (accessed 2024-12-06).

(3) eBioMedicine. Leishmania: An Urgent Need for New Treatments. eBioMedicine 2023, 87, 104440. 10.1016/J.EBIOM.2023.104440.

(4) Baker, N.; Catta-Preta, C. M. C.; Neish, R.; Sadlova, J.; Powell, B.; Alves-Ferreira, E. V. C.; Geoghegan, V.; Carnielli, J. B. T.; Newling, K.; Hughes, C.; Vojtkova, B.; Anand, J.; Mihut, A.; Walrad, P. B.; Wilson, L. G.; Pitchford, J. W.; Volf, P.; Mottram, J. C. Systematic Functional Analysis of Leishmania Protein Kinases Identifies Regulators of Differentiation or Survival. Nat. Commun. 2021 121 2021, 12 (1), 1–15. 10.1038/s41467-021-21360-8.

(5) Burza, S.; Croft, S. L.; Boelaert, M. Leishmaniasis. Lancet (London, England) 2018, 392 (10151), 951–970. 10.1016/S0140-6736(18)31204-2.

(6) De Rycker, M.; Wyllie, S.; Horn, D.; Read, K. D.; Gilbert, I. H. Anti-Trypanosomatid Drug Discovery: Progress and Challenges. Nat. Rev. Microbiol. 2022, 21 (1), 35. 10.1038/S41579-022-00777-Y.

(7) Gilbert, I. H. Drug Discovery for Neglected Diseases: Molecular Target-Based and Phenotypic Approaches: Miniperspectives Series on Phenotypic Screening for Antiinfective Targets. J. Med. Chem. 2013, 56 (20), 7719. 10.1021/JM400362B.

(8) Wyllie, S.; Thomas, M.; Patterson, S.; Crouch, S.; De Rycker, M.; Lowe, R.; Gresham, S.; Urbaniak, M. D.; Otto, T. D.; Stojanovski, L.; Simeons, F. R. C.; Manthri, S.; MacLean, L. M.; Zuccotto, F.; Homeyer, N.; Pflaumer, H.; Boesche, M.; Sastry, L.; Connolly, P.; Albrecht, S.; Berriman, M.; Drewes, G.; Gray, D. W.; Ghidelli-Disse, S.; Dixon, S.; Fiandor, J. M.; Wyatt, P. G.; Ferguson, M. A. J.; Fairlamb, A. H.; Miles, T. J.; Read, K. D.; Gilbert, I. H. Cyclin-Dependent Kinase 12 Is a Drug Target for Visceral Leishmaniasis. Nature 2018, 560 (7717), 192–197. 10.1038/s41586-018-0356-z.

(9) Thomas, M. G.; De Rycker, M.; Ajakane, M.; Albrecht, S.; Álvarez-Pedraglio, A. I.; Boesche, M.; Brand, S.; Campbell, L.; Cantizani-Perez, J.; Cleghorn, L. A. T.; Copley, R. C. B.; Crouch, S. D.; Daugan, A.; Drewes, G.; Ferrer, S.; Ghidelli-Disse, S.; Gonzalez, S.; Gresham, S. L.; Hill, A. P.; Hindley, S. J.; Lowe, R. M.; Mackenzie, C. J.; Maclean, L.; Manthri, S.; Martin, F.; Miguel-Siles, J.; Nguyen, V. L.; Norval, S.; Osuna-Cabello, M.; Woodland, A.; Patterson, S.; Pena, I.; Quesada-Campos, M. T.; Reid, I. H.; Revill, C.; Riley, J.; Ruiz-Gomez, J. R.; Shishikura, Y.; Simeons, F. R. C.; Smith, A.; Smith, V. C.; Spinks, D.; Stojanovski, L.; Thomas, J.; Thompson, S.; Underwood, T.; Gray, D. W.; Fiandor, J. M.; Gilbert, I. H.; Wyatt, P. G.; Read, K. D.; Miles, T. J. Identification of GSK3186899/DDD853651 as a Preclinical Development Candidate for the Treatment of Visceral Leishmaniasis. J. Med. Chem. 2019, 62 (3), 1180–1202. 10.1021/ACS.JMEDCHEM.8B01218/SUPPL_FILE/JM8B01218_SI_002.CSV.

(10) Wirjanata, G.; Lin, J.; Dziekan, J. M.; El Sahili, A.; Chung, Z.; Tjia, S.; Binte Zulkifli, N. E.; Boentoro, J.; Tham, R.; Jia, L. S.; Go, K. D.; Yu, H.; Partridge, A.; Olsen, D.; Prabhu, N.; Sobota, R. M.; Nordlund, P.; Lescar, J.; Bozdech, Z. Identification of an Inhibitory Pocket in Falcilysin Provides a New Avenue for Malaria Drug Development. Cell Chem. Biol. 2024, 31 (4), 743–759.e8. 10.1016/J.CHEMBIOL.2024.03.002.

(11) Dziekan, J. M.; Yu, H.; Chen, D.; Dai, L.; Wirjanata, G.; Larsson, A.; Prabhu, N.; Sobota, R. M.; Bozdech, Z.; Nordlund, P. Identifying Purine Nucleoside Phosphorylase as the Target of Quinine Using Cellular Thermal Shift Assay. Sci. Transl. Med. 2019, 11 (473). 10.1126/SCITRANSLMED.AAU3174/SUPPL_FILE/AAU3174_SM.PDF.

(12) Giannangelo, C.; Challis, M. P.; Siddiqui, G.; Edgar, R.; Malcolm, T. R.; Webb, C. T.; Drinkwater, N.; Vinh, N.; Macraild, C.; Counihan, N.; Duffy, S.; Wittlin, S.; Devine, S. M.; Avery, V. M.; De Koning-Ward, T.; Scammells, P.; McGowan, S.; Creek, D. J. Chemoproteomics Validates Selective Targeting of Plasmodium M1 Alanyl Aminopeptidase as an Antimalarial Strategy. Elife 2024, 13. 10.7554/ELIFE.92990,.

(13) Jones, N. G.; Catta-Preta, C. M. C.; Lima, A. P. C. A.; Mottram, J. C. Genetically Validated Drug Targets in Leishmania: Current Knowledge and Future Prospects. ACS Infect. Dis. 2018, 4 (4), 467–477. 10.1021/ACSINFECDIS.7B00244/SUPPL_FILE/ID7B00244_SI_003.PDF.

(14) Robers, M. B.; Dart, M. L.; Woodroofe, C. C.; Zimprich, C. A.; Kirkland, T. A.; Machleidt, T.; Kupcho, K. R.; Levin, S.; Hartnett, J. R.; Zimmerman, K.; Niles, A. L.; Ohana, R. F.; Daniels, D. L.; Slater, M.; Wood, M. G.; Cong, M.; Cheng, Y.-Q.; Wood, K. V. Target Engagement and Drug Residence Time Can Be Observed in Living Cells with BRET. Nat. Commun. 2015, 6 (1), 10091. 10.1038/ncomms10091.

(15) Robers, M. B.; Vasta, J. D.; Corona, C. R.; Ohana, R. F.; Hurst, R.; Jhala, M. A.; Comess, K. M.; Wood, K. V. Quantitative, Real-Time Measurements of Intracellular Target Engagement Using Energy Transfer. Methods Mol. Biol. 2019, 1888, 45–71. 10.1007/978-1-4939-8891-4_3.

(16) Fanti, R. C.; Vasconcelos, S. N. S.; Catta-Preta, C. M. C.; Sullivan, J. R.; Riboldi, G. P.; Dos Reis, C. V.; Ramos, P. Z.; Edwards, A. M.; Behr, M. A.; Couñago, R. M. A Target Engagement Assay to Assess Uptake, Potency, and Retention of Antibiotics in Living Bacteria. ACS Infect. Dis. 2022, 8 (8), 1449–1467. 10.1021/ACSINFECDIS.2C00073.

(17) Akiyoshi, B.; Gull, K. Discovery of Unconventional Kinetochores in Kinetoplastids. Cell 2014, 156 (6), 1247–1258. 10.1016/j.cell.2014.01.049.

(18) Nerusheva, O. O.; Akiyoshi, B. Divergent Polo Box Domains Underpin the Unique Kinetoplastid Kinetochore. 10.1098/rsob.150206.

(19) Nerusheva, O. O.; Ludzia, P.; Akiyoshi, B. Identification of Four Unconventional Kinetoplastid Kinetochore Proteins KKT22-25 in Trypanosoma Brucei. Open Biol. 2019, 9 (12). 10.1098/RSOB.190236,.

(20) Saldivia, M.; Fang, E.; Ma, X.; Myburgh, E.; Carnielli, J. B. T.; Bower-Lepts, C.; Brown, E.; Ritchie, R.; Lakshminarayana, S. B.; Chen, Y. L.; Patra, D.; Ornelas, E.; Koh, H. X. Y.; Williams, S. L.; Supek, F.; Paape, D.; McCulloch, R.; Kaiser, M.; Barrett, M. P.; Jiricek, J.; Diagana, T. T.; Mottram, J. C.; Rao, S. P. S. Targeting the Trypanosome Kinetochore with CLK1 Protein Kinase Inhibitors. Nat. Microbiol. 2020 510 2020, 5 (10), 1207–1216. 10.1038/s41564-020-0745-6.

(21) Saldivia, M.; Wollman, A. J. M.; Carnielli, J. B. T.; Jones, N. G.; Leake, M. C.; Bower-Lepts, C.; Rao, S. P. S.; Mottram, J. C. A CLK1-KKT2 Signaling Pathway Regulating Kinetochore Assembly in Trypanosoma Brucei. MBio 2021, 12 (3), e00687–21. 10.1128/MBIO.00687-21.

(22) Geoghegan, V.; Carnielli, J. B. T.; Jones, N. G.; Saldivia, M.; Antoniou, S.; Hughes, C.; Neish, R.; Dowle, A.; Mottram, J. C. CLK1/CLK2-Driven Signalling at the Leishmania Kinetochore Is Captured by Spatially Referenced Proximity Phosphoproteomics. Commun. Biol. 2022, 5 (1). 10.1038/S42003-022-04280-1,.

(23) Abramson, J.; Adler, J.; Dunger, J.; Evans, R.; Green, T.; Pritzel, A.; Ronneberger, O.; Willmore, L.; Ballard, A. J.; Bambrick, J.; Bodenstein, S. W.; Evans, D. A.; Hung, C. C.; O’Neill, M.; Reiman, D.; Tunyasuvunakool, K.; Wu, Z.; Žemgulytė, A.; Arvaniti, E.; Beattie, C.; Bertolli, O.; Bridgland, A.; Cherepanov, A.; Congreve, M.; Cowen-Rivers, A. I.; Cowie, A.; Figurnov, M.; Fuchs, F. B.; Gladman, H.; Jain, R.; Khan, Y. A.; Low, C. M. R.; Perlin, K.; Potapenko, A.; Savy, P.; Singh, S.; Stecula, A.; Thillaisundaram, A.; Tong, C.; Yakneen, S.; Zhong, E. D.; Zielinski, M.; Žídek, A.; Bapst, V.; Kohli, P.; Jaderberg, M.; Hassabis, D.; Jumper, J. M. Accurate Structure Prediction of Biomolecular Interactions with AlphaFold 3. Nature 2024, 630 (8016), 493–500. 10.1038/S41586-024-07487-W,.

(24) Vasta, J. D.; Corona, C. R.; Wilkinson, J.; Zimprich, C. A.; Hartnett, J. R.; Ingold, M. R.; Zimmerman, K.; Machleidt, T.; Kirkland, T. A.; Huwiler, K. G.; Ohana, R. F.; Slater, M.; Otto, P.; Cong, M.; Wells, C. I.; Berger, B. T.; Hanke, T.; Glas, C.; Ding, K.; Drewry, D. H.; Huber, K. V. M.; Willson, T. M.; Knapp, S.; Müller, S.; Meisenheimer, P. L.; Fan, F.; Wood, K. V.; Robers, M. B. Quantitative, Wide-Spectrum Kinase Profiling in Live Cells for Assessing the Effect of Cellular ATP on Target Engagement. Cell Chem. Biol. 2018, 25 (2), 206–214.e11. 10.1016/J.CHEMBIOL.2017.10.010.

(25) Wells, C. I.; Vasta, J. D.; Corona, C. R.; Wilkinson, J.; Zimprich, C. A.; Ingold, M. R.; Pickett, J. E.; Drewry, D. H.; Pugh, K. M.; Schwinn, M. K.; Hwang, B. (Brian); Zegzouti, H.; Huber, K. V. M.; Cong, M.; Meisenheimer, P. L.; Willson, T. M.; Robers, M. B. Quantifying CDK Inhibitor Selectivity in Live Cells. Nat. Commun. 2020, 11 (1). 10.1038/S41467-020-16559-0.

(26) Walker, J. R.; Hall, M. P.; Zimprich, C. A.; Robers, M. B.; Duellman, S. J.; Machleidt, T.; Rodriguez, J.; Zhou, W. Highly Potent Cell-Permeable and Impermeable NanoLuc Luciferase Inhibitors. ACS Chem. Biol. 2017, 12 (4), 1028–1037. 10.1021/ACSCHEMBIO.6B01129/ASSET/IMAGES/LARGE/CB-2016-01129A_0005.JPEG.

(27) Niesen, F. H.; Berglund, H.; Vedadi, M. The Use of Differential Scanning Fluorimetry to Detect Ligand Interactions That Promote Protein Stability. Nat. Protoc. 2007, 2 (9), 2212–2221. 10.1038/NPROT.2007.321.

(28) Fedorov, O.; Niesen, F. H.; Knapp, S. Kinase Inhibitor Selectivity Profiling Using Differential Scanning Fluorimetry. In Methods in molecular biology (Clifton, N.J.); 2012; Vol. 795, pp 109–118. 10.1007/978-1-61779-337-0_7.

(29) Ren, X.; Pan, X.; Zhang, Z.; Wang, D.; Lu, X.; Li, Y.; Wen, D.; Long, H.; Luo, J.; Feng, Y.; Zhuang, X.; Zhang, F.; Liu, J.; Leng, F.; Lang, X.; Bai, Y.; She, M.; Tu, Z.; Pan, J.; Ding, K. Identification of GZD824 as an Orally Bioavailable Inhibitor That Targets Phosphorylated and Nonphosphorylated Breakpoint Cluster Region-Abelson (Bcr-Abl) Kinase and Overcomes Clinically Acquired Mutation-Induced Resistance against Imatinib. J. Med. Chem. 2013, 56 (3), 879–894. 10.1021/JM301581Y.

(30) Yung-Chi, C.; Prusoff, W. H. Relationship between the Inhibition Constant (K1) and the Concentration of Inhibitor Which Causes 50 per Cent Inhibition (I50) of an Enzymatic Reaction. Biochem. Pharmacol. 1973, 22 (23), 3099–3108. 10.1016/0006-2952(73)90196-2.

(31) O’Brien, J.; Wilson, I.; Orton, T.; Pognan, F. Investigation of the Alamar Blue (Resazurin) Fluorescent Dye for the Assessment of Mammalian Cell Cytotoxicity. Eur. J. Biochem. 2000, 267 (17), 5421–5426. 10.1046/J.1432-1327.2000.01606.X,.

(32) Martin, M. M.; Lindqvist, L. The PH Dependence of Fluorescein Fluorescence. J. Lumin. 1975, 10 (6), 381–390. 10.1016/0022-2313(75)90003-4.

(33) Sjöback, R.; Nygren, J.; Kubista, M. Absorption and Fluorescence Properties of Fluorescein. Spectrochim. Acta Part A Mol. Biomol. Spectrosc. 1995, 51 (6), L7–L21. 10.1016/0584-8539(95)01421-P.

(34) Zhou, W.; Ercan, D.; Chen, L.; Yun, C. H.; Li, D.; Capelletti, M.; Cortot, A. B.; Chirieac, L.; Iacob, R. E.; Padera, R.; Engen, J. R.; Wong, K. K.; Eck, M. J.; Gray, N. S.; Jänne, P. A. Novel Mutant-Selective EGFR Kinase Inhibitors against EGFR T790M. Nature 2009, 462 (7276), 1070–1074. 10.1038/NATURE08622.

(35) Torrance, C. J.; Jackson, P. E.; Montgomery, E.; Kinzler, K. W.; Vogelstein, B.; Wissner, A.; Nunes, M.; Frost, P.; Discafani, C. M. Combinatorial Chemoprevention of Intestinal Neoplasia. Nat. Med. 2000, 6 (9), 1024–1028. 10.1038/79534.

(36) Gehringer, M.; Borsari, C.; Augusto, R.; Serafim, M.; Tavares, E.; Alves, M.; Gomes Pernichelle, F.; Nascimento, L. A.; Ferreira, G. M.; Ferreira, E. I. Covalent Inhibitors for Neglected Diseases: An Exploration of Novel Therapeutic Options. Pharm. 2023, Vol. 16, Page 1028 2023, 16 (7), 1028. 10.3390/PH16071028.

(37) Singh, J.; Petter, R. C.; Baillie, T. A.; Whitty, A. The Resurgence of Covalent Drugs. Nat. Rev. Drug Discov. 2011 104 2011, 10 (4), 307–317. 10.1038/nrd3410.

(38) Sutanto, F.; Konstantinidou, M.; Dömling, A. Covalent Inhibitors: A Rational Approach to Drug Discovery. RSC Med. Chem. 2020, 11 (8), 876–884. 10.1039/D0MD00154F.

(39) Krippendorff, B. F.; Neuhaus, R.; Lienau, P.; Reichel, A.; Huisinga, W. Mechanism-Based Inhibition: Deriving K(I) and k(Inact) Directly from Time-Dependent IC(50) Values. J. Biomol. Screen. 2009, 14 (8), 913–923. 10.1177/1087057109336751.

(40) Mader, L. K.; Keillor, J. W. Fitting of Kinact and KI Values from Endpoint Pre-Incubation IC50 Data. ACS Med. Chem. Lett. 2024, 15 (5), 731–738. 10.1021/ACSMEDCHEMLETT.4C00054.

(41) Tan, L.; Gurbani, D.; Weisberg, E. L.; Hunter, J. C.; Li, L.; Jones, D. S.; Ficarro, S. B.; Mowafy, S.; Tam, C. P.; Rao, S.; Du, G.; Griffin, J. D.; Sorger, P. K.; Marto, J. A.; Westover, K. D.; Gray, N. S. Structure-Guided Development of Covalent TAK1 Inhibitors. Bioorg. Med. Chem. 2017, 25 (3), 838–846. 10.1016/J.BMC.2016.11.035.

(42) Rao, S.; Gurbani, D.; Du, G.; Everley, R. A.; Browne, C. M.; Chaikuad, A.; Li, T.; Schröder, M.; Gondi, S.; Ficarro, S. B.; Sim, T.; Kim, N. D.; Berberich, M. J.; Knapp, S.; Marto, J. A.; Westover, K. D.; Sorger, P. K.; Gray, N. S. Leveraging Compound Promiscuity to Identify Targetable Cysteines within the Kinome. Cell Chem. Biol. 2019. 10.1016/j.chembiol.2019.02.021.

(43) Savitsky, P.; Bray, J.; Cooper, C. D. O.; Marsden, B. D.; Mahajan, P.; Burgess-Brown, N. A.; Gileadi, O. High-Throughput Production of Human Proteins for Crystallization: The SGC Experience. J. Struct. Biol. 2010, 172 (1), 3–13. 10.1016/J.JSB.2010.06.008.

(44) Aslanidis, C.; de Jong, P. J. Ligation-Independent Cloning of PCR Products (LIC-PCR). Nucleic Acids Res. 1990, 18 (20), 6069–6074. 10.1093/nar/18.20.6069.

(45) Gileadi, O.; Burgess-Brown, N. A.; Colebrook, S. M.; Berridge, G.; Savitsky, P.; Smee, C. E. A.; Loppnau, P.; Johansson, C.; Salah, E.; Pantic, N. H. High Throughput Production of Recombinant Human Proteins for Crystallography. Methods Mol. Biol. 2008, 426, 221–246. 10.1007/978-1-60327-058-8_14.

(46) Tosarini, T. R.; Ramos, P. Z.; Profeta, G. S.; Baroni, R. M.; Massirer, K. B.; Couñago, R. M.; Mondego, J. M. C. Cloning, Expression and Purification of Kinase Domains of Cacao PR-1 Receptor-like Kinases. Protein Expr. Purif. 2018, 146, 78–84. 10.1016/j.pep.2018.01.004.

(47) Santiago, A. da S.; Couñago, R. M.; Ramos, P. Z.; Godoi, P. H. C.; Massirer, K. B.; Gileadi, O.; Elkins, J. M. Structural Analysis of Inhibitor Binding to CAMKK1 Identifies Features Necessary for Design of Specific Inhibitors. Sci. Rep. 2018, 8 (1), 2–11. 10.1038/s41598-018-33043-4.

(48) Beneke, T.; Madden, R.; Makin, L.; Valli, J.; Sunter, J.; Gluenz, E. A CRISPR Cas9 High-Throughput Genome Editing Toolkit for Kinetoplastids. R. Soc. open Sci. 2017, 4 (5), 1–16. 10.1098/RSOS.170095.

(49) Schumann Burkard, G.; Jutzi, P.; Roditi, I. Genome-Wide RNAi Screens in Bloodstream Form Trypanosomes Identify Drug Transporters. Mol. Biochem. Parasitol. 2011, 175 (1), 91–94. 10.1016/J.MOLBIOPARA.2010.09.002.

