## Supplementary Information for "Discovery and characterization of cell-permeable inhibitors of *Leishmania mexicana* CLK1 using an in-cell target engagement assay"

### Supplementary Tables

#### **Supplementary Table S1.** Information of plasmids used in this work.

| **Supplementary Table S1.** Information for plasmids used in this work. | | |
|---|---|---|
| **Plasmid** | **Features** | **Reference** |
| LmxCLK1-cb001 | LmxCLK1 (residues Met48 to Met469) in pNIC28-Bsa4, $Km^r$. This construct was used for recombinant protein production in *E. coli*. | This work |
| LmxCLK1-cb020 | LmxCLK1 (residues Met48 to Met469) tagged with NLuc N-terminal in pFN31A, $Amp^r$. This construct was used as a template for cloning the fused protein in pNIC28-Bsa4 and pNUS-6myc-G418. | This work |
| LmxCLK1-cb021 | LmxCLK1 (residues Met48 to Met469) tagged with NLuc N-terminal in pNIC28-Bsa4, $Km^r$. This construct was used for recombinant protein production in *E. coli*. | This work |
| LmxCLK1-cb029 | LmxCLK1 (residues Met48 to Met469) tagged with NLuc N-terminal in pNUS-6myc-G418, $Amp^r$. This construct was used for LmxCLK1 overexpression in *L. mexicana*, $G418^r$. | This work |
| NLuc-cb001 | Template plasmid for CRISPR-Cas9 template repair amplification of myc::NLuc-BSD or BSD-NLuc::myc. Allows endogenous expression of the tagged gene of interest and *Leishmania* selection $BSD^r$. Bacterial resistance, $Amp^r$. | This work * |
| pFN31A | Protein fusion Flexi® Vector (Promega), generate N-terminal Fusions to NLuc® Luciferase, $Amp^r$. | Promega #N1311 (GenBank KF793052.1) |
| pNIC28-Bsa4 | N-terminal $His_6$ tag, T7 promoter, TEV protease cleavage site, sites for LIC cloning, sacB gene allows negative selection, $Km^r$. pET-derived *E. coli* expression vector. | [1] (GenBank EF198106) |
| pNUS-6myc-G418 | For episomal overexpression of N-terminus myc-tagged proteins in *Leishmania*. Bacterial selection, $Amp^r$; *Leishmania* selection, $G418^r$. | [2] |
| pPLOTv1 blast-mNeonGreen-blast | Template vector for CRISPR-Cas9 template repair amplification of myc::mNG-BSD or BSD-mNG::myc, $Amp^r$. Herein used as a backbone for introduction of NLuc in place of mNeonGreen. | [3] |

* Original plasmid [3] was modified by cloning Promega NLuc® using Gibson assembly.

#### Supplementary Table S2. DNA sequence of oligonucleotides used in this work.

| **Supplementary Table S2.** DNA sequence of oligonucleotides used in this work. | |
|---|---|
| **Primer** | **Sequence 5’- 3’** |
| LmxCLK1-fb001 | TACTTCCAATCCATGTCGCGCAGCCAGAGCGA |
| LmxCLK1-rb001 | TATCCACCTTTACTGTCACATGATCGGCGGCGGGCG |
| LmxCLK1-fb002 | TAAAGCGATCGCCATGTCGCGCAGCCAGAGCGACGC |
| LmxCLK1-rb002 | CTAGGTTTAAACCATGATCGGCGGCGGGCGCA |
| NLuc_M1_LIC_Fwd | TACTTCCAATCCATGGTCTTCACACTCGAAGATTTCGTTGG |
| pLIC-forward | TGTGAGCGGATAACAATTCC |
| pLIC-reverse | AGCAGCCAACTCAGCTTCC |
| LmxCLK1-fb006 | TACTTCCAATCCATGCGCTTCAAGATCCTGTCGC |
| NLuc-fb002 | ccggttccggttctaagcttaagcttATGGTCTTCACACTCGAAGATTTCG |
| NLuc-rb003 | ccagatcctgatccggatccggatccCGCCAGAATGCGTTCGCAC |
| LmxCLK1 CRISPR_Up F | GTCAGCCGAGAATACGGTGAGAGCTCCCCGgtataatgcagacctgctgc |
| LmxCLK1 CRISPR_Up R | AGCATCAGCAGCTGCGGAAGAGGAATCCATactacccgatcctgatccag |
| LmxCLK1 CRISPR_5' sgRNA | gaaattaatacgactcactataggGTGGGCTCTGGCGGACTTGAgttttagagctagaaatagc |
| Flexi-Rev-Seq | CCTTTCGGGCTTTGTTAG |
| LmxCLK1-fb010 | agctagCCTAGGatggtcttcacactcgaagat |
| LmxCLK1-rb010 | gctagcCCTAGGcatgatcggcggcgggcg |
| LmxCLK1-fb007 | ACCTCAAGCCCGAGAACATC |
| pNUS-Fwd-Seq | CCACTTGTCAAGCGAATTCCATATGA |
| pNUS-Rev-Seq | CGTCGAAGGAGCTCTTAAAAC |

#### Supplementary Table S3. Results for DSF-based compound screening against LmxCLK1.

**Supplementary Table S3.** Results for DSF-based compound screening against LmxCLK1.

| Ligand | ΔTm (°C) | | | |
|---|---|---|---|---|
| | **Run 1** | **Run 2** | **Average** | **St. dev.** |
| GZD824 | 17.1 | 17.7 | 17.4 | 0.3 |
| WZ8040 | 14.7 | 15.0 | 14.8 | 0.1 |
| WZ3146 | 14.2 | 14.4 | 14.3 | 0.1 |
| Pelitinib (EKB-569) | 9.1 | 12.5 | 10.8 | 1.7 |
| Staurosporine | 10.3 | 10.0 | 10.1 | 0.2 |
| SNS-314 Mesylate | 3.6 | 8.3 | 5.9 | 2.4 |
| AZD7762 | 3.5 | 3.1 | 3.3 | 0.2 |
| LY2835219 | 2.2 | 2.1 | 2.2 | 0.0 |
| AT9283 | 2.2 | 2.1 | 2.2 | 0.0 |
| IKK-16 (IKK Inhibitor VII) | 1.9 | 1.6 | 1.8 | 0.1 |
| PP242 | 1.6 | 1.7 | 1.7 | 0.0 |
| JNJ-7706621 | 1.4 | 1.7 | 1.6 | 0.2 |
| Hesperadin | 1.7 | 1.3 | 1.5 | 0.2 |
| TG100-115 | 1.7 | 1.3 | 1.5 | 0.2 |
| Crizotinib (PF-02341066) | 1.4 | 1.4 | 1.4 | 0.0 |
| Ro 31-8220 Mesylate | 1.1 | 1.6 | 1.3 | 0.3 |
| PRT062607 (P505-15) | 1.3 | 1.3 | 1.3 | 0.0 |
| TG003 | 1.4 | 1.2 | 1.3 | 0.1 |
| TAK-901 | 1.5 | 1.1 | 1.3 | 0.2 |
| PF-477736 | 1.3 | 1.2 | 1.2 | 0.0 |
| AZD5363 | 1.2 | 1.1 | 1.2 | 0.0 |
| Sotrastaurin | 1.2 | 0.9 | 1.0 | 0.2 |
| MK-8745 | 1.1 | 1.0 | 1.0 | 0.1 |
| PHA-767491 | 0.7 | 1.3 | 1.0 | 0.3 |
| AZD5438 | 1.2 | 0.8 | 1.0 | 0.2 |
| Ruxolitinib (INCB018424) | 0.9 | 1.1 | 1.0 | 0.1 |
| AP26113 | 1.1 | 0.9 | 1.0 | 0.1 |
| AMG-458 | 0.9 | 1.0 | 1.0 | 0.0 |
| AT7867 | 1.2 | 0.7 | 0.9 | 0.2 |
| SB415286 | 0.7 | 1.0 | 0.9 | 0.1 |
| NVP-BSK805 2HCl | 0.9 | 0.7 | 0.8 | 0.1 |
| PP121 | 0.9 | 0.7 | 0.8 | 0.1 |
| BGT226 (NVP-BGT226) | 0.9 | 0.7 | 0.8 | 0.1 |
| Fasudil (HA-1077) HCl | 0.9 | 0.6 | 0.7 | 0.2 |
| Saracatinib (AZD0530) | 0.8 | 0.7 | 0.7 | 0.0 |
| CCT128930 | 0.8 | 0.6 | 0.7 | 0.1 |

| | | | | |
|---|---|---|---|---|
| A-674563 | 0.8 | 0.6 | 0.7 | 0.1 |
| TAK-715 | 0.5 | 0.8 | 0.7 | 0.1 |
| PHA-680632 | 0.7 | 0.6 | 0.6 | 0.0 |
| Golvatinib (E7050) | 0.5 | 0.7 | 0.6 | 0.1 |
| SGI-1776 free base | 0.7 | 0.5 | 0.6 | 0.1 |
| AZD1480 | 0.6 | 0.5 | 0.6 | 0.0 |
| PF-3758309 | 0.7 | 0.5 | 0.6 | 0.1 |
| S-Ruxolitinib (INCB018424) | 0.6 | 0.6 | 0.6 | 0.0 |
| INK 128 (MLN0128) | 0.6 | 0.5 | 0.5 | 0.0 |
| Quercetin | 0.5 | 0.6 | 0.5 | 0.1 |
| Bosutinib (SKI-606) | 0.5 | 0.6 | 0.5 | 0.0 |
| Bardoxolone Methyl | 0.6 | 0.5 | 0.5 | 0.1 |
| Palbociclib (PD-0332991) HCl | 0.6 | 0.4 | 0.5 | 0.1 |
| Axitinib | 0.6 | 0.4 | 0.5 | 0.1 |
| XL019 | 0.6 | 0.4 | 0.5 | 0.1 |
| CEP-33779 | 0.5 | 0.5 | 0.5 | 0.0 |
| AZD3463 | 0.6 | 0.3 | 0.4 | 0.1 |
| Linifanib (ABT-869) | 0.3 | 0.6 | 0.4 | 0.1 |
| SB203580 | 0.5 | 0.4 | 0.4 | 0.0 |
| AT7519 | 0.4 | 0.5 | 0.4 | 0.0 |
| BIRB 796 (Doramapimod) | 0.5 | 0.4 | 0.4 | 0.1 |
| Triciribine | 0.6 | 0.2 | 0.4 | 0.2 |
| TGX-221 | 0.5 | 0.3 | 0.4 | 0.1 |
| CP-724714 | 0.6 | 0.2 | 0.4 | 0.2 |
| KU-55933 (ATM Kinase Inhibitor) | 0.6 | 0.2 | 0.4 | 0.2 |
| KW-2449 | 0.4 | 0.4 | 0.4 | 0.0 |
| R406 | 0.4 | 0.3 | 0.4 | 0.1 |
| XL147 | -0.1 | 0.8 | 0.4 | 0.4 |
| GSK461364 | 0.6 | 0.1 | 0.4 | 0.2 |
| VE-822 | 0.6 | 0.1 | 0.4 | 0.2 |
| AZD8055 | 0.4 | 0.3 | 0.3 | 0.1 |
| LY294002 | 0.5 | 0.2 | 0.3 | 0.2 |
| CCT137690 | 0.3 | 0.4 | 0.3 | 0.0 |
| CUDC-101 | 0.4 | 0.2 | 0.3 | 0.1 |
| P276-00 | 0.4 | 0.3 | 0.3 | 0.1 |
| Dovitinib (TKI-258) | 0.5 | 0.2 | 0.3 | 0.1 |
| PHT-427 | 0.4 | 0.3 | 0.3 | 0.1 |
| AZD8330 | 0.5 | 0.2 | 0.3 | 0.1 |
| VX-745 | 0.4 | 0.2 | 0.3 | 0.1 |
| AZ 628 | 0.4 | 0.2 | 0.3 | 0.1 |
| AC480 (BMS-599626) | 0.4 | 0.2 | 0.3 | 0.1 |
| Go 6983 | 0.6 | 0.0 | 0.3 | 0.3 |
| MK-8776 (SCH 900776) | 0.5 | 0.0 | 0.3 | 0.3 |

| | | | | |
|---|---|---|---|---|
| Degrasyn (WP1130) | 0.4 | 0.2 | 0.3 | 0.1 |
| SB202190 (FHPI) | 0.2 | 0.4 | 0.3 | 0.1 |
| AZ 960 | 0.3 | 0.3 | 0.3 | 0.0 |
| Cediranib (AZD2171) | 0.5 | 0.1 | 0.3 | 0.2 |
| AEE788 (NVP-AEE788) | 0.2 | 0.3 | 0.3 | 0.0 |
| Sorafenib Tosylate | 0.3 | 0.2 | 0.3 | 0.0 |
| Quizartinib (AC220) | 0.3 | 0.2 | 0.3 | 0.1 |
| TG101348 (SAR302503) | 0.2 | 0.3 | 0.3 | 0.1 |
| PIK-75 | 0.5 | 0.1 | 0.3 | 0.2 |
| Tofacitinib (CP-690550) | 0.4 | 0.1 | 0.3 | 0.1 |
| Selumetinib (AZD6244) | 0.3 | 0.3 | 0.3 | 0.0 |
| Sunitinib Malate | 0.3 | 0.2 | 0.2 | 0.0 |
| Masitinib (AB1010) | 0.3 | 0.2 | 0.2 | 0.1 |
| Genistein | 0.2 | 0.3 | 0.2 | 0.1 |
| PD318088 | 0.4 | 0.1 | 0.2 | 0.1 |
| PF-04691502 | 0.3 | 0.1 | 0.2 | 0.1 |
| CH5132799 | 0.4 | 0.1 | 0.2 | 0.1 |
| BI-D1870 | 0.4 | 0.0 | 0.2 | 0.2 |
| HMN-214 | 0.4 | 0.1 | 0.2 | 0.1 |
| PF-04217903 | 0.1 | 0.3 | 0.2 | 0.1 |
| Losmapimod (GW856553X) | 0.4 | 0.1 | 0.2 | 0.2 |
| Nintedanib (BIBF 1120) | 0.3 | 0.2 | 0.2 | 0.0 |
| AG-1478 (Tyrphostin AG-1478) | 0.1 | 0.3 | 0.2 | 0.1 |
| BX-912 | 0.2 | 0.2 | 0.2 | 0.0 |
| GDC-0068 | 0.4 | 0.0 | 0.2 | 0.2 |
| Brivanib Alaninate (BMS-582664) | 0.3 | 0.1 | 0.2 | 0.1 |
| Butein | 0.0 | 0.4 | 0.2 | 0.2 |
| OSI-930 | 0.4 | 0.0 | 0.2 | 0.2 |
| ENMD-2076 | 0.3 | 0.1 | 0.2 | 0.1 |
| Mubritinib (TAK 165) | 0.4 | 0.0 | 0.2 | 0.2 |
| GF109203X | 0.3 | 0.1 | 0.2 | 0.1 |
| MLN8054 | 0.3 | 0.1 | 0.2 | 0.1 |
| Tyrphostin AG 1296 | 0.3 | 0.1 | 0.2 | 0.1 |
| Fostamatinib (R788) | 0.2 | 0.2 | 0.2 | 0.0 |
| Crenolanib (CP-868596) | 0.5 | -0.1 | 0.2 | 0.3 |
| TIC10 | 0.2 | 0.2 | 0.2 | 0.0 |
| Motesanib Diphosphate (AMG-706) | 0.3 | 0.1 | 0.2 | 0.1 |
| YM201636 | 0.4 | 0.0 | 0.2 | 0.2 |
| Danusertib (PHA-739358) | 0.3 | 0.1 | 0.2 | 0.1 |
| LDK378 | 0.4 | 0.0 | 0.2 | 0.2 |
| BYL719 | 0.1 | 0.2 | 0.2 | 0.0 |
| GSK1904529A | 0.2 | 0.1 | 0.2 | 0.0 |
| BMS-777607 | 0.5 | -0.2 | 0.2 | 0.3 |

| | | | | |
|---|---|---|---|---|
| Volasertib (BI 6727) | 0.3 | 0.0 | 0.2 | 0.2 |
| Asiatic Acid | 0.2 | 0.1 | 0.2 | 0.1 |
| Pacritinib (SB1518) | 0.2 | 0.1 | 0.2 | 0.0 |
| CHIR-124 | 0.3 | 0.0 | 0.2 | 0.2 |
| BX-795 | -0.1 | 0.4 | 0.2 | 0.2 |
| TPCA-1 | 0.1 | 0.2 | 0.2 | 0.1 |
| NVP-BVU972 | 0.3 | 0.0 | 0.2 | 0.1 |
| Thiazovivin | 0.3 | 0.0 | 0.2 | 0.1 |
| Lapatinib (GW-572016) Ditosylate | 0.3 | 0.0 | 0.1 | 0.1 |
| BKM120 (NVP-BKM120) | 0.1 | 0.2 | 0.1 | 0.0 |
| Imatinib Mesylate (STI571) | 0.3 | 0.0 | 0.1 | 0.1 |
| ZM 306416 | 0.2 | 0.0 | 0.1 | 0.1 |
| OSI-420 | 0.2 | 0.1 | 0.1 | 0.0 |
| GSK429286A | 0.1 | 0.2 | 0.1 | 0.0 |
| TAK-632 | 0.2 | 0.1 | 0.1 | 0.1 |
| TWS119 | 0.1 | 0.2 | 0.1 | 0.0 |
| BMS-536924 | 0.2 | 0.1 | 0.1 | 0.1 |
| GSK2636771 | 0.2 | 0.1 | 0.1 | 0.1 |
| PD184352 (CI-1040) | 0.1 | 0.1 | 0.1 | 0.0 |
| CYC116 | 0.4 | -0.1 | 0.1 | 0.2 |
| KX2-391 | 0.2 | 0.0 | 0.1 | 0.1 |
| PFK15 | 0.2 | 0.1 | 0.1 | 0.1 |
| Palomid 529 (P529) | 0.3 | -0.1 | 0.1 | 0.2 |
| PHA-665752 | 0.1 | 0.1 | 0.1 | 0.0 |
| AG-490 (Tyrphostin B42) | 0.1 | 0.2 | 0.1 | 0.0 |
| PF-562271 | 0.3 | -0.1 | 0.1 | 0.2 |
| 3-Methyladenine | 0.2 | 0.0 | 0.1 | 0.1 |
| Varlitinib | 0.2 | 0.0 | 0.1 | 0.1 |
| MK-2461 | 0.0 | 0.2 | 0.1 | 0.1 |
| SNS-032 (BMS-387032) | 0.2 | 0.0 | 0.1 | 0.1 |
| Barasertib (AZD1152-HQPA) | 0.4 | -0.1 | 0.1 | 0.2 |
| Y-27632 2HCl | 0.1 | 0.1 | 0.1 | 0.0 |
| BMS-754807 | 0.2 | 0.0 | 0.1 | 0.1 |
| Erlotinib HCl (OSI-744) | 0.1 | 0.1 | 0.1 | 0.0 |
| WHI-P154 | 0.1 | 0.1 | 0.1 | 0.0 |
| Tivozanib (AV-951) | 0.2 | 0.0 | 0.1 | 0.1 |
| JNK Inhibitor IX | 0.3 | -0.1 | 0.1 | 0.2 |
| Amuvatinib (MP-470) | 0.1 | 0.0 | 0.1 | 0.1 |
| HER2-Inhibitor-1 | 0.2 | -0.1 | 0.1 | 0.2 |
| PF-543 | 0.2 | 0.0 | 0.1 | 0.1 |
| KU-0063794 | 0.1 | 0.1 | 0.1 | 0.0 |
| Rapamycin (Sirolimus) | 0.3 | -0.1 | 0.1 | 0.2 |
| SKI II | 0.1 | 0.1 | 0.1 | 0.0 |

| | | | | |
|---|---|---|---|---|
| Skepinone-L | 0.0 | 0.1 | 0.1 | 0.1 |
| Zotarolimus(ABT-578) | 0.2 | 0.0 | 0.1 | 0.1 |
| GDC-0941 | 0.1 | 0.0 | 0.1 | 0.1 |
| Piceatannol | 0.0 | 0.1 | 0.1 | 0.0 |
| PHA-793887 | 0.1 | 0.0 | 0.1 | 0.1 |
| Fingolimod (FTY720) HCl | 0.2 | -0.1 | 0.1 | 0.1 |
| Tie2 kinase inhibitor | 0.1 | 0.0 | 0.1 | 0.1 |
| WP1066 | 0.1 | 0.0 | 0.0 | 0.1 |
| VX-702 | 0.3 | -0.2 | 0.0 | 0.2 |
| DMSO | 0.1 | 0.0 | 0.0 | 0.0 |
| IMD 0354 | 0.2 | -0.1 | 0.0 | 0.1 |
| ZM 323881 HCl | 0.1 | 0.0 | 0.0 | 0.0 |
| CUDC-907 | 0.3 | -0.2 | 0.0 | 0.3 |
| H 89 2HCl | 0.2 | -0.1 | 0.0 | 0.2 |
| Cabozantinib (XL184) | 0.3 | -0.2 | 0.0 | 0.2 |
| GSK1059615 | 0.3 | -0.2 | 0.0 | 0.2 |
| Foretinib (GSK1363089) | 0.4 | -0.4 | 0.0 | 0.4 |
| R406 (free base) | 0.0 | 0.0 | 0.0 | 0.0 |
| Dinaciclib (SCH727965) | 0.2 | -0.1 | 0.0 | 0.2 |
| BI 2536 | 0.3 | -0.2 | 0.0 | 0.2 |
| Flavopiridol HCl | 0.1 | -0.1 | 0.0 | 0.1 |
| A66 | 0.4 | -0.3 | 0.0 | 0.3 |
| WYE-125132 (WYE-132) | 0.2 | -0.1 | 0.0 | 0.1 |
| Indirubin | 0.1 | -0.1 | 0.0 | 0.1 |
| TAE226 (NVP-TAE226) | 0.2 | -0.1 | 0.0 | 0.1 |
| CP-673451 | 0.1 | 0.0 | 0.0 | 0.0 |
| AZD2014 | 0.3 | -0.3 | 0.0 | 0.3 |
| ZSTK474 | 0.0 | 0.0 | 0.0 | 0.0 |
| Pimasertib (AS-703026) | 0.2 | -0.1 | 0.0 | 0.2 |
| OSU-03012 (AR-12) | -0.1 | 0.2 | 0.0 | 0.2 |
| Nilotinib (AMN-107) | 0.0 | 0.0 | 0.0 | 0.0 |
| GDC-0980 (RG7422) | 0.1 | -0.1 | 0.0 | 0.1 |
| Ibrutinib (PCI-32765) | 0.2 | -0.2 | 0.0 | 0.2 |
| Honokiol | -0.2 | 0.2 | 0.0 | 0.2 |
| ZCL278 | 0.1 | -0.1 | 0.0 | 0.1 |
| TSU-68 (SU6668) | 0.1 | -0.1 | 0.0 | 0.1 |
| KRN 633 | 0.1 | -0.1 | 0.0 | 0.1 |
| NVP-AEW541 | 0.1 | -0.1 | 0.0 | 0.1 |
| Gefitinib (ZD1839) | 0.0 | 0.0 | 0.0 | 0.0 |
| WZ4003 | 0.1 | -0.1 | 0.0 | 0.1 |
| AZD6482 | 0.1 | -0.1 | 0.0 | 0.1 |
| Tandutinib (MLN518) | 0.1 | -0.1 | 0.0 | 0.1 |
| TCS 359 | 0.0 | 0.0 | 0.0 | 0.0 |

| | | | | |
|---|---|---|---|---|
| NU6027 | 0.1 | -0.2 | 0.0 | 0.2 |
| PP1 | 0.2 | -0.2 | 0.0 | 0.2 |
| NMS-P937 (NMS1286937) | 0.0 | 0.0 | 0.0 | 0.0 |
| SB216763 | 0.1 | -0.1 | 0.0 | 0.1 |
| PIK-93 | 0.0 | -0.1 | 0.0 | 0.0 |
| Everolimus (RAD001) | 0.2 | -0.2 | 0.0 | 0.2 |
| PP2 | 0.2 | -0.2 | 0.0 | 0.2 |
| SAR131675 | 0.0 | -0.1 | 0.0 | 0.0 |
| SAR245409 (XL765) | 0.1 | -0.2 | 0.0 | 0.2 |
| Enzastaurin (LY317615) | 0.1 | -0.2 | 0.0 | 0.1 |
| CHIR-99021 (CT99021) HCl | 0.0 | 0.0 | 0.0 | 0.0 |
| Ridaforolimus (Deforolimus) | 0.0 | 0.0 | 0.0 | 0.0 |
| CYT387 | 0.0 | -0.1 | 0.0 | 0.1 |
| PI-103 | 0.0 | 0.0 | 0.0 | 0.0 |
| CAY10505 | 0.1 | -0.1 | 0.0 | 0.1 |
| CCT129202 | -0.1 | 0.0 | 0.0 | 0.0 |
| BIX 02188 | 0.1 | -0.2 | 0.0 | 0.1 |
| PD173074 | 0.1 | -0.1 | 0.0 | 0.1 |
| Torin 2 | -0.1 | 0.0 | 0.0 | 0.0 |
| EHop-016 | -0.1 | 0.0 | 0.0 | 0.0 |
| NSC 23766 | 0.0 | -0.1 | 0.0 | 0.1 |
| TG101209 | 0.2 | -0.3 | 0.0 | 0.2 |
| Milciclib (PHA-848125) | 0.1 | -0.2 | 0.0 | 0.2 |
| MGCD-265 | 0.0 | -0.1 | -0.1 | 0.0 |
| Aurora A Inhibitor I | 0.0 | -0.1 | -0.1 | 0.1 |
| Ponatinib (AP24534) | 0.0 | -0.1 | -0.1 | 0.1 |
| TG100713 | 0.1 | -0.2 | -0.1 | 0.1 |
| AVL-292 | 0.0 | -0.2 | -0.1 | 0.1 |
| TAK-733 | 0.2 | -0.3 | -0.1 | 0.3 |
| AZD1080 | 0.1 | -0.2 | -0.1 | 0.1 |
| Tofacitinib (CP-690550) Citrate | 0.0 | -0.2 | -0.1 | 0.1 |
| Sorafenib | -0.3 | 0.2 | -0.1 | 0.2 |
| Ro3280 | 0.1 | -0.2 | -0.1 | 0.1 |
| ETP-46464 | 0.3 | -0.4 | -0.1 | 0.3 |
| 6H05 | 0.1 | -0.2 | -0.1 | 0.1 |
| NU7441 (KU-57788) | 0.0 | -0.2 | -0.1 | 0.1 |
| Pazopanib HCl (GW786034 HCl) | -0.1 | 0.0 | -0.1 | 0.0 |
| A-769662 | -0.1 | 0.0 | -0.1 | 0.0 |
| Vandetanib (ZD6474) | 0.0 | -0.1 | -0.1 | 0.0 |
| PIK-293 | 0.1 | -0.2 | -0.1 | 0.1 |
| Filgotinib (GLPG0634) | 0.2 | -0.4 | -0.1 | 0.3 |
| SSR128129E | -0.1 | -0.1 | -0.1 | 0.0 |
| ZM 447439 | 0.0 | -0.2 | -0.1 | 0.1 |

| | | | | |
|---|---|---|---|---|
| GSK690693 | 0.1 | -0.3 | -0.1 | 0.2 |
| LDC000067 | -0.1 | 0.0 | -0.1 | 0.0 |
| PF-4708671 | 0.1 | -0.2 | -0.1 | 0.2 |
| Alisertib (MLN8237) | 0.1 | -0.3 | -0.1 | 0.2 |
| Semaxanib (SU5416) | 0.1 | -0.2 | -0.1 | 0.2 |
| NVP-ADW742 | 0.0 | -0.2 | -0.1 | 0.1 |
| PD98059 | 0.0 | -0.2 | -0.1 | 0.1 |
| AR-A014418 | 0.0 | -0.1 | -0.1 | 0.0 |
| PQ 401 | 0.0 | -0.2 | -0.1 | 0.1 |
| MK-2206 2HCl | -0.1 | -0.1 | -0.1 | 0.0 |
| GDC-0879 | 0.0 | -0.2 | -0.1 | 0.1 |
| CA75 (GAK) | -0.2 | 0.0 | -0.1 | 0.1 |
| AG-1024 | 0.0 | -0.2 | -0.1 | 0.1 |
| RKI-1447 | -0.1 | -0.1 | -0.1 | 0.0 |
| SP600125 | 0.1 | -0.3 | -0.1 | 0.2 |
| CZC24832 | 0.0 | -0.2 | -0.1 | 0.1 |
| Roscovitine (Seliciclib) | 0.0 | -0.2 | -0.1 | 0.1 |
| Dabrafenib (GSK2118436) | -0.2 | 0.0 | -0.1 | 0.1 |
| EHT 1864 | 0.1 | -0.3 | -0.1 | 0.2 |
| Regorafenib (BAY 73-4506) | 0.1 | -0.3 | -0.1 | 0.2 |
| Ki8751 | 0.0 | -0.2 | -0.1 | 0.1 |
| WYE-354 | -0.1 | -0.2 | -0.1 | 0.1 |
| Cabozantinib malate (XL184) | 0.1 | -0.3 | -0.1 | 0.2 |
| Vatalanib (PTK787) 2HCl | -0.2 | 0.0 | -0.1 | 0.1 |
| VS-5584 (SB2343) | 0.1 | -0.4 | -0.1 | 0.3 |
| U0126-EtOH | -0.4 | 0.1 | -0.1 | 0.3 |
| Chrysophanic Acid | 0.0 | -0.2 | -0.1 | 0.1 |
| PD0325901 | -0.2 | -0.1 | -0.1 | 0.0 |
| GDC-0349 | -0.1 | -0.1 | -0.1 | 0.0 |
| AZD1208 | 0.1 | -0.4 | -0.1 | 0.2 |
| NVP-BHG712 | 0.1 | -0.3 | -0.1 | 0.2 |
| PF-573228 | 0.1 | -0.3 | -0.1 | 0.2 |
| KU-60019 | -0.1 | -0.2 | -0.1 | 0.1 |
| GSK2334470 | 0.3 | -0.5 | -0.1 | 0.4 |
| SGX-523 | -0.1 | -0.2 | -0.1 | 0.0 |
| PD173955 | 0.0 | -0.3 | -0.1 | 0.1 |
| MK-5108 (VX-689) | -0.1 | -0.2 | -0.1 | 0.0 |
| Vemurafenib (PLX4032) | -0.1 | -0.1 | -0.1 | 0.0 |
| GNE-0877 | -0.1 | -0.2 | -0.1 | 0.0 |
| CX-6258 HCl | 0.1 | -0.4 | -0.1 | 0.2 |
| CO-1686 (AVL-301) | 0.0 | -0.3 | -0.1 | 0.2 |
| JNK-IN-8 | 0.2 | -0.5 | -0.1 | 0.4 |
| Dasatinib | -0.1 | -0.2 | -0.1 | 0.0 |

| | | | | |
|---|---|---|---|---|
| K-Ras(G12C) inhibitor 9 | 0.3 | -0.6 | -0.1 | 0.4 |
| AZD8931 (Sapitinib) | -0.1 | -0.2 | -0.1 | 0.1 |
| PF-00562271 | 0.0 | -0.3 | -0.1 | 0.1 |
| Wortmannin | 0.0 | -0.3 | -0.2 | 0.1 |
| OSI-027 | 0.1 | -0.4 | -0.2 | 0.3 |
| OSI-906 (Linsitinib) | 0.0 | -0.3 | -0.2 | 0.1 |
| BMS-345541 | -0.1 | -0.3 | -0.2 | 0.1 |
| AZD9291 | 0.0 | -0.3 | -0.2 | 0.2 |
| Tyrphostin AG 879 | -0.1 | -0.2 | -0.2 | 0.1 |
| WAY-600 | 0.0 | -0.4 | -0.2 | 0.2 |
| SU11274 | -0.1 | -0.2 | -0.2 | 0.1 |
| PH-797804 | -0.1 | -0.3 | -0.2 | 0.1 |
| CNX-774 | -0.3 | 0.0 | -0.2 | 0.1 |
| LY2603618 | 0.0 | -0.4 | -0.2 | 0.2 |
| ZM 39923 HCl | 0.0 | -0.3 | -0.2 | 0.2 |
| Pazopanib | 0.1 | -0.4 | -0.2 | 0.3 |
| CGI1746 | 0.0 | -0.4 | -0.2 | 0.2 |
| R547 | 0.0 | -0.4 | -0.2 | 0.2 |
| PLX-4720 | -0.3 | -0.1 | -0.2 | 0.1 |
| Imatinib (STI571) | 0.0 | -0.4 | -0.2 | 0.2 |
| SMI-4a | -0.3 | -0.1 | -0.2 | 0.1 |
| CAL-101 (Idelalisib) | -0.2 | -0.2 | -0.2 | 0.0 |
| SC-514 | -0.1 | -0.3 | -0.2 | 0.1 |
| Temsirolimus (CCI-779) | -0.1 | -0.3 | -0.2 | 0.1 |
| Phenformin HCl | 0.0 | -0.4 | -0.2 | 0.2 |
| GSK1838705A | -0.1 | -0.3 | -0.2 | 0.1 |
| BMS-265246 | -0.1 | -0.3 | -0.2 | 0.1 |
| WZ4002 | -0.1 | -0.3 | -0.2 | 0.1 |
| Acadesine | -0.2 | -0.2 | -0.2 | 0.0 |
| RAF265 (CHIR-265) | -0.1 | -0.3 | -0.2 | 0.1 |
| SL-327 | -0.2 | -0.2 | -0.2 | 0.0 |
| GW5074 | 0.0 | -0.4 | -0.2 | 0.2 |
| PD168393 | 0.3 | -0.7 | -0.2 | 0.5 |
| AZD4547 | -0.1 | -0.3 | -0.2 | 0.1 |
| UNC-AA-1-0013 (AAK1) | -0.2 | -0.3 | -0.2 | 0.1 |
| KN-93 Phosphate | 0.1 | -0.5 | -0.2 | 0.3 |
| AST-1306 | -0.1 | -0.4 | -0.2 | 0.1 |
| Trametinib (GSK1120212) | 0.0 | -0.5 | -0.2 | 0.2 |
| Afatinib (BIBW2992) | 0.1 | -0.5 | -0.2 | 0.3 |
| BIO | -0.1 | -0.4 | -0.2 | 0.1 |
| SB590885 | -0.1 | -0.4 | -0.2 | 0.1 |
| PIK-294 | 0.0 | -0.5 | -0.2 | 0.2 |
| IM-12 | -0.2 | -0.3 | -0.2 | 0.1 |

| | | | | |
|---|---|---|---|---|
| GNF-2 | -0.2 | -0.3 | -0.2 | 0.1 |
| Icotinib | -0.3 | -0.2 | -0.3 | 0.0 |
| BIX 02189 | -0.2 | -0.4 | -0.3 | 0.1 |
| Rigosertib (ON-01910) | -0.2 | -0.4 | -0.3 | 0.1 |
| BS-181 HCl | -0.3 | -0.2 | -0.3 | 0.1 |
| Lenvatinib (E7080) | -0.1 | -0.4 | -0.3 | 0.2 |
| BMS-794833 | -0.1 | -0.4 | -0.3 | 0.2 |
| JNJ-38877605 | -0.1 | -0.4 | -0.3 | 0.1 |
| GNF-5 | -0.2 | -0.3 | -0.3 | 0.0 |
| GSK2126458 (GSK458) | -0.2 | -0.3 | -0.3 | 0.0 |
| CGK 733 | -0.6 | 0.0 | -0.3 | 0.3 |
| AZD2858 | -0.1 | -0.4 | -0.3 | 0.1 |
| CEP-32496 | -0.4 | -0.2 | -0.3 | 0.1 |
| DCC-2036 (Rebastinib) | -0.2 | -0.4 | -0.3 | 0.1 |
| Apatinib | -0.2 | -0.4 | -0.3 | 0.1 |
| MEK162 (ARRY-162) | -0.2 | -0.4 | -0.3 | 0.1 |
| Tyrphostin 9 | -0.3 | -0.3 | -0.3 | 0.0 |
| BAY 11-7082 | -0.4 | -0.2 | -0.3 | 0.1 |
| LP-935509 (AAK1) | -0.2 | -0.4 | -0.3 | 0.1 |
| GNE-7915 | -0.1 | -0.5 | -0.3 | 0.2 |
| ZM 336372 | -0.1 | -0.5 | -0.3 | 0.2 |
| WH-4-023 | -0.4 | -0.2 | -0.3 | 0.1 |
| LY2228820 | -0.3 | -0.3 | -0.3 | 0.0 |
| TAK-285 | -0.5 | -0.2 | -0.3 | 0.2 |
| KN-62 | -0.4 | -0.2 | -0.3 | 0.1 |
| VX-680 (Tozasertib) | -0.3 | -0.4 | -0.3 | 0.0 |
| GSK650394 | -0.4 | -0.3 | -0.3 | 0.0 |
| XMD8-92 | -0.2 | -0.5 | -0.3 | 0.1 |
| AZ20 | -0.2 | -0.5 | -0.3 | 0.2 |
| LY2784544 | -0.3 | -0.4 | -0.4 | 0.1 |
| GNE-9605 | -0.2 | -0.5 | -0.4 | 0.1 |
| Telatinib | -0.3 | -0.5 | -0.4 | 0.1 |
| VE-821 | -0.2 | -0.5 | -0.4 | 0.1 |
| Dovitinib (TKI-258) Dilactic Acid | -0.5 | -0.4 | -0.4 | 0.0 |
| Dacomitinib (PF299804) | -0.4 | -0.5 | -0.4 | 0.0 |
| AS-252424 | -0.3 | -0.6 | -0.5 | 0.2 |
| Brivanib (BMS-540215) | -0.7 | -0.2 | -0.5 | 0.2 |
| AS-604850 | -0.6 | -0.4 | -0.5 | 0.1 |
| Refametinib (RDEA119) | -0.7 | -0.3 | -0.5 | 0.2 |
| HS-173 | 0.3 | -1.5 | -0.6 | 0.9 |
| 10058-F4 | -0.6 | -0.5 | -0.6 | 0.0 |
| AMG-900 | -0.6 | -0.6 | -0.6 | 0.0 |
| IPA-3 | -1.1 | -0.1 | -0.6 | 0.5 |

| | | | | |
|---|---|---|---|---|
| AG-18 | -0.7 | -0.6 | -0.7 | 0.1 |
| CNX-2006 | -0.7 | -1.3 | -1.0 | 0.3 |
| IPI-145 (INK1197) | -1.2 | -0.9 | -1.0 | 0.1 |

#### Supplementary Table S4. Results for NanoBRET screening against *Leishmania* promastigotes.

**Supplementary Table S4.** Results for NanoBRET screening against *Leishmania* promastigotes.

| Ligand | % Occupancy | | | |
|---|---|---|---|---|
| | Replicate1 | Replicate2 | Mean | S.D. |
| WZ8040 | 60.3 | 61.4 | 60.8 | 0.8 |
| GZD824 | 48.7 | 44.3 | 46.5 | 3.2 |
| WZ3146 | 32.9 | 37.5 | 35.2 | 3.3 |
| Linifanib (ABT-869) | 24.4 | 29.1 | 26.7 | 3.4 |
| Staurosporine | 24.0 | 26.9 | 25.4 | 2.0 |
| CCT128930 | 27.0 | 22.9 | 24.9 | 2.9 |
| MK-8745 | 10.9 | 35.6 | 23.2 | 17.5 |
| CUDC-101 | 19.4 | 25.7 | 22.5 | 4.5 |
| TG101348 (SAR302503) | 21.1 | 18.1 | 19.6 | 2.1 |
| PIK-93 | 12.4 | 26.2 | 19.3 | 9.7 |
| CNX-2006 | 16.8 | 21.5 | 19.1 | 3.3 |
| Nilotinib (AMN-107) | 19.7 | 18.1 | 18.9 | 1.2 |
| Volasertib (BI 6727) | 15.6 | 22.1 | 18.9 | 4.5 |
| PHA-793887 | 11.0 | 26.0 | 18.5 | 10.5 |
| AT7867 | 9.8 | 26.9 | 18.4 | 12.1 |
| AZ 628 | 13.6 | 22.9 | 18.2 | 6.5 |
| Ponatinib (AP24534) | 11.9 | 24.4 | 18.2 | 8.8 |
| AEE788 (NVP-AEE788) | 13.2 | 22.3 | 17.7 | 6.5 |
| TG101209 | 11.9 | 22.5 | 17.2 | 7.5 |
| Pazopanib HCl (GW786034 HCl) | 19.3 | 15.2 | 17.2 | 2.9 |
| Motesanib Diphosphate (AMG-706) | 18.6 | 15.6 | 17.1 | 2.1 |
| Quercetin | 14.5 | 19.4 | 17.0 | 3.5 |
| MK-2206 2HCl | 10.2 | 23.4 | 16.8 | 9.4 |
| Tandutinib (MLN518) | 11.4 | 21.7 | 16.6 | 7.3 |
| ENMD-2076 | 19.2 | 13.8 | 16.5 | 3.8 |
| Danusertib (PHA-739358) | 15.0 | 18.0 | 16.5 | 2.1 |
| BI 2536 | 12.9 | 19.9 | 16.4 | 5.0 |
| Gefitinib (ZD1839) | 9.1 | 23.6 | 16.4 | 10.2 |
| CYC116 | 4.6 | 28.0 | 16.3 | 16.6 |
| NSC 23766 | 19.0 | 13.5 | 16.3 | 3.9 |
| GSK461364 | 8.9 | 23.6 | 16.2 | 10.4 |
| PD173074 | 1.7 | 30.5 | 16.1 | 20.4 |
| SGI-1776 free base | 10.2 | 21.9 | 16.0 | 8.3 |
| CUDC-907 | 15.6 | 16.0 | 15.8 | 0.3 |
| Crenolanib (CP-868596) | 13.5 | 17.9 | 15.7 | 3.1 |

| | | | | |
|---|---|---|---|---|
| Crizotinib (PF-02341066) | 10.7 | 20.0 | 15.3 | 6.6 |
| TG100-115 | 9.3 | 21.1 | 15.2 | 8.3 |
| CHIR-124 | 16.2 | 13.4 | 14.8 | 2.0 |
| MEK162 (ARRY-162) | 13.0 | 16.4 | 14.7 | 2.4 |
| PD318088 | 4.3 | 25.0 | 14.7 | 14.6 |
| H 89 2HCl | 11.4 | 17.6 | 14.5 | 4.4 |
| SNS-032 (BMS-387032) | 5.6 | 23.1 | 14.3 | 12.4 |
| PHA-680632 | 8.8 | 19.7 | 14.3 | 7.7 |
| WYE-125132 (WYE-132) | 14.3 | 14.2 | 14.2 | 0.0 |
| AT9283 | 17.9 | 9.6 | 13.8 | 5.9 |
| PH-797804 | 6.7 | 20.7 | 13.7 | 9.9 |
| SNS-314 Mesylate | 8.3 | 19.1 | 13.7 | 7.7 |
| ZSTK474 | 13.5 | 13.7 | 13.6 | 0.1 |
| Afatinib (BIBW2992) | 8.2 | 18.7 | 13.5 | 7.4 |
| Axitinib | 7.6 | 18.8 | 13.2 | 7.9 |
| Amuvatinib (MP-470) | 21.6 | 4.6 | 13.1 | 12.0 |
| NVP-BHG712 | 2.9 | 23.1 | 13.0 | 14.3 |
| AC480 (BMS-599626) | 19.9 | 5.9 | 12.9 | 9.8 |
| AVL-292 | 9.7 | 15.9 | 12.8 | 4.4 |
| JNK-IN-8 | 10.9 | 14.4 | 12.6 | 2.5 |
| ZM 447439 | 9.5 | 15.7 | 12.6 | 4.4 |
| Vandetanib (ZD6474) | 0.0 | 25.2 | 12.6 | 17.8 |
| AMG-900 | 5.9 | 19.2 | 12.6 | 9.4 |
| SAR245409 (XL765) | 17.8 | 7.2 | 12.5 | 7.5 |
| Losmapimod (GW856553X) | 15.0 | 9.8 | 12.4 | 3.6 |
| WH-4-023 | 6.4 | 18.4 | 12.4 | 8.5 |
| PF-04217903 | 8.6 | 16.1 | 12.4 | 5.3 |
| OSU-03012 (AR-12) | 9.4 | 14.9 | 12.2 | 3.9 |
| SAR131675 | -4.6 | 29.0 | 12.2 | 23.8 |
| KU-60019 | 7.1 | 17.2 | 12.2 | 7.1 |
| Lapatinib (GW-572016) Ditosylate | 19.6 | 4.5 | 12.0 | 10.7 |
| Zotarolimus(ABT-578) | 13.4 | 10.6 | 12.0 | 2.0 |
| BMS-536924 | 20.0 | 3.8 | 11.9 | 11.4 |
| MGCD-265 | 9.4 | 14.2 | 11.8 | 3.4 |
| GDC-0879 | 17.5 | 5.7 | 11.6 | 8.3 |
| AZD6482 | 4.8 | 18.4 | 11.6 | 9.6 |
| Selumetinib (AZD6244) | 10.4 | 12.8 | 11.6 | 1.7 |
| NVP-ADW742 | 6.4 | 16.5 | 11.4 | 7.1 |
| Barasertib (AZD1152-HQPA) | 4.1 | 18.8 | 11.4 | 10.4 |
| LY2228820 | 14.5 | 8.4 | 11.4 | 4.3 |
| Piceatannol | 24.9 | -2.1 | 11.4 | 19.1 |
| HMN-214 | 14.0 | 8.7 | 11.4 | 3.7 |
| Telatinib | -0.2 | 23.0 | 11.4 | 16.4 |

| | | | | |
|---|---|---|---|---|
| SB590885 | 15.4 | 7.3 | 11.4 | 5.7 |
| PF-543 | 14.4 | 8.1 | 11.3 | 4.5 |
| LDK378 | 14.6 | 7.7 | 11.2 | 4.8 |
| AZD7762 | 12.0 | 10.3 | 11.2 | 1.2 |
| CP-724714 | 11.9 | 10.2 | 11.1 | 1.2 |
| AZD8931 (Sapitinib) | -4.8 | 26.8 | 11.0 | 22.3 |
| CGI1746 | 8.6 | 13.5 | 11.0 | 3.4 |
| Ruxolitinib (INCB018424) | 8.4 | 13.5 | 10.9 | 3.6 |
| CEP-33779 | 20.4 | 1.4 | 10.9 | 13.4 |
| BIRB 796 (Doramapimod) | 0.1 | 21.7 | 10.9 | 15.3 |
| Tofacitinib (CP-690550) | 1.0 | 20.7 | 10.9 | 14.0 |
| Sorafenib Tosylate | 0.0 | 21.7 | 10.8 | 15.3 |
| CZC24832 | 6.6 | 14.9 | 10.8 | 5.9 |
| Foretinib (GSK1363089) | 9.3 | 12.0 | 10.7 | 1.9 |
| PLX-4720 | 9.3 | 12.0 | 10.6 | 1.9 |
| CP-673451 | 1.3 | 19.8 | 10.6 | 13.1 |
| OSI-906 (Linsitinib) | 11.7 | 9.5 | 10.6 | 1.5 |
| Lenvatinib (E7080) | 5.8 | 15.1 | 10.4 | 6.5 |
| CCT137690 | 0.1 | 20.7 | 10.4 | 14.6 |
| Milciclib (PHA-848125) | 12.3 | 8.4 | 10.3 | 2.8 |
| AZD5438 | 12.2 | 8.4 | 10.3 | 2.7 |
| AZ20 | 15.7 | 4.7 | 10.2 | 7.8 |
| R547 | 13.3 | 6.9 | 10.1 | 4.6 |
| SP600125 | 4.3 | 15.9 | 10.1 | 8.2 |
| IPA-3 | 13.4 | 6.7 | 10.1 | 4.8 |
| PF-573228 | 12.5 | 7.4 | 10.0 | 3.6 |
| DCC-2036 (Rebastinib) | 6.6 | 13.1 | 9.9 | 4.5 |
| BMS-754807 | 6.7 | 13.0 | 9.9 | 4.5 |
| Tivozanib (AV-951) | 4.2 | 15.2 | 9.7 | 7.8 |
| TWS119 | 8.4 | 10.9 | 9.7 | 1.8 |
| Indirubin | 15.7 | 3.3 | 9.5 | 8.8 |
| AZ 960 | 7.2 | 11.8 | 9.5 | 3.2 |
| VX-680 (Tozasertib) | 9.4 | 9.6 | 9.5 | 0.1 |
| Aurora A Inhibitor I | 3.5 | 15.4 | 9.4 | 8.4 |
| Brivanib (BMS-540215) | 2.2 | 16.5 | 9.4 | 10.1 |
| GSK2334470 | 9.4 | 9.3 | 9.4 | 0.1 |
| NU7441 (KU-57788) | 14.7 | 3.9 | 9.3 | 7.6 |
| LY2784544 | 6.2 | 12.4 | 9.3 | 4.4 |
| KN-62 | 5.1 | 13.4 | 9.3 | 5.9 |
| SKI II | 16.9 | 1.6 | 9.2 | 10.8 |
| Vemurafenib (PLX4032) | 6.6 | 11.8 | 9.2 | 3.7 |
| Triciribine | 9.0 | 9.4 | 9.2 | 0.3 |
| Y-27632 2HCl | 4.7 | 13.5 | 9.1 | 6.3 |

| | | | | |
|---|---|---|---|---|
| SB203580 | 12.9 | 5.3 | 9.1 | 5.3 |
| MK-8776 (SCH 900776) | 8.4 | 9.8 | 9.1 | 1.0 |
| NVP-BSK805 2HCl | 21.6 | -3.5 | 9.0 | 17.8 |
| Pazopanib | 9.0 | 9.0 | 9.0 | 0.0 |
| YM201636 | 6.1 | 11.8 | 9.0 | 4.0 |
| Trametinib (GSK1120212) | 2.0 | 15.9 | 9.0 | 9.9 |
| AZD8055 | 9.7 | 8.1 | 8.9 | 1.2 |
| PF-00562271 | 9.1 | 8.6 | 8.9 | 0.4 |
| HS-173 | 15.5 | 2.2 | 8.9 | 9.3 |
| AZD9291 | 15.9 | 1.7 | 8.8 | 10.0 |
| TAK-715 | 8.0 | 9.7 | 8.8 | 1.2 |
| Rapamycin (Sirolimus) | 11.9 | 5.5 | 8.7 | 4.5 |
| GSK429286A | 3.0 | 14.3 | 8.6 | 8.0 |
| BI-D1870 | 6.1 | 11.1 | 8.6 | 3.6 |
| VX-702 | 6.4 | 10.8 | 8.6 | 3.2 |
| BMS-794833 | 8.6 | 8.5 | 8.6 | 0.1 |
| Saracatinib (AZD0530) | 12.1 | 4.9 | 8.5 | 5.1 |
| Alisertib (MLN8237) | -0.7 | 17.6 | 8.4 | 12.9 |
| WZ4003 | -4.0 | 20.8 | 8.4 | 17.6 |
| Acadesine | 8.1 | 8.6 | 8.3 | 0.4 |
| TAK-285 | 10.1 | 6.5 | 8.3 | 2.6 |
| RAF265 (CHIR-265) | 8.0 | 8.4 | 8.2 | 0.3 |
| KW-2449 | 1.4 | 14.6 | 8.0 | 9.3 |
| HER2-Inhibitor-1 | 11.1 | 4.8 | 8.0 | 4.4 |
| GSK1838705A | -3.6 | 19.2 | 7.8 | 16.1 |
| Brivanib Alaninate (BMS-582664) | -3.4 | 19.0 | 7.8 | 15.9 |
| PIK-75 | 1.6 | 14.0 | 7.8 | 8.8 |
| LP-935509 (AAK1) | 9.1 | 6.4 | 7.7 | 1.9 |
| R406 (free base) | 0.7 | 14.7 | 7.7 | 9.9 |
| Genistein | 8.3 | 6.9 | 7.6 | 0.9 |
| EHop-016 | 12.1 | 3.1 | 7.6 | 6.4 |
| Ibrutinib (PCI-32765) | -5.9 | 21.0 | 7.6 | 19.0 |
| Bardoxolone Methyl | -1.0 | 15.9 | 7.5 | 11.9 |
| BKM120 (NVP-BKM120) | 8.7 | 6.2 | 7.4 | 1.8 |
| AS-252424 | 2.9 | 11.5 | 7.2 | 6.1 |
| Vatalanib (PTK787) 2HCl | 4.3 | 10.1 | 7.2 | 4.1 |
| BMS-777607 | 11.4 | 2.9 | 7.2 | 6.0 |
| JNK Inhibitor IX | 4.4 | 10.0 | 7.2 | 4.0 |
| ZM 306416 | 8.4 | 5.7 | 7.1 | 1.9 |
| BX-912 | 8.7 | 5.5 | 7.1 | 2.3 |
| Dovitinib (TKI-258) Dilactic Acid | 5.8 | 8.2 | 7.0 | 1.7 |
| Pacritinib (SB1518) | 10.8 | 2.8 | 6.8 | 5.6 |
| Cediranib (AZD2171) | 19.7 | -6.1 | 6.8 | 18.3 |

| | | | | |
|---|---|---|---|---|
| PRT062607 (P505-15) | -1.7 | 15.2 | 6.7 | 12.0 |
| CH5132799 | 9.5 | 3.8 | 6.6 | 4.1 |
| Tyrphostin AG 879 | 8.7 | 4.5 | 6.6 | 3.0 |
| GSK690693 | 8.0 | 5.2 | 6.6 | 1.9 |
| TAK-632 | 9.1 | 4.0 | 6.6 | 3.6 |
| GSK2126458 (GSK458) | 6.7 | 6.5 | 6.6 | 0.1 |
| Torin 2 | 11.0 | 2.1 | 6.6 | 6.3 |
| AZD1480 | 4.1 | 9.0 | 6.5 | 3.4 |
| PD173955 | 3.3 | 9.7 | 6.5 | 4.6 |
| PP242 | 2.7 | 10.1 | 6.4 | 5.2 |
| 10058-F4 | -2.1 | 14.9 | 6.4 | 12.0 |
| BS-181 HCl | 4.0 | 8.7 | 6.3 | 3.3 |
| NU6027 | 11.6 | 1.0 | 6.3 | 7.5 |
| Quizartinib (AC220) | 5.2 | 7.4 | 6.3 | 1.6 |
| JNJ-7706621 | 10.5 | 2.1 | 6.3 | 5.9 |
| Tyrphostin AG 1296 | 11.1 | 1.5 | 6.3 | 6.7 |
| KN-93 Phosphate | 13.4 | -0.9 | 6.3 | 10.1 |
| NVP-AEW541 | 8.9 | 3.6 | 6.2 | 3.7 |
| SGX-523 | 6.0 | 6.3 | 6.2 | 0.2 |
| Everolimus (RAD001) | 1.3 | 11.0 | 6.2 | 6.9 |
| PD0325901 | 9.7 | 2.6 | 6.2 | 5.0 |
| Palbociclib (PD-0332991) HCl | 7.6 | 4.5 | 6.1 | 2.2 |
| CYT387 | 0.9 | 11.1 | 6.0 | 7.2 |
| GDC-0349 | 14.4 | -2.5 | 5.9 | 11.9 |
| Temsirolimus (CCI-779) | 3.5 | 8.4 | 5.9 | 3.4 |
| TCS 359 | 1.0 | 10.8 | 5.9 | 6.9 |
| Sorafenib | 4.0 | 7.7 | 5.9 | 2.6 |
| Rigosertib (ON-01910) | 12.5 | -0.9 | 5.8 | 9.5 |
| GSK1904529A | -3.3 | 14.9 | 5.8 | 12.9 |
| PP1 | 9.3 | 2.3 | 5.8 | 4.9 |
| MLN8054 | -10.5 | 22.0 | 5.8 | 23.0 |
| AMG-458 | 12.1 | -0.5 | 5.8 | 8.9 |
| P276-00 | 9.2 | 2.3 | 5.7 | 4.9 |
| TAK-733 | 8.3 | 3.1 | 5.7 | 3.7 |
| KU-0063794 | 4.1 | 7.3 | 5.7 | 2.2 |
| BIO | 2.8 | 8.5 | 5.7 | 4.0 |
| 6H05 | -1.3 | 12.5 | 5.6 | 9.7 |
| Hesperadin | 13.3 | -2.2 | 5.5 | 11.0 |
| CA75 (GAK) | 5.7 | 5.3 | 5.5 | 0.3 |
| CCT129202 | -0.8 | 11.7 | 5.4 | 8.8 |
| BMS-265246 | 5.5 | 5.3 | 5.4 | 0.1 |
| SMI-4a | 13.0 | -2.2 | 5.4 | 10.8 |
| Ridaforolimus (Deforolimus) | 6.7 | 3.9 | 5.3 | 2.0 |

| | | | | |
|---|---|---|---|---|
| SC-514 | 10.3 | 0.1 | 5.2 | 7.2 |
| BGT226 (NVP-BGT226) | -1.1 | 11.5 | 5.2 | 8.9 |
| Chrysophanic Acid | 3.4 | 6.8 | 5.1 | 2.4 |
| ZM 323881 HCl | 5.7 | 4.4 | 5.1 | 0.9 |
| PI-103 | 8.6 | 1.5 | 5.0 | 5.0 |
| SL-327 | 12.8 | -2.7 | 5.0 | 11.0 |
| Tie2 kinase inhibitor | -0.1 | 10.0 | 5.0 | 7.2 |
| A66 | 11.1 | -1.4 | 4.8 | 8.9 |
| U0126-EtOH | -4.6 | 14.1 | 4.8 | 13.2 |
| PIK-293 | 3.0 | 6.6 | 4.8 | 2.5 |
| AZD2858 | 6.3 | 3.2 | 4.7 | 2.2 |
| AZD1080 | 5.0 | 4.0 | 4.5 | 0.7 |
| NVP-BVU972 | 8.6 | 0.3 | 4.4 | 5.9 |
| Masitinib (AB1010) | 7.7 | 1.1 | 4.4 | 4.7 |
| Cabozantinib (XL184) | 2.7 | 5.8 | 4.3 | 2.2 |
| VS-5584 (SB2343) | 4.3 | 3.9 | 4.1 | 0.3 |
| WZ4002 | 1.3 | 6.7 | 4.0 | 3.8 |
| GDC-0980 (RG7422) | 4.7 | 3.2 | 4.0 | 1.1 |
| Dovitinib (TKI-258) | -1.3 | 9.0 | 3.9 | 7.3 |
| PF-3758309 | 7.6 | 0.2 | 3.9 | 5.2 |
| Cabozantinib malate (XL184) | 4.4 | 3.1 | 3.8 | 0.9 |
| LY2835219 | 6.8 | 0.3 | 3.6 | 4.6 |
| PF-4708671 | -3.0 | 10.1 | 3.5 | 9.2 |
| Dinaciclib (SCH727965) | 2.0 | 5.0 | 3.5 | 2.1 |
| GDC-0941 | 1.6 | 5.5 | 3.5 | 2.7 |
| PF-562271 | 4.3 | 2.7 | 3.5 | 1.2 |
| Roscovitine (Seliciclib) | 10.7 | -3.8 | 3.4 | 10.3 |
| TGX-221 | 0.4 | 6.3 | 3.3 | 4.2 |
| SB202190 (FHPI) | 3.4 | 3.1 | 3.3 | 0.2 |
| WYE-354 | 12.5 | -6.0 | 3.2 | 13.0 |
| KX2-391 | -4.5 | 10.9 | 3.2 | 10.9 |
| XL019 | -1.4 | 7.7 | 3.2 | 6.5 |
| VX-745 | -12.9 | 19.2 | 3.1 | 22.7 |
| AS-604850 | 5.9 | 0.2 | 3.0 | 4.1 |
| BYL719 | 2.8 | 2.8 | 2.8 | 0.0 |
| PD168393 | 10.7 | -5.2 | 2.7 | 11.3 |
| XMD8-92 | -1.6 | 6.9 | 2.7 | 6.0 |
| VE-822 | 8.4 | -3.5 | 2.4 | 8.5 |
| Fingolimod (FTY720) HCl | 10.7 | -5.9 | 2.4 | 11.7 |
| LY294002 | 11.2 | -6.5 | 2.4 | 12.5 |
| Refametinib (RDEA119) | 7.6 | -2.9 | 2.4 | 7.5 |
| A-674563 | 5.3 | -0.7 | 2.3 | 4.3 |
| S-Ruxolitinib (INCB018424) | 8.0 | -3.5 | 2.3 | 8.1 |

| | | | | |
|---|---|---|---|---|
| PIK-294 | -5.2 | 9.7 | 2.3 | 10.5 |
| Golvatinib (E7050) | 0.2 | 4.3 | 2.2 | 2.9 |
| GNE-0877 | -3.0 | 7.4 | 2.2 | 7.4 |
| Honokiol | 2.1 | 2.1 | 2.1 | 0.0 |
| LDC000067 | 6.3 | -2.2 | 2.1 | 6.0 |
| NMS-P937 (NMS1286937) | -4.4 | 8.4 | 2.0 | 9.1 |
| AZD5363 | 3.0 | 0.9 | 2.0 | 1.5 |
| BIX 02189 | 4.7 | -1.0 | 1.8 | 4.1 |
| Degrasyn (WP1130) | -2.3 | 6.0 | 1.8 | 5.9 |
| EHT 1864 | 7.8 | -4.3 | 1.8 | 8.6 |
| AZD8330 | 2.0 | 1.4 | 1.7 | 0.4 |
| ETP-46464 | 0.3 | 3.1 | 1.7 | 2.0 |
| PF-477736 | 20.3 | -17.0 | 1.6 | 26.3 |
| BX-795 | 11.4 | -8.5 | 1.4 | 14.1 |
| JNJ-38877605 | 5.8 | -3.0 | 1.4 | 6.2 |
| Ki8751 | 2.5 | 0.2 | 1.3 | 1.7 |
| CO-1686 (AVL-301) | -9.1 | 11.8 | 1.3 | 14.8 |
| RKI-1447 | -6.8 | 9.4 | 1.3 | 11.5 |
| KRN 633 | -0.2 | 2.8 | 1.3 | 2.1 |
| CGK 733 | 11.5 | -9.0 | 1.2 | 14.5 |
| Imatinib (STI571) | 9.9 | -7.5 | 1.2 | 12.3 |
| Phenformin HCl | 5.7 | -3.3 | 1.2 | 6.4 |
| PHT-427 | -5.2 | 7.5 | 1.2 | 9.0 |
| Tofacitinib (CP-690550) Citrate | 1.7 | 0.6 | 1.2 | 0.8 |
| Palomid 529 (P529) | 14.0 | -11.9 | 1.0 | 18.3 |
| Pimasertib (AS-703026) | 7.5 | -5.4 | 1.0 | 9.1 |
| PHA-767491 | -2.2 | 4.2 | 1.0 | 4.5 |
| Tyrphostin 9 | -9.0 | 10.9 | 0.9 | 14.1 |
| Pelitinib (EKB-569) | 2.7 | -1.0 | 0.8 | 2.6 |
| AG-1478 (Tyrphostin AG-1478) | 5.1 | -3.6 | 0.8 | 6.1 |
| AP26113 | 0.0 | 1.2 | 0.6 | 0.8 |
| Mubritinib (TAK 165) | 1.4 | -0.4 | 0.5 | 1.3 |
| ZCL278 | -0.4 | 1.3 | 0.5 | 1.3 |
| Icotinib | 7.0 | -6.3 | 0.4 | 9.4 |
| TG100713 | -0.5 | 1.2 | 0.3 | 1.2 |
| Enzastaurin (LY317615) | 2.6 | -1.9 | 0.3 | 3.2 |
| GDC-0068 | 6.0 | -5.5 | 0.3 | 8.2 |
| R406 | 6.2 | -5.8 | 0.2 | 8.5 |
| CAL-101 (Idelalisib) | 0.2 | 0.1 | 0.1 | 0.1 |
| OSI-930 | -0.4 | 0.6 | 0.1 | 0.7 |
| IKK-16 (IKK Inhibitor VII) | 10.2 | -10.2 | 0.0 | 14.5 |
| DMSO (MaxBRET) | 0.0 | 0.0 | 0.0 | 0.0 |
| DMSO (Blank) | 0.0 | 0.0 | 0.0 | 0.0 |

| | | | | |
|---|---|---|---|---|
| A-769662 | 1.2 | -1.3 | 0.0 | 1.7 |
| INK 128 (MLN0128) | -0.8 | 0.1 | -0.3 | 0.6 |
| Varlitinib | 2.7 | -4.3 | -0.8 | 4.9 |
| Flavopiridol HCl | 1.6 | -3.3 | -0.8 | 3.4 |
| MK-5108 (VX-689) | -2.8 | 1.0 | -0.9 | 2.7 |
| TIC10 | 4.1 | -6.1 | -1.0 | 7.2 |
| Thiazovivin | -3.6 | 1.5 | -1.1 | 3.6 |
| GNE-9605 | 3.1 | -5.2 | -1.1 | 5.8 |
| IPI-145 (INK1197) | -2.8 | 0.6 | -1.1 | 2.4 |
| BMS-345541 | 7.2 | -9.4 | -1.1 | 11.7 |
| CAY10505 | -12.3 | 10.0 | -1.2 | 15.8 |
| PP2 | -3.1 | 0.5 | -1.3 | 2.5 |
| Skepinone-L | 6.5 | -9.2 | -1.4 | 11.0 |
| AZD2014 | 8.9 | -11.6 | -1.4 | 14.5 |
| AZD1208 | 5.0 | -7.9 | -1.4 | 9.2 |
| PQ 401 | -4.0 | 1.2 | -1.4 | 3.7 |
| OSI-420 | -6.8 | 3.7 | -1.5 | 7.4 |
| AZD4547 | -24.6 | 21.5 | -1.5 | 32.6 |
| Fasudil (HA-1077) HCl | 6.9 | -10.2 | -1.7 | 12.1 |
| 3-Methyladenine | 2.8 | -7.1 | -2.2 | 7.0 |
| GSK2636771 | 2.9 | -7.3 | -2.2 | 7.3 |
| Ro3280 | -2.3 | -2.3 | -2.3 | 0.0 |
| AZD3463 | 8.0 | -12.9 | -2.4 | 14.8 |
| TG003 | 10.1 | -15.2 | -2.5 | 17.9 |
| MK-2461 | -9.3 | 3.8 | -2.7 | 9.3 |
| WHI-P154 | 7.3 | -12.9 | -2.8 | 14.3 |
| GSK1059615 | -4.2 | -1.5 | -2.8 | 1.9 |
| VE-821 | -11.1 | 5.3 | -2.9 | 11.6 |
| TPCA-1 | 1.2 | -7.6 | -3.2 | 6.2 |
| Imatinib Mesylate (STI571) | -2.0 | -4.5 | -3.2 | 1.8 |
| WAY-600 | 4.0 | -10.8 | -3.4 | 10.5 |
| Dabrafenib (GSK2118436) | 0.7 | -7.9 | -3.6 | 6.1 |
| Wortmannin | -2.4 | -4.8 | -3.6 | 1.7 |
| GNE-7915 | -10.5 | 3.1 | -3.7 | 9.6 |
| Erlotinib HCl (OSI-744) | -0.1 | -7.3 | -3.7 | 5.1 |
| CEP-32496 | -2.2 | -5.2 | -3.7 | 2.1 |
| UNC-AA-1-0013 (AAK1) | 3.7 | -11.6 | -4.0 | 10.8 |
| SB216763 | 0.9 | -9.6 | -4.4 | 7.4 |
| Asiatic Acid | 8.6 | -17.4 | -4.4 | 18.4 |
| Regorafenib (BAY 73-4506) | -4.0 | -5.3 | -4.6 | 1.0 |
| XL147 | -8.3 | -1.7 | -5.0 | 4.6 |
| GNF-5 | -7.9 | -2.3 | -5.1 | 3.9 |
| GF109203X | -5.7 | -5.0 | -5.4 | 0.6 |

| | | | | |
|---|---|---|---|---|
| ZM 336372 | -4.5 | -6.5 | -5.5 | 1.4 |
| TAK-901 | -2.7 | -9.1 | -5.9 | 4.5 |
| Dacomitinib (PF299804) | 1.4 | -14.2 | -6.4 | 11.0 |
| BIX 02188 | 4.1 | -17.0 | -6.5 | 14.9 |
| Nintedanib (BIBF 1120) | -1.8 | -11.3 | -6.6 | 6.7 |
| SU11274 | -7.3 | -6.0 | -6.7 | 0.9 |
| GW5074 | -7.5 | -6.2 | -6.8 | 0.9 |
| CNX-774 | -4.4 | -9.3 | -6.8 | 3.5 |
| SB415286 | 2.4 | -16.5 | -7.0 | 13.4 |
| IM-12 | -0.3 | -14.5 | -7.4 | 10.0 |
| WP1066 | 3.4 | -18.3 | -7.5 | 15.3 |
| PP121 | -0.4 | -14.8 | -7.6 | 10.2 |
| Fostamatinib (R788) | 0.9 | -17.9 | -8.5 | 13.3 |
| Apatinib | -9.2 | -7.8 | -8.5 | 1.0 |
| PD98059 | -16.6 | -0.7 | -8.6 | 11.3 |
| GSK650394 | -24.3 | 6.7 | -8.8 | 21.9 |
| OSI-027 | 5.5 | -23.2 | -8.9 | 20.3 |
| KU-55933 (ATM Kinase Inhibitor) | 4.0 | -21.9 | -9.0 | 18.3 |
| AST-1306 | -7.7 | -10.7 | -9.2 | 2.1 |
| ZM 39923 HCl | 0.2 | -18.8 | -9.3 | 13.5 |
| PFK15 | -10.7 | -9.2 | -10.0 | 1.0 |
| Butein | 3.3 | -25.0 | -10.8 | 20.0 |
| PD184352 (CI-1040) | -2.3 | -19.9 | -11.1 | 12.5 |
| GNF-2 | -8.5 | -14.2 | -11.3 | 4.1 |
| LY2603618 | 7.0 | -30.5 | -11.8 | 26.5 |
| Go 6983 | -11.0 | -14.1 | -12.6 | 2.3 |
| CX-6258 HCl | -0.5 | -26.8 | -13.6 | 18.5 |
| TAE226 (NVP-TAE226) | -26.7 | -0.7 | -13.7 | 18.4 |
| AT7519 | 6.4 | -34.0 | -13.8 | 28.6 |
| Bosutinib (SKI-606) | 7.7 | -36.0 | -14.2 | 30.9 |
| Dasatinib | 1.5 | -30.7 | -14.6 | 22.8 |
| PF-04691502 | -3.4 | -25.8 | -14.6 | 15.8 |
| Filgotinib (GLPG0634) | -9.1 | -21.6 | -15.3 | 8.8 |
| K-Ras(G12C) inhibitor 9 | -21.1 | -10.0 | -15.5 | 7.9 |
| BAY 11-7082 | -17.1 | -14.7 | -15.9 | 1.7 |
| AG-490 (Tyrphostin B42) | -25.0 | -10.6 | -17.8 | 10.2 |
| IMD 0354 | -39.3 | -10.2 | -24.8 | 20.6 |
| Semaxanib (SU5416) | -36.5 | -17.9 | -27.2 | 13.1 |
| Sotrastaurin | -23.3 | -37.6 | -30.5 | 10.1 |
| Ro 31-8220 Mesylate | -39.9 | -26.9 | -33.4 | 9.2 |
| SSR128129E | -35.8 | -32.8 | -34.3 | 2.1 |
| TSU-68 (SU6668) | -51.3 | -43.8 | -47.5 | 5.3 |
| PHA-665752 | -60.9 | -54.3 | -57.6 | 4.7 |

| | | | | |
|---|---|---|---|---|
| Sunitinib Malate | -94.9 | -51.0 | -73.0 | 31.0 |
| AR-A014418 | -89.7 | -72.9 | -81.3 | 11.9 |
| AG-1024 | -120.2 | -86.4 | -103.3 | 23.9 |
| AG-18 | -78.7 | -139.9 | -109.3 | 43.3 |
| CHIR-99021 (CT99021) HCl | -257.5 | -376.8 | -317.1 | 84.4 |

#### Supplementary Figure S1. Sequence alignment and deep learning-based structure prediction for LmxCLK1 and LmxCLK2.

**A**

```
LmxCLK2      MAVAAVTGSSHGEEATGRRSGSKRDHEASIT--------------TDQN-PQDAAKALPPPKKKKVTYTLPHQNMEEGHFY 66
LmxCLK1      ---------MSRSQSDARRSGSKRTFDEATATDNVQQRRHDGTAVPNTAPNQVVEVVAPPPKKKKVTYALPHQNMEEGHFY 72
                              *******                           *      ********** ************

LmxCLK2      VVLGEDIDVSTQRFKILSLLGEGTFGKVVESWDRKRKEYCAVKIVRNVPKYTRDAKIEIQFMEKVRQADPADRFPLMKIQR 147
LmxCLK1      VVLGEDIDVSTQRFKILSLLGEGTFGKVVESWDRKRKEYCAVKIVRNVPKYTRDAKIEIQFMEKVRQADPADRFPLMKIQR 153
             *********************************************************************************

LmxCLK2      YFQNDSGHMCIVMPKYGPCLLDWIMKHGPFNHRHLAQIVFQTGVALDYFHSELHLMHTDLKPENILMETSDTTVDPATNRH 228
LmxCLK1      YFQNDSGHMCIVMPKYGPCLLDWIMKHGPFNHRHLAQIVFQTGVALDYFHSELHLMHTDLKPENILMETSDTTVDPATNRH 234
             *********************************************************************************

LmxCLK2      LPPDPCRVRICDLGGCCDERHSRTAIVSTRHYRSPEVILGLGWMYSTDMWSMGCIIYELYTGKLLYDTHDNLEHLHLMEKT 309
LmxCLK1      LPPDPCRVRICDLGGCCDERHSRTAIVSTRHYRSPEVILGLGWMYSTDMWSMGCIIYELYTGKLLYDTHDNLEHLHLMEKT 305
             *********************************************************************************

LmxCLK2      LGRLPSEWAARCGTEEARLLYNSAGQLRPCTDPKHLARIARARTVRDVIRDDLLCDLIYGLLHYDRQKRLNARQMTTHPYV 390
LmxCLK1      LGRLPSEWAARCGTEEARLLYNSAGQLRPCTDPKHLARIARARTVRDVIRDDLLCDLIYGLLHYDRQKRLNARQMTTHPYV 396
             *********************************************************************************

LmxCLK2      LKYYPEASQAPSYPDNRPMLRPPPIM 416
LmxCLK1      LKYYPEASQAPSYPDNRPMLRPPPIM 422
             **************************
```

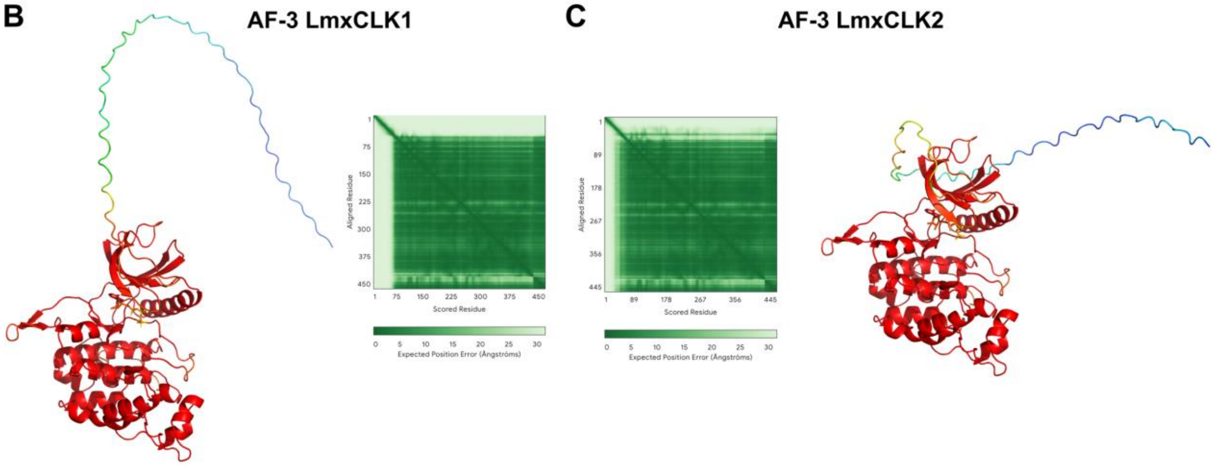

**Supplementary Figure S1.** Sequence alignment and deep-learning based structure prediction for LmxCLK1 and LmxCLK2. **A**: Sequence alignment for LmxCLK1 (TritrypDB ID LmxM.09.0400, UniProt ID: E9AMP2) and LmxCLK2 (TritrypDB ID LmxM.09.0410, UniProt ID: E8NHK0). Stars indicate identical residues; amino acid sequence for the kinase domain is shown in bold font. **B,C**: Cartoon representation and quality assessment of the AlphaFold3-predicted structure using the amino acid sequences for LmxCLK1 (**B**) and LmxCLK2 (**C**). The color scheme for the cartoon representation summarizes the per-residue confidence scores (pLDDT), ranging from 0 (very low confidence or likely disordered regions; blue) to 100 (very high confidence; red). The middle panels show Predicted Aligned Error (PAE) heatmaps indicating the expected positional error (in Å) between residue pairs when aligned on the true structure. Lower values (dark green) indicate high confidence in the relative positioning of residues, while higher values (dark to light green) reflect increased uncertainty, often associated with inter-domain flexibility or disorder.

#### Supplementary Figure S2. Protein purification of full-length LmxCLK1 (cb001) and NLuc-LmxCLK1 (cb021).

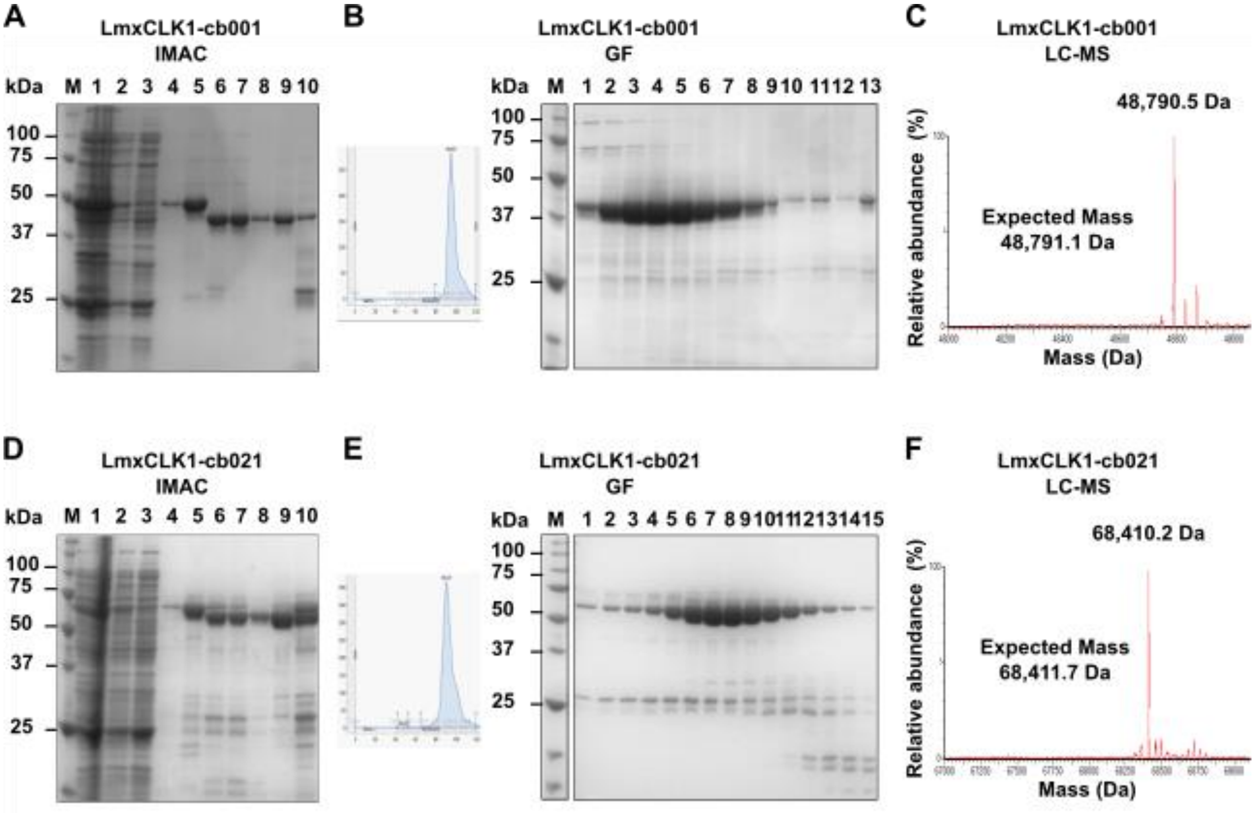

**Supplementary Figure S2.** Purification of LmxCLK1 (cb001) (top panels) and NLuc-LmxCLK1 (cb021) (bottom panels) expressed in BL21(DE3)-R3-lambda-PPase *E. coli* strain. **A, D:** SDS-PAGE analysis of IMAC fractions: total lysate (1), supernatant (2), flow-through (3), wash in 30 mM imidazole (4) and eluate in 300 mM imidazole (5). After TEV protease treatment (6), samples were further purified using reverse affinity chromatography on Ni-Sepharose and the following fractions were analyzed: flow-through (7), wash in 30 mM imidazole (8), wash in 60 mM imidazole (9) and eluate in 300 mM imidazole (10). **B, E:** Gel filtration (GF) chromatograms of TEV-treated proteins from step A, D (fractions 7 and 8 – flow-through and wash in 30 mM imidazole, respectively), and SDS-PAGE analysis of GF fractions. M: Precision Plus Protein Unstained Protein Standards (Bio-Rad). **C, F:** Liquid chromatography-mass spectrometry (LC-MS) analysis of purified proteins. Deconvoluted mass/charge spectra are shown. Expected and observed mass values are indicated.

#### Supplementary Figure S3. Enzymatic Assay development.

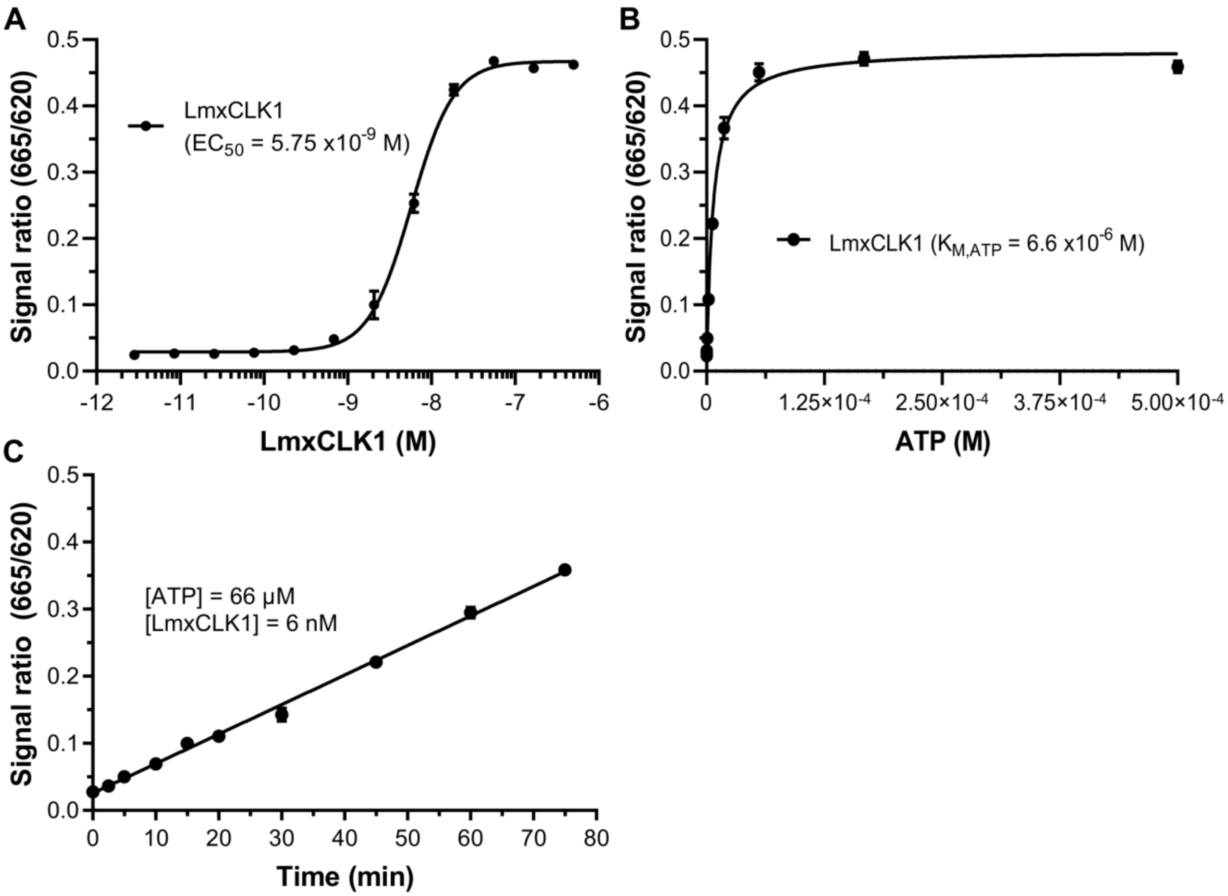

**Supplementary Figure S3.** Results obtained during LmxCLK1 (cb001) enzymatic assay development. **A:** Establishing enzyme working concentration. Concentration-dependent phosphorylation of peptide substrate by LmxCLK1 as indicated by increasing fluorescent signal ratio. $EC_{50}$ value shown was calculated by fitting the experimental data to a sigmoidal dose-response (variable slope) model (four-parameters). **B:** $K_{M,ATP}$ determination. LmxCLK1 (15 nM final concentration) activity as a function of ATP concentration. $K_{M,ATP}$ value shown was obtained by fitting the experimental data to the Michaelis-Menten model. **C:** Time-course of peptide phosphorylation by LmxCLK1. Enzyme and ATP concentrations used are indicated. Line shown represents linear regression ($r^2$ = 0.9934) of experimental points. In panels A-C data shown are mean ± SEM of two independent measurements.

#### Supplementary Figure S4. Mass spectrometry analysis confirms the formation of covalent adducts between LmxCLK1 and compounds WZ8040 and WZ3146.

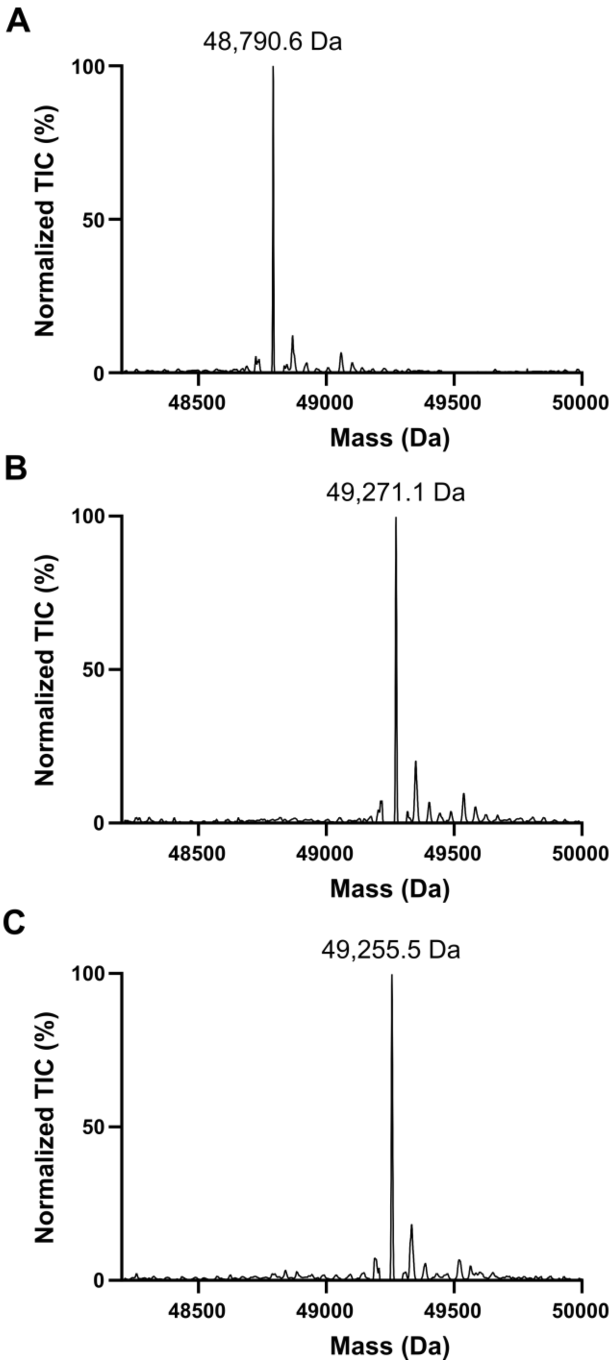

**Supplementary Figure S4.** Mass spectrometry analysis confirms the formation of covalent adducts between LmxCLK1 and compounds WZ8040 and WZ3146. Intact mass determination of purified LmxCLK1 (cb001) following incubation with: buffer **(A)**, WZ8040 **(B)** or WZ3146 **(C)** for 2 hours at room temperature. The expected intact mass for recombinant LmxCLK1 is 48,791.1 Da. Adduct formation was observed for both compounds, resulting in mass shifts consistent with the addition of one molecule of WZ8040 (+481.01 Da) or WZ3146 (+464.95 Da). Deconvoluted mass/charge spectra are shown. TIC: total ion count.

### Supplementary Figure S5. Sequence coverage and WZ8040 covalent adducts identified by LC-MS/MS following tryptic digest of full-length LmxCLK1.

| | | | | | |
|---|---|---|---|---|---|
| **1** | SMSRSQSDAR | RSGSKRTFDE | ATATDNVQQR | RHDGTAVPNT | APNQVVEVVA |
| **51** | PPPKKKKVTY | ALPHQNMEEG | HFYVVLGEDI | DVSTQRFKIL | SLLGEGTFGK |
| **101** | VVESWDRKRK | EYCAVKIVRN | VPKYTRDAKI | EIQFMEKVRQ | ADPADRFPLM |
| **151** | KIQRYFQNDS | GHMCIVMPK**Y** | **GPCLLDWIM** | HGPFNHRHLA | QIVFQTGVAL |
| **201** | DYFHSELHLM | HTDLKPENIL | METSDTTVDP | ATNR**HLPPDP** | **CR**VR**ICDLGG** |
| **251** | **CCDER**HSRTA | IVSTRHYRSP | EVILGLGWMY | STDMWSMGCI | IYELYTGKLL |
| **301** | YDTHDNLEHL | HLMEKTLGRL | PSEWAAR**CGT** | **EEAR**LLYNSA | GQLRPCTDPK |
| **351** | HLARIARART | VRDVIRDDLL | CDLIYGLLHY | DRQKRLNARQ | MTTHPYVLKY |
| **401** | YPEASQAPSY | PDNRPMLRPP | PIM | | |

**Supplementary Figure S5.** Sequence coverage and WZ8040 covalent adducts identified by LC-MS/MS following tryptic digest of full-length LmxCLK1. Overall sequence coverage was approximately 92% (residues in gray were not identified). Modified peptides are underlined in bold and the sites of covalent modification highlighted in red. Recombinant full-length LmxCLK1 has an additional N-terminal serine introduced by our cloning strategy and left over after cleavage of N-terminal His-tag with TEV protease.

#### Supplementary Figure S6. Identification of modified peptides by LC-MS/MS.

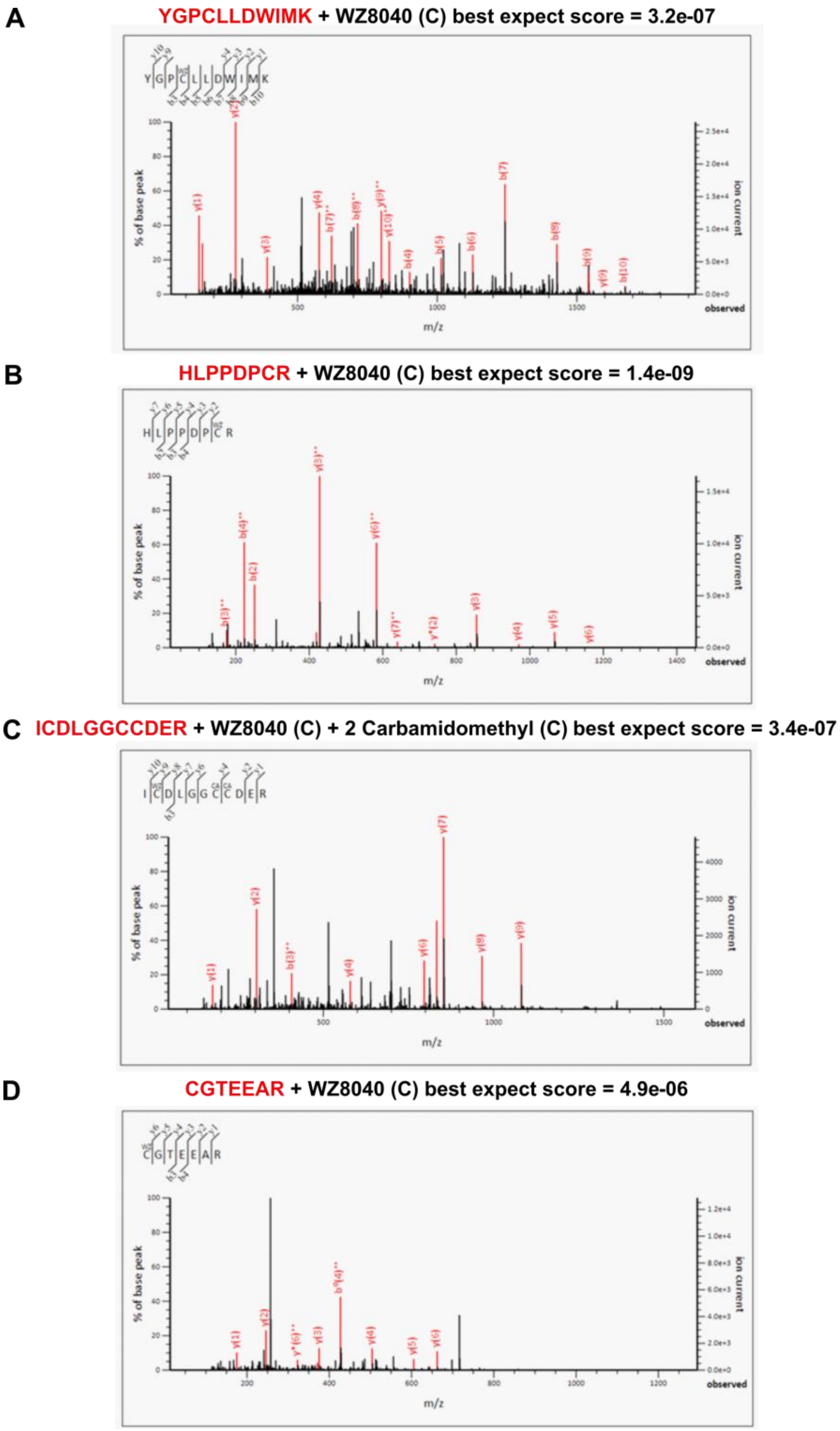

**Supplementary Figure S6.** Identification of modified peptides by LC-MS/MS. Purified LmxCLK1 (CQMED lone ID LmxCLK1-cb001) was incubated with compound WZ8040 in order to determine the probable LmxCLK1 cysteine interacting with WZ8040. Fragmentation spectra of four modified peptides obtained from the tryptic digest are shown: YGPCLLDWIMK **(A)**, HLPPDPCR **(B)**, ICDLGGCCDER **(C)**, and CGTEEAR **(D)**.

Supplementary Figure S7. Determination of the probable LmxCLK1 cysteine interacting with WZ8040.

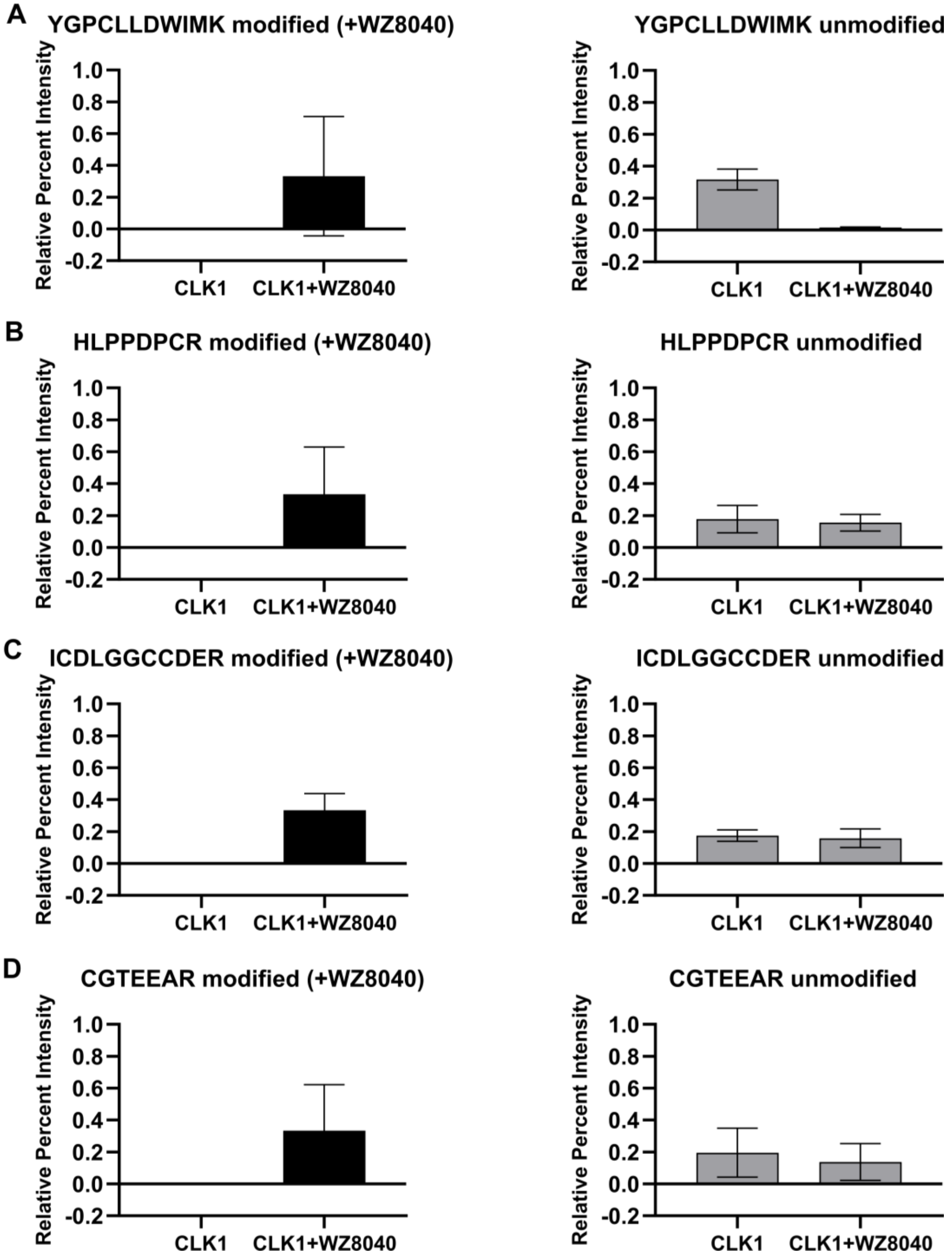

**Supplementary Figure S7.** Identification of LmxCLK1 cysteine residues modified by WZ8040. Bar graphs show the relative percent intensity of modified versus unmodified peptides in untreated and treated samples. The data indicate that Cys172 (peptide: YGPCLLDWIMK) is the predominant site of covalent modification. Additional, lower-frequency adducts were observed at Cys240 (HLPPDPCR), Cys245 (ICDLGGCCDER), and Cys327 (CGTEEAR).

#### Supplementary Figure S8. AlphaFold model.

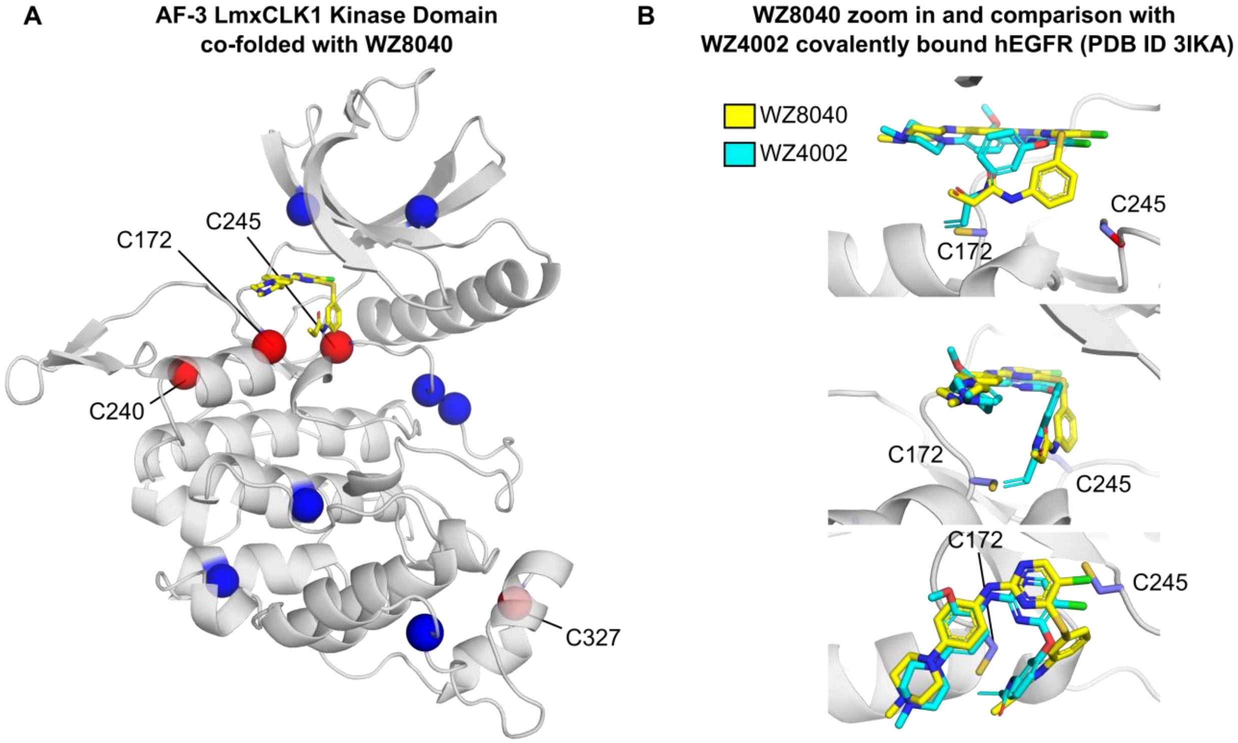

**Supplementary Figure S8.** Location of cysteine residues in LmxCLK1 and predicted binding mode of WZ8040. (A) AlphaFold-3–predicted structure of the LmxCLK1 kinase domain (grey, semi-transparent cartoon), co-folded in the presence of WZ8040 (shown as sticks). Cysteine Cα atoms are depicted as spheres; red spheres indicate residues identified by LC-MS/MS as susceptible to covalent adduct formation following incubation with WZ8040. (B) Predicted binding mode of WZ8040 to the LmxCLK1 kinase domain (carbon atoms in yellow), overlaid with the covalently bound pose of the close analog WZ4002 (carbon atoms in cyan) in human EGFR (PDB ID: 3IKA).

### References

(1) Savitsky, P.; Bray, J.; Cooper, C. D. O.; Marsden, B. D.; Mahajan, P.; Burgess-Brown, N. A.; Gileadi, O. High-Throughput Production of Human Proteins for Crystallization: The SGC Experience. *J. Struct. Biol.* **2010**, *172* (1), 3–13. https://doi.org/10.1016/J.JSB.2010.06.008.

(2) Tetaud, E.; Lecuix, I.; Sheldrake, T.; Baltz, T.; Fairlamb, A. H. A New Expression Vector for Crithidia Fasciculata and Leishmania. *Mol. Biochem. Parasitol.* **2002**, *120* (2), 195–204. https://doi.org/10.1016/S0166-6851(02)00002-6.

(3) Beneke, T.; Madden, R.; Makin, L.; Valli, J.; Sunter, J.; Gluenz, E. A CRISPR Cas9 High-Throughput Genome Editing Toolkit for Kinetoplastids. *R. Soc. open Sci.* **2017**, *4* (5), 1–16. https://doi.org/10.1098/RSOS.170095.
